# Differential levels of IFNα subtypes in autoimmunity and viral infection

**DOI:** 10.1101/2021.03.04.433900

**Authors:** Vincent Bondet, Mathieu P Rodero, Céline Posseme, Pierre Bost, Jérémie Decalf, Liis Haljasmägi, Nassima Bekaddour, Gillian Rice, Vinit Upasani, Jean-Philippe Herbeuval, John A Reynolds, Tracy A Briggs, Ian N Bruce, Claudia Mauri, David Isenberg, Madhvi Menon, David Hunt, Benno Schwikowski, Xavier Mariette, Stanislas Pol, Flore Rozenberg, Tineke Cantaert, J Eric Gottenberg, Kai Kisand, Darragh Duffy

**Author notes:** Corresponding author: Darragh Duffy, Translational Immunology Lab, Institut Pasteur 25, rue du Dr. Roux, 75724 Paris Cedex 15, France.

## Abstract

Type I interferons are essential for host response to viral infections, while dysregulation of their response can result in autoinflammation or autoimmunity. Among IFNα (alpha) responses, 13 subtypes exist that signal through the same receptor, but have been reported to have different effector functions. However, the lack of available tools for discriminating these closely related subtypes, in particular at the protein level, has restricted the study of their differential roles in disease. We developed a digital ELISA with specificity and high sensitivity for the IFNα2 subtype. Application of this assay, in parallel with our previously described pan-IFNα assay, allowed us to study different IFNα protein responses following cellular stimulation and in diverse patient cohorts. We observed different ratios of IFNα protein responses between viral infection and autoimmune patients. This analysis also revealed a small percentage of autoimmune patients with high IFNα2 protein measurements but low pan-IFNα measurements. Correlation with an ISG score and functional activity showed that in this small sub group of patients, IFNα2 protein measurements did not reflect its biological activity. This unusual phenotype was partly explained by the presence of anti-IFNα auto-antibodies in a subset of autoimmune patients. This study reports ultrasensitive assays for the study of IFNα proteins in patient samples and highlights the insights that can be obtained from the use of multiple phenotypic readouts in translational and clinical studies.

## Introduction

Type I interferons are essential for host responses to viral infections ^1^. Their dysregulation is also implicated in many autoimmune diseases, such as systemic lupus erythematosus (SLE) and primary Sjögren’s syndrome (pSS) ^2^, while their exact role in bacterial infection remains unclear ^3^. Within the type I interferon class there are 9 types; IFNα (alpha), IFNβ (beta), IFNκ (kappa), IFNω (omega), IFNδ (delta), IFNε (epsilon), IFNτ (tau), IFNζ (zeta) and IFNν (nu) (IFNδ/τ/ζ/ν not found in humans) that all signal through the IFNΑR receptor complex ^4^. Furthermore, within the IFNα types there are 13 subtypes that exhibit marked evolutionary differences in terms of selective conservation, suggesting different immunological relevance ^5^.

Some studies have reported differential functions, at least in terms of anti-viral responses ^6,7,8,9^, but much remains unknown in terms of broader differential activity between the different IFNα subtypes. Furthermore, in autoimmunity, the potential role of different IFNα subtypes is even less clear, despite their strong implication in the pathogenesis of many such diseases ^10^. Such studies have been restricted by the high level of sequence homology among the different subtypes, as well as the lack of tools to study them, in particular at the protein level. Indeed, until recently, IFNα protein was challenging to directly measure in human samples, with interferon stimulated genes (ISG) or functional activity used as proxy readouts. This limitation was overcome with the application of Simoa digital ELISA ^11^, and the use of unique monoclonal antibodies isolated from APECED patients ^12^ that enabled the detection of all IFNα protein subtypes with more or less equivalent and high sensitivity ^13^. Direct measurement of the protein allowed us to identify the cellular sources of pathological IFNα in patients with STING mutations and juvenile dermatomyositis, to be circulating monocytes and pDCs ^13^ and tissue myoblasts ^14^, respectively. In contrast in active TB infection, we demonstrated an absence of plasma IFN-I ^15^, suggesting that the widely reported blood ISG signature most likely reflects signaling in the infected tissue ^3^.

In this study we adapted the digital ELISA using commercial antibodies specific for IFNα2 and used both assays to study autoimmune and infected patient cohorts. While both assays correlated well in the majority of patients, discrepancies between the two were observed which revealed interesting clinical observations. In particular, the balance between the results obtained using the two assays was modified by the presence of autoantibodies in some patients and the nature of the TLR that was activated. This provides a starting point for beginning to dissect potential different roles of IFNα subtypes in human disease.

## Materials and Methods

### Patient cohorts

Patients were not specifically enrolled for this study, all quantifications were done on samples from existing clinical cohorts that were previously described: Rodero *et al*.^13^, Reynolds *et al.* (LEAP cohort) ^16^, Bost *et al.* (In review) (ASSESS cohort), Menon *et al*. (In preparation), Sultanik *et al.*^*117*^ (C10-08 cohort) and Upasani *et al.*^18^.

The Rodero *et al*. (2017) cohort includes patients with systemic lupus erythematosus (SLE), juvenile SLE (JSLE), connective tissue disease (CTD), retinal vasculopathy with cerebral leukodystrophy (RVCL), juvenile dermatomyositis (JDM), monogenic interferonopathies (MI), and central nervous system (CNS) infections. All samples were collected with informed consent. The study was approved by the Leeds (East) Research Ethics Committee (reference number 10/H1307/132), by the Comité de Protection des Personnes (ID-RCB/EUD RACT:2014-A01017-40), and by the South-East Scotland Research Ethics Committee (0114/SS/0003).

The Lupus Extended Autoimmune Phenotype (LEAP) cohort includes patients with systemic lupus erythematosus (SLE), primary Sjögren’s syndrome (pSS), undifferentiated connective tissue disease (UCTD), mixed connective tissue disease (MCTD), systemic sclerosis (SSc) and idiopathic inflammatory myopathies (IIM). Patients were recruited from Manchester University NHS Foundation Trust and Salford Royal NHS Foundation Trust. Ethical approval was obtained from the Greater Manchester East Research Ethics Committee (13/NW/0564).

The ASSESS national multi-center prospective cohort (Assessment of Systemic complications and Evolution in Sjögren’s Syndrome) was set up in 2006 thanks to a grant from the French Ministry of Health. Fifteen centers for autoimmune disease were established to recruit consecutive patients with primary Sjögren’s syndrome fulfilling American-European Consensus Criteria (AECG) between 2006 and 2009. The study was approved by the Ethics Committee of Bichat Hospital in 2006. All patients gave their informed written consent. This study was followed for 5 years with the grant of the French Ministry of Health and this study will be extended for 20 years by the French Society of Rheumatology (SFR).

The Menon *et al*. (in preparation) cohort consists of serum samples from 476 SLE patients (mean age 46 ± 15 years, 40 male) recruited at the University College of London. 11 patients with high autoantibody values were studied longitudinally. The ethics committee of the University College London Hospitals National Health Service Trust approved this study. Patients were recruited after obtaining informed consent.

The C10-08 cHCV cohort (n=88) were sampled as part of a prospective study (C10-08) sponsored by the Institut National de la Sante et de la Recherche Medicale (Inserm) and by the Agence Nationale de Recherche sur le Sida et les Hepatites Virales (ANRS-C10-08). Approval of the study was obtained from the French Comite de protection des personnes (CPP IDF II) in 2010, August 2nd, and all patients gave written informed consent. This cohort includes patients with chronic hepatitis C virus infection (genotype is 1 and 4) prior to treatment with either pegylated-IFNα and ribavirin or a direct acting anti-viral (boceprevir or telaprevir).

The Upasani *et al.*^18^ cohort includes 115 children (≥ 2 years) who presented with dengue-like symptoms at the Kanta Bopha Hospital in Phnom Penh, Cambodia. Blood samples were obtained within 96 h of fever onset at hospital admittance. Plasmas were tested for presence of dengue virus using a nested RT-qPCR at the Institut Pasteur du Cambodge. Patients were classified according to the WHO 1997 severity criteria upon hospital discharge as dengue fever (DF), dengue haemorrhagic fever (DHF) and dengue shock syndrome (DSS). Ethical approval for the study was obtained from the National Ethics Committee of Health Research of Cambodia. Patients were recruited after obtaining written informed consent. The dengue cohort of this paper includes 56 of these 115 patients, randomly selected. The dengue cohort and the Upasani *et al.*^18^ cohort are not statistically different in terms of age (10, IQR 6-12 vs 9, IQR 5-11) and pan-alpha interferon concentrations (p=0.536, Mann-Whitney test), and these two cohorts show similar percentages of female (48% vs 53), DF (82% vs 72), DHF (13% vs 17), DSS (5% vs 9), primary infection (25% vs 17) or secondary infection (75% vs 71) patients.

The viral CNS infection cohort (n=18) includes all Rodero *et al*.^13^ related patients whose diagnosis was mainly viral meningitis or viral encephalitis. Details are shown on Table S1.

For this study, samples were categorized based on clinical diagnosis and are summarized in Table 1. All SLE and JSLE samples from Rodero *et al.*^13^ (n=56) and LEAP cohorts (n=39) were combined here to form the SLE cohort. All anti-IFNα positive and a randomly selected group of anti-IFNα negative SLE patients with a similar size (n=111) were added from Menon *et al.* (in preparation) cohort to this SLE cohort (n=206). All CTD samples from Rodero *et al.*^13^ cohort (n=19) and UCTD and MCTD samples from LEAP cohort (n=33) were merged here to form the CTD cohort (n=52). All pSS samples from LEAP cohort (n=12) and all samples from ASSESS cohort (n=380) were merged to form the pSS cohort (n=392). Longitudinal SLE samples (n=11) from Menon *et al.* (in preparation) study were studied in a different part. Patient numbers, gender and age are shown in Table 1 for each cohort.

**Table 1.** Demographic characteristics of each cohort. Patient number, gender, age and diagnosis for each cohort and diagnosis. Data are shown as the n (%) or median (IRQ). LEAP (lupus extended autoimmune phenotype) cohort: see Reynolds et al.^16^. Rodero et al. cohort: see reference 13. ASSESS (assessment of systemic complications (signs) and evolution in Sjögren’s syndrome) cohort: see Bost et al. (In review). Menon et al. cohort: paper in preparation. C10-08 cohort: see Sultanik et al.^17^. Upasani et al. cohort: see reference 18. MCTD: mixed CTD. UCDT: undifferentiated CTD. pSS: primary Sjögren’s syndrome. SLE: systemic lupus erythematosus. CTD: connective tissue disease. JSLE : juvenile SLE. CNS: central nervous system. cHCV: chronic hepatitis C virus infection. DF: dengue fever. DHF: dengue haemorrhagic fever. DSS: dengue shock syndrome.

| Cohort | Patient number | Gender, female | Age | Diagnosis |
| --- | --- | --- | --- | --- |
| LEAP | 84 | 79 (94%) | 49 (36-57) | MCTD: 6 (7%)<br>UCTD: 27 (32%)<br>pSS: 12 (14%)<br>SLE: 39 (46%) |
| Rodero *et al.* (2017) | 93 | 76 (82%) | 42 (26-54) | CTD: 19 (20%)<br>JSLE: 6 (6%)<br>SLE: 50 (54%)<br>Viral CNS infection: 18 (19%) |
| ASSESS | 380 | 353 (93%) | 59 (51-67) | pSS: 380 (100%) |
| Menon *et al.* (date) | 111 | 100 (90%) | 53 (40-63) | SLE: 111 (100%) |
| C10-08 | 88 | 41 (47%) | 58 (50-65) | cHCV: 88 (100%) |
| Upasani *et al.* (2020) | 56 | 27 (48%) | 10 (6-12) | DF: 46 (82%)<br>DHF: 7 (13%)<br>DSS: 3 (5%) |

**Table 1. Demographic characteristics of each cohort.**
| Cohort | Patient number | Gender, female | Age |
| --- | --- | --- | --- |
| CTD | 52 | 48 (92%) | 48 (33-53) |
| pSS | 392 | 365 (93%) | 58 (51-67) |
| SLE | 206 | 189 (92%) | 50 (38-60) |
| cHCV | 88 | 41 (47%) | 58 (50-65) |
| Dengue | 56 | 27 (48%) | 10 (6-12) |
| Viral CNS infection | 18 | 6 (33%) | 2 (<1-31) |

### Healthy donors

Fresh whole blood was collected into sodium heparin tubes from healthy French volunteers enrolled at the Clinical Investigation and Access to BioResources (ICAReB) platform (Center for Translational Research, Institut Pasteur, Paris, France). These donors were part of the CoSImmGEn cohort (NCT03925272). The biobank activity of this platform is certified ISO 9001 and NFS 96-900. Written informed consent was obtained from all study participants. Fresh whole blood was diluted 1:3 with RPMI medium 1640 (1X) + GlutaMAX into 5ml Polystyrene round-bottom tube. Diluted whole blood was either left without any stimulant to mimic the Null condition or stimulated with LPS (LPS-EB ultrapure InvivoGen) at the final concentration of 10ng/ml, Poly(I:C) (HMW VacciGrade InvivoGen) at the final concentration of 20μg/ml, R848 (Vaccigrade InvivoGen) at the final concentration of 1uM. Tubes were vortexed and incubated at 37°C, 5% CO2 for 22 hours. Supernatants were collected and stored at −20°C until use.

### pDC isolation and stimulation

Blood from healthy donors was obtained from “Etablissement Français du Sang” (convention # 07/CABANEL/106; Paris, France). Experimental procedures with human blood were done according to the European Union guidelines and the Declaration of Helsinki. Human peripheral blood mononuclear cells (PBMCs) were isolated by density centrifugation from peripheral blood leukocyte separation medium (STEMCELL Technologies). *In vitro* experiments were performed using human plasmacytoid dendritic cells (pDCs) purified from PBMCs by negative selection with the EasySep Human Plasmacytoid DC enrichment kit (STEMCELL Technologies). pDCs were cultured in RPMI 1640 (Invitrogen, Gaithersburg, MD) containing 10% heat-inactivated fetal bovine serum and 1mM glutamine (Hyclone, Logan, UT). For stimulation experiments purified pDCs were seeded at 1×10^5^/100 μl. Cells were stimulated for 16 hours with the TLR7/TLR8 agonist Resiquimod (R848; Invivogen) at 5 μg/mL or the TLR4 agonist lipopolysaccharide (LPS; Invivogen) at 100 ng/mL or the TLR9 agonist class A CpG ODNs (CpG-A; Invivogen) at 5 μM or the STING agonist cGAMP (Invivogen) or the TLR3 agonist polyinosinic-polycytidylic acid (poly(I:C); Invivogen) at 5 ug/mL. Supernatants were then collected for IFNα detection.

### Sample preparation for digital-ELISA assays

All serum and plasma samples were thawed and centrifugated at 10,000g, +4°C for 10 minutes. Supernatants were diluted in the Detector / Sample Diluent (Quanterix) for pan-alpha or IFNα2 quantification, then incubated one hour at room temperature before analysis. Biological samples were diluted from 1/3 to 1/300 depending on the amount of material available and to avoid saturation. NP40 was added in the Detector / Sample Diluent used for cHCV, dengue or viral CNS infection samples at a final concentration of 0.5% (v/v) to inactivate viruses.

### Calibrators and pure recombinant IFNα subtypes

Recombinant IFNα1, IFNα1(Val114), IFNα4a, IFNα4b, IFNα5, IFNα6, IFNα7, IFNα8, IFNα10, IFNα14, IFNα16, IFNα17, and IFNα21 were purchased from PBL Assay Science. Recombinant IFNα2c was purchased from eBioscience. Recombinant IFNβ, IFNλ1, IFNλ2, IFNω and IFNγ were purchased from PeproTech.

### Pan-IFNα digital ELISA assay

The Simoa pan-IFNα assay was developed using the Quanterix Homebrew kit according to the manufacturer’s instructions and using two autoantibodies specific for IFNα isolated and cloned from two APS1/APECED patients. The 8H1 antibody clone was used as a capture antibody after coating on paramagnetic beads (0.3mg/mL), and the 12H5 was biotinylated (biotin/antibody ratio = 30/1) and used as the detector at a concentration of 0.3ug/mL. The SBG revelation enzyme concentration was 150pM. Recombinant IFNα17 was used as calibrator. The limit of detection was calculated by the mean value of all blank runs + 2SD after log conversion. The assay is fully described in Rodero *et al*.^13^ and a video presenting all the steps is shown in Llibre *et al* ^19^.

### IFNα2 digital ELISA assay

The Simoa IFNα2 assay was also developed using the Quanterix Homebrew kit. The BMS216C (eBioscience) antibody clone was used as a capture antibody after coating on paramagnetic beads (0.3mg/mL), and the BMS216BK already biotinylated antibody clone was used as the detector at a concentration of 0.3ug/mL. The SBG revelation enzyme concentration was 150pM. Recombinant IFNα2c was used as calibrator. The limit of detection was calculated by the mean value of all blank runs + 2SD after log conversion.

### ISG score determination

For SLE and CTD patients ISG score was calculated by qPCR as previously described using six ISG genes (Rice *et al.*^*20*^). Blood was collected in PAXgene tubes (PreAnalytix), and total RNA was extracted using a PAXgene (PreAnalytix) RNA isolation kit. RNA concentration was assessed by fluorimetric analysis. After DNAse treatment, RNA was converted into cDNA with the High Capacity cDNA reverse transcriptase kit (Qiagen). Quantitative reverse transcription polymerase chain reaction (RT-qPCR) analysis was performed using the TaqMan Universal PCR Master Mix (Applied Biosystems) and cDNA derived from 40ng total RNA. Using TaqMan probes for IFI27 (Hs01086370_m1), IFI44L (Hs00199115_m1), IFIT1 (Hs00356631_g1), ISG15 (Hs00192713_m1), RSAD2 (Hs01057264_m1), and SIG LEC1 (Hs00988063_m1), the relative abundance of each target transcript was normalized to the expression level of HPRT1 (Hs03929096_g1) and 18S (Hs999999001_s1) and assessed with the Applied Biosystems StepOne Software v2.1 and DataAssist Software v.3.01. For each of the six probes, individual (patient and control) data were expressed relative to a single calibrator (control C25). The median fold change of the six ISGs, when compared with the median of the combined 29 healthy controls, was used to create an IFN score for each patient. RQ is equal to 2^−ΔCt^ (i.e., the normalized fold change relative to a control). All patients were selected for this analysis, except the ASSESS ones for which the protocol was different.

For pSS patients the ISG-score was determined from microarray data. After validation of the RNA quality with Bioanalyzer 2100 (using Agilent RNA6000 nano chip kit), 75 ng of total RNA was reverse transcribed using the GeneChip® WT Plus Reagent Kit (Affymetrix) without globin mRNA reduction. Briefly, the resulting double-stranded cDNA was used for *in vitro* transcription with T7 RNA polymerase (all these steps are included in the WT cDNA synthesis and amplification kit of Affymetrix). After purification according to Affymetrix’s protocol, 5.5 μg of Sens Target DNA were fragmented and biotin-labelled. After control of fragmentation using Bioanalyzer 2100, cDNA was then hybridized to GeneChip® Clariom S Human (Affymetrix) at 45°C for 17 hours. After overnight hybridization, chips were washed on the GeneChip Fluidics Station 450 following specific protocols (Affymetrix) and scanned using the GCS3000 7G. The scanned images were then analyzed with Expression Console software (Affymetrix) to obtain raw data (.cel files) and Quality Control metrics. Computational analyses were performed on R 3.4.3. Microarray data were loaded and normalised using the oligo package. Briefly .CEL files were loaded using the read.celfiles() functions and normalised using the Robust Multi Array algorithm implemented in the rma() function with default parameters. Quality of data normalization was checked through visual inspection of intensity distribution across chips. Microarrays probes were annotated using the clariomshumanhttranscriptcluster.db package and if several probes were mapped to the same gene, the probe with the highest coefficient of variation was selected. Then, the ISG score for each patient was calculated as the median of normalized data available for the same genes, i.e. IFI27, IFI44L, IFIT1, ISG15, RSAD2 and SIG LEC1 as previously defined (Rice *et al.*^*20*^).

### IFN-activity determination

Type I IFN activity was measured by determining the cytopathic reduction (i.e., protection of Madin–Darby bovine kidney (MDBK) cells against cell death after infection with vesicular stomatitis virus) afforded by patient serum. A reference of human IFNα, standardized against the National Institutes of Health reference Ga 023–902-530, was included with each titration. IFNα activity in normal healthy serum is <2IU/ml. Patients were selected randomly from the SLE, CTD and pSS cohorts to equilibrate the number of results in each cohort and at each IFNα17/α2c protein ratio (n=113).

### Competition assays

SLE patient plasma samples or pure recombinant IFNα subtypes were diluted with Detector / Sample Diluent (Quanterix) and preincubated with 50μg/ml of the IFNα capture antibody 8H1 or BMS216C (eBioscience) for 30min at room temperature before Simoa analysis.

### Anti-IFNα autoantibodies quantification

Anti-IFNα (1,2,8,21) and anti-IFNα (4,5,6,7) quantifications were done in each LEAP SLE, LEAP CTD and ASSESS Sjögren’s patients cohorts from a similar restricted number of samples, randomly selected (n=113). Autoantibodies against IFNα were measured with luciferase based immunoprecipitation assay (LIPS) as described previously (Meyer *et al.^12^*). IFNA subtype sequences were cloned into modified pPK-CMV-F4 fusion vector (PromoCell GmbH, Germany) where Firefly luciferase was substituted in the plasmid for NanoLuc luciferase (Promega, USA), transfected to HEK293 cells and IFNα-luciferase fusion proteins collected with tissue culture supernatant. 1×10^6^ luminescence units (LU) of IFNα1, IFNα2, IFNα8 and IFNα21 fusion proteins were combined to one IP reaction (pool 1), and IFNα4, IFNα5, IFNα6 and IFNα7 fusion proteins to another (pool 2). Serum samples were incubated with Protein G Agarose beads (Exalpha Biologicals, USA) at room temperature for 1 h in 96-well microfilter plate (Merck Millipore, Germany) before the antigen mix was added for another hour. After washing the plate with vacuum system, Nano-Glo^®^ Luciferase Assay Reagent was added (Promega, USA). Luminescence intensity was measured by VICTOR X Multilabel Plate Reader (PerkinElmer Life Sciences, USA). The results were expressed in arbitrary units (AU) representing fold over the mean of the negative control samples. The cut-off was set at 2.0 AU. Anti-IFNα8 and anti-IFNα2 quantifications were done for five patients with high autoantibody values studied longitudinally. The previous protocol is employed using only the IFNα8 and IFNα2 fusion proteins. Serial dilutions from serum samples are co-incubated with fixed concentrations of these antigens. IC50 is calculated based on the dose-response curve. This is the dilution factor that halves the signal or reduces IFN activity 50% from its maximum.

### Statistical analyses

GraphPad Prism was used for statistical analysis. A Mann-Whitney test was used to test for differences induced by the buffer used to treat samples. ANOVA tests (Kruskal–Wallis) with Dunn’s post testing for multiple comparisons were used to test for differences between patients groups, or stimulation groups. *p< 0.05; **p<0.01; ***p<0.001; ****p<0.0001. A Chi square test was used to characterize the differences between two distributions of results. For all analyses, p values less than 0.05 were considered statistically significant. Median and 95% confidence interval were reported. Spearman correlations are used to compare the two IFNα assays.

## Data availability

All available patient data is shown in Supplementary Table S1.

## Acknowledgements

We thank all patients and healthy donors who provided samples used in this study, as well as the clinical support teams that enabled their collection. We thank Immunoqure for provision of mAbs for the pan-IFNα assay. We acknowledge support from the ANR (CE17001002) awarded to DD.

## Results

### Comparison of IFNα2 and pan-IFNα digital ELISA assays

To enable the study of IFNα protein subtypes we developed a Simoa digital ELISA utilizing a monoclonal antibody (mAb) pair specific for IFNα2. This assay had a limit of detection (LOD) of 1fg/ml and no cross-reactivity for IFNβ, IFNλ1, IFNλ2, IFNω, and IFNγ (Fig 1a). This level of sensitivity, and lack of cross-reactivity, were similar to our previously described pan-IFNα Simoa assay that utilized mAbs isolated from APECED patients (Fig 1b). We also tested the reactivity of the new IFNα2 assay for all IFNα 14 subtypes which revealed differing levels of specificity, with the highest specificity for IFNα2 and IFNα6 (Fig 1c). This was in contrast to pan-IFNα assay which has equally high specificity for all IFNα subtypes, with the exception of IFNα2 (Fig 1d).

**Figure 1.**
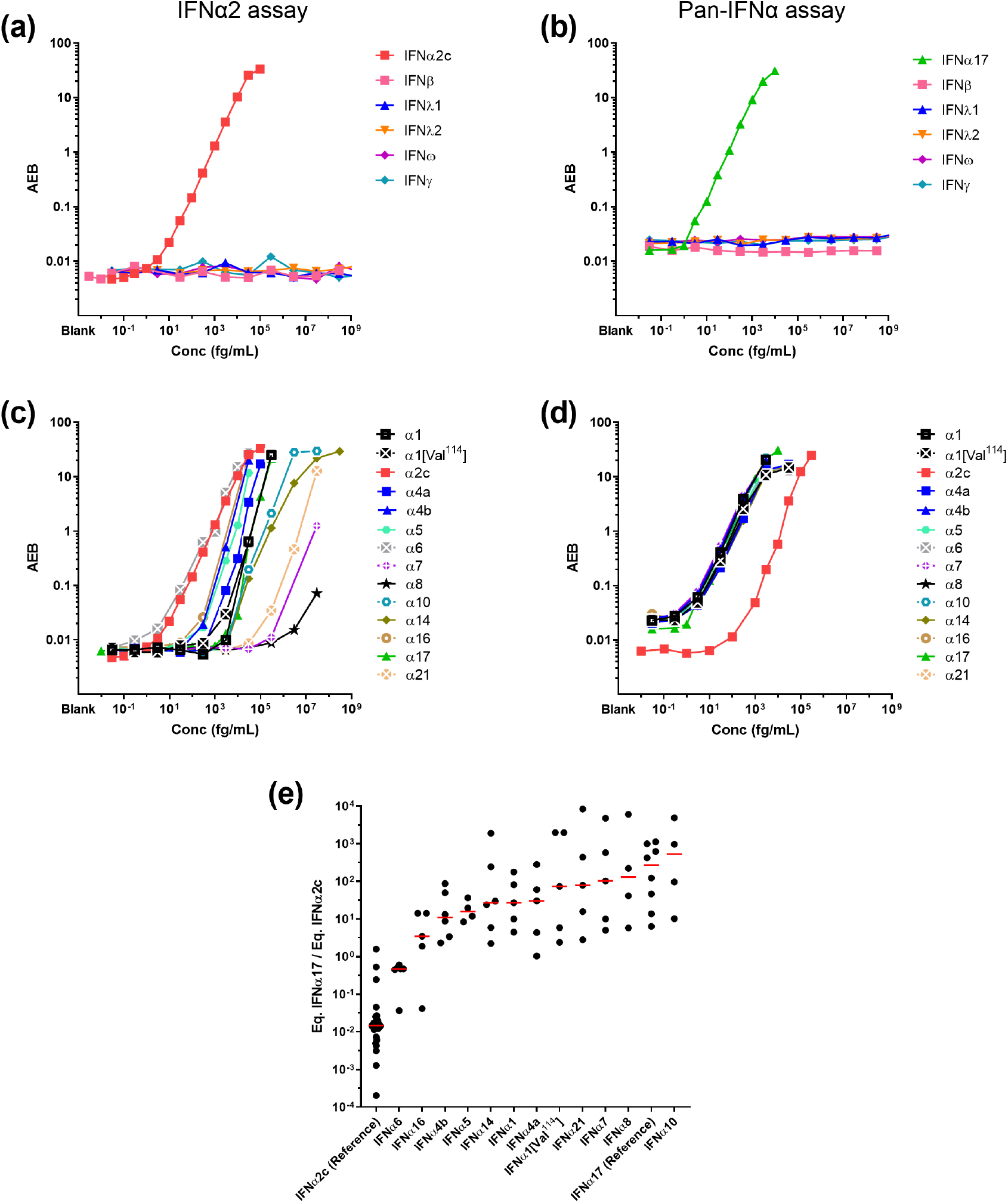
Digital ELISAs to study IFNα subtypes. Response (Average Enzyme per Bead) of the IFNα2 assay (a) and pan-IFNα (b) at different concentrations for the recombinant proteins IFNβ, IFNλ, IFNω and IFNγ. Response (Average Enzyme per Bead) of the IFNα2 assay (c) and pan-IFNα (d) at different concentrations for recombinant IFNα subtypes and the natural variant IFNα1(Val114). (e) The IFNα17/IFNα2c ratio for different concentrations of recombinant IFNα subtypes and the natural variant IFNα1(Val114) as measured by both the IFNα2 assay and pan-IFNα assays.

To explore this differential specificity between the two assays for the IFNα subtypes, we tested each of the 13 (and the natural variant IFNα1/Val114) IFNα subtypes at different concentrations, from no response to saturation, and plotted the ratio of the results obtained in the two assays (Fig 1e). Results are expressed as concentrations after calibration, to IFNα17 for the pan-IFN assay and IFNα2 for the IFNα2 assay. This range of values therefore captures the possible biological distributions of IFNα17/α2, with ratios ranging from 10^4^ to 2×10^−4^. Recombinant IFNα2 presented a low ratio as it was recognized with high specificity by the IFNα2 assay, but with low specificity by the pan-IFNα assay. IFNα6 had an intermediate ratio as it was recognized equally well by both assays, and all other subtypes presented a ratio higher than 1 due to greater specificity with the pan-IFNα assay.

### TLR activation can modify the balance of IFNα protein subtypes

To test the potential biological relevance of these observations we tested the hypothesis that different TLR stimulation could modify the balance of IFNα subtypes. To do this, we stimulated whole blood and purified plasmacytoid Dendritic Cells (pDCs) (the main IFNα producing cell type) with different TLR ligands and quantified IFNα in the supernatant with both assays. Whole blood from healthy donors was stimulated with LPS (TLR4), Poly(I:C) (TLR3), and R848 (TLR7/8) plus a Null control for 22 hours using concentrations previously defined to be within the linear range of response ^21^. Response with both assays correlated extremely well as expected (Rs=0.96, p<0.0001 Spearman test), with R848 showing the strongest induction followed by equivalent responses for LPS and Poly(I:C) (Fig 2a). Purified pDCs from healthy donors were stimulated with relevant ligands for TLR9, their main innate sensor, namely CpG-A and cGAMP, as well as LPS (TLR4), Poly(I:C) (TLR3), and R848 (TLR7/8). Strong responses were observed after stimulation with all TLR9 agonists and R848, with a good positive correlation between both assays (Rs=0.79, p<0.0001 Spearman test) (Fig 2b). A high background (10^4^ fg/mL) is observed using the IFNα2 assay, probably due to non-specific cell activation due to experimental manipulation. Because this non-specific activation could modify the results, we next calculated the IFNα17/α2 ratio between the different stimulation conditions for samples that were effectively stimulated by our molecules (significant difference and fold-induction compared to non-stimulated condition). For whole blood we observed a significantly lower ratio (p=0.0008) between R848 and LPS and Poly(I:C) (Fig 2c). For pDCs we observed a significantly (p=0.007) higher ratio between TLR9 and TLR7/8 ligands (Fig 2d). Therefore, in both whole blood and pDC responses to stimulation our assays could detect differential induction of IFNα protein subtypes depending on the TLR pathway targeted validating the biological relevance of this approach.

**Figure 2.**
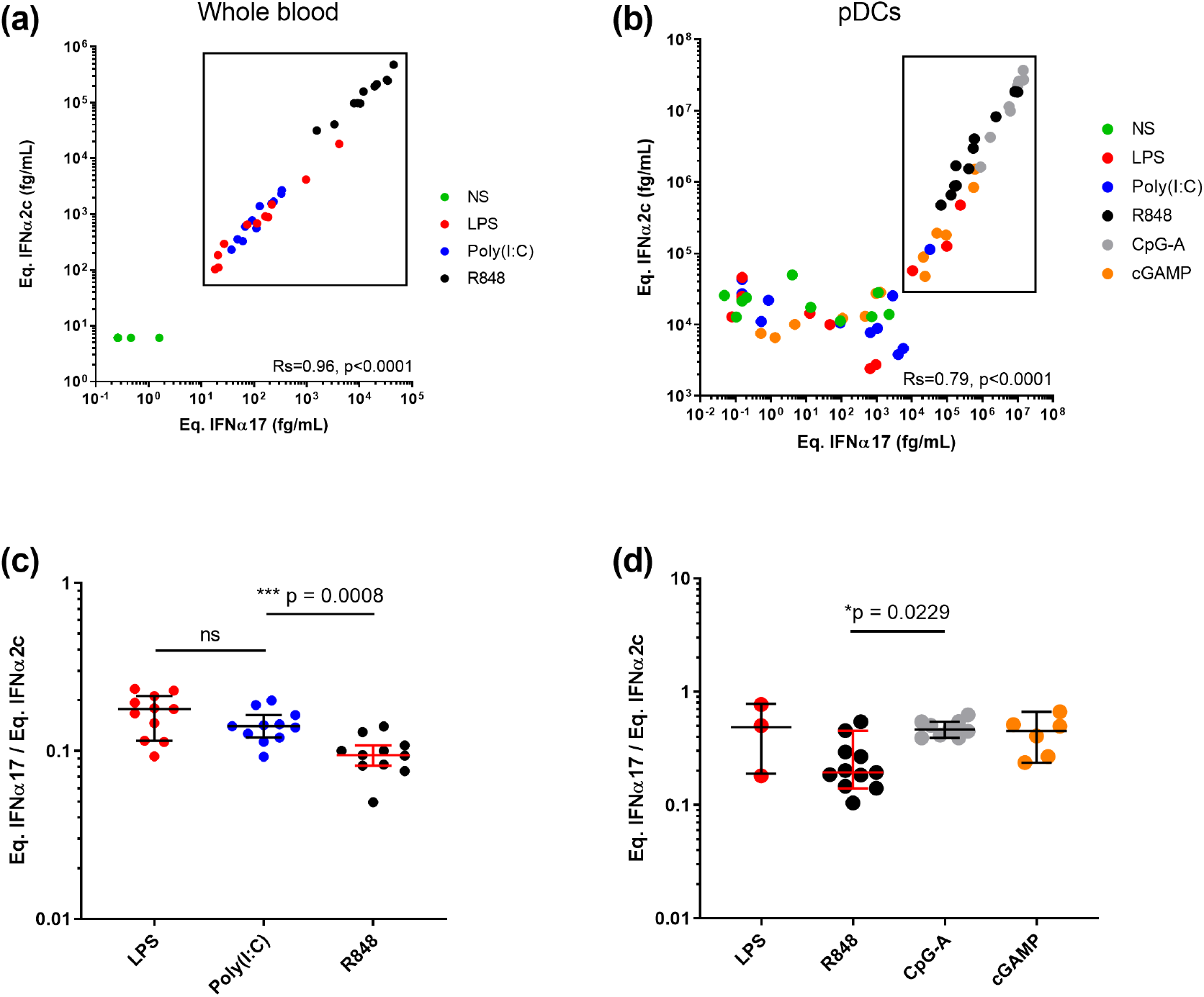
TLR activation can modify the IFNα subtype balance. IFNα concentrations obtained using the IFNα2 and pan-IFNα assays after (a) whole blood stimulation with LPS (TLR4), Poly(I:C) (TLR3) or R848 (TLR7/8) and (b) pDC stimulation with LPS, Poly(I:C), R848, CpG-A (TLR9) or cGAMP. Comparison of the IFNα17/α2 protein ratios obtained from (c) whole blood and (d) pDC stimulation with different agonists. Kruskal-Wallis test with Dunn’s correction for multiple comparisons and Spearman correlations are reported.

### Comparison of IFNα subtype protein ratios in autoimmune and infection

To test the potential clinical relevance of the IFNα17/α2 ratio in disease we applied both IFNα2 and pan-IFNα assays to plasma from patients with acute (dengue and viral central nervous system infection) or chronic (HCV) viral infection, as well as autoimmune conditions; systemic lupus erythematosus (SLE), connective tissue disease (CTD), and primary Sjögren’s Syndrome (pSS), all pathologies previously shown to be associated with increased IFN signaling. Both assays showed an overall strong and positive correlation (Rs=0.6198 p<0.0001, Spearman test), however there were certain patients from both chronic viral infection and autoimmune groups whose IFNα results did not correlate well between the two assays (Fig 3a).

**Figure 3.**
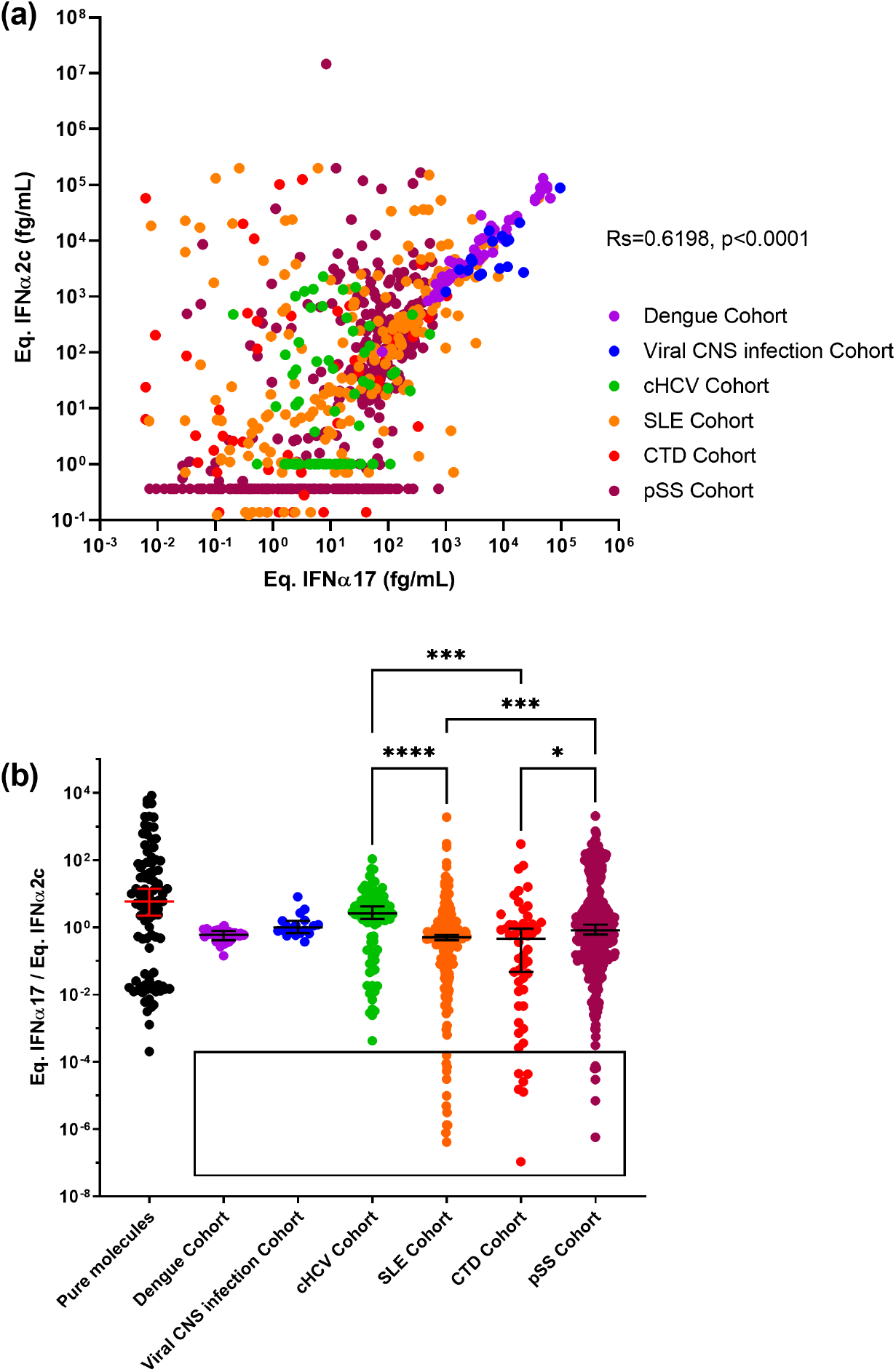
IFNα subtype protein ratios in autoimmune and infection. IFNα concentrations obtained using the IFNα2 and pan-IFNα assays in plasma samples from patients with acute dengue infection, chronic HCV infection (cHCV), systemic lupus erythematosus (SLE), connective tissue disease (CTD), and primary Sjögren’s syndrome (pSS). Values for viral central nervous system (CNS) infections are obtained from cerebrospinal fluid samples (a). The IFNα17/IFNα2c ratio as measured by both the IFNα2 and pan-IFNα assays for different concentrations of recombinant IFNα subtypes (left) and for the same samples from patients with acute dengue infection, viral central nervous system (CNS) infections, chronic HCV infection (cHCV), systemic lupus erythematosus (SLE), connective tissue disease (CTD), and primary Sjögren’s syndrome (pSS) (b). Kruskal-Wallis test with Dunn’s correction for multiple comparisons and Spearman correlations are reported.

To explore this further we examined the IFNα ratio as previously described. Patients with acute viral diseases – dengue or CNS infections – showed the narrowest distribution range of IFNα17/α2 ratios with intermediate medians at 0.6 and 1.0 respectively (Fig 3b). In contrast, patients chronically infected with HCV showed a wider distribution range of IFNα17/α2 ratios from 2×10^−4^ to 10^2^ and a median of 2.6 suggesting the presence of multiple IFNα subtypes. Strikingly the three autoimmune cohorts all showed a similar wide distribution range with median values from 0.46 to 0.82. Intriguingly the autoimmune cohorts also revealed interesting sub-groups of outlier individuals with extremely low IFNα17/α2 ratios below 2 x10^−4^. These represented 3% of the total patients tested. A Kruskal–Wallis test with Dunn’s post testing for multiple comparisons revealed significant differences between the SLE patients and pSS and cHCV patients, with a greater presence of low values in patients with autoimmune diseases suggesting a higher proportion of IFNα2 protein. These results collectively suggest that the level of disease complexity could explain the scale of the ratio distribution.

### Patients with extremely low IFNα17/α2 protein ratio display reduced IFN signaling and function

The identification of SLE, CTD and pSS patients with an IFNα17/α2 ratio below 2×10^−4^ was intriguing, as results with recombinant molecules showed the extreme low probability of this phenotype occurring (Fig 1e). Our SLE, CTD and pSS patient cohorts contained respectively 12/206 (5,8%), 6/52 (12%) and 6/392 (1,5%) samples with this phenotype (Fig 4a, 4d, 4g). To examine a potential functional relevance for this abnormal ratio we plotted an interferon stimulated gene (ISG) score calculated by qPCR or microarray (previously described) against the individual equivalent IFNα17 (Fig 4b, 4e and 4h) and IFNα2c (Fig 4c, 4f, 4i) concentrations. This analysis revealed that for patients with low IFNα17/α2 ratios (red dots), the pan-IFNα value positively correlated as expected with the ISG score, and in line with other patients (blue dots). In contrast, these patients showed abnormally low ISG scores in regard to their IFNα2 results (orange dots). This suggested that in certain patients the IFNα2 protein value did not well reflect *in vivo* activity. To further test this hypothesis, we performed an IFN cytopathic assay on a subset of samples selected randomly from each cohort and at each IFNα17/α2c protein ratio. This revealed that patients with extremely low IFNα17/α2 ratios, also had low IFN activity (Fig 4j and 4k), despite these samples having apparent high IFNα2 protein concentrations.

**Figure 4.**
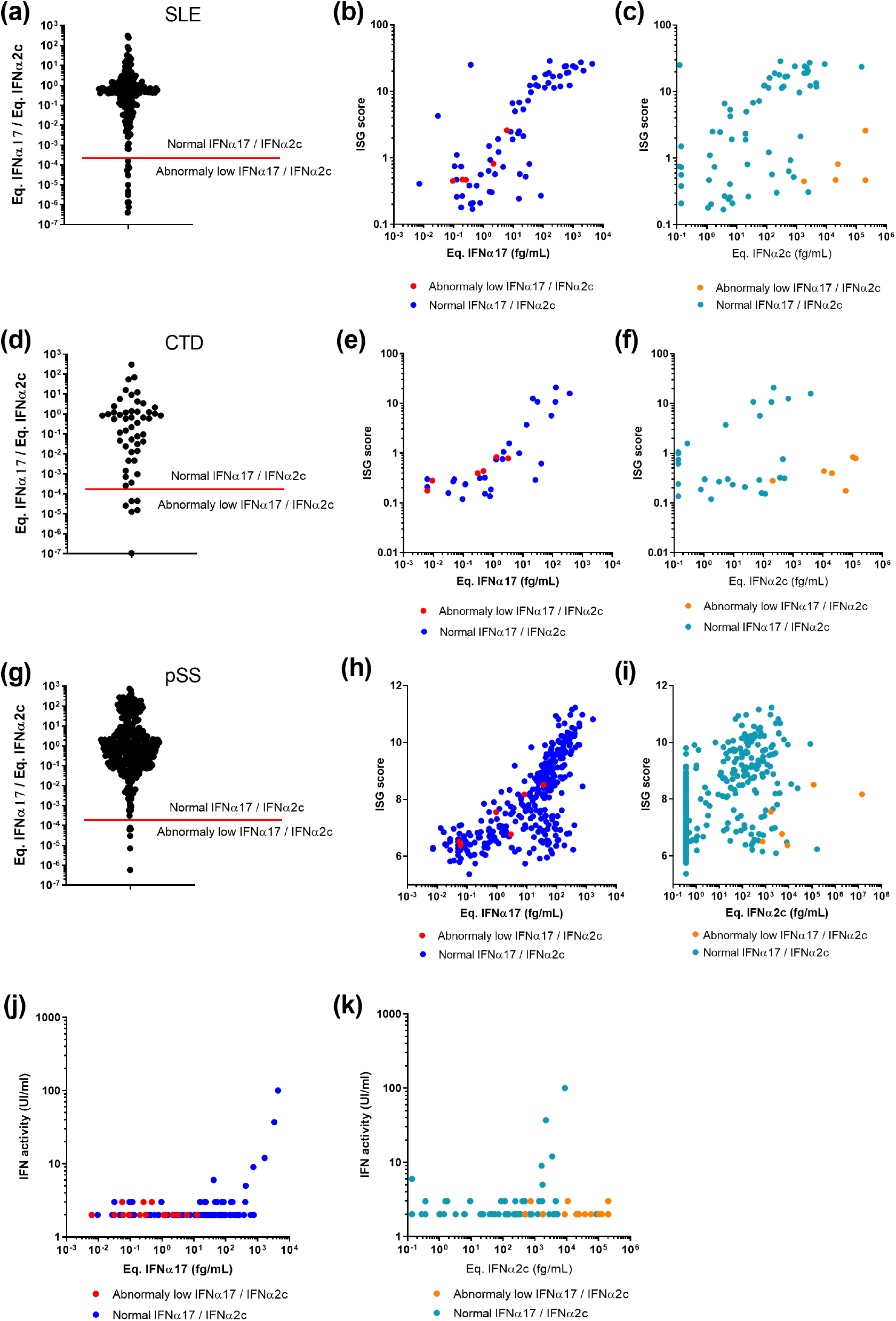
Integration of IFNα17/α2 protein ratios with ISG score and functional activity. IFNα17/α2 protein ratios obtained from samples of patients with (a) SLE, (d) CTD and (g) pSS. The red bar indicates the lower IFNα17/α2 protein ratio using pure molecules. Correlation of ISG score with pan-IFNα assay (b, e, h) or IFNα2 assay results (c, f, i) for same patients. Correlation of IFN functional activity with pan-IFNα assay (j) or IFNα2 assay results(k) for same patients. Patients with an IFNα17/α2 protein ratio threshold above the 2.10^−4^ cut off are in blue, those below in red.

### Extremely low IFNα17/α2 protein ratio is not due to non-specific effects

A possible technical explanation for these unusual observations was non-specific binding within the assays. To test this hypothesis, we re-analyzed a series of 32 SLE samples in which the proportion of samples showing extremely low IFNα17/α2 ratios was increased in an assay buffer (Buffer B, Quanterix) specifically developed for low background. We have previously shown that this buffer successfully suppresses non-specific binding in an IFNβ assay ^15^. This revealed identical results (no difference between the buffer groups, p=0.1103, Fig S1), and moreover samples with a low IFNα17/α2 protein ratio maintained this phenotype irrespective of the buffer used (surrounded points), indicating that the phenotype was not due to non-specific binding.

### Auto-antibodies against IFNα can explain the abnormal IFN subtype ratios

A potential explanation for the extreme IFNα17/α2 ratio is the presence of auto-antibodies, which have been reported to develop against cytokines in autoimmune patients. To test this possibility, we performed a series of *in vitro* experiments. The addition of the 8H1 pan-alpha capture antibody in pure IFNα17 solutions at different concentrations reduces the IFNα17/α2 ratio (Fig 5a). The anti-IFNα2c capture antibody produces no effect in these solutions but, added into IFNα2c solutions, the antibody increases the ratio (Fig 5b). And the addition of 8H1 in a selection of five SLE samples reduces the IFNα17/α2 ratio (Fig 5c). This demonstrates that the presence of anti-IFNα auto-antibodies in biological samples could modify the ratio.

**Figure 5.**
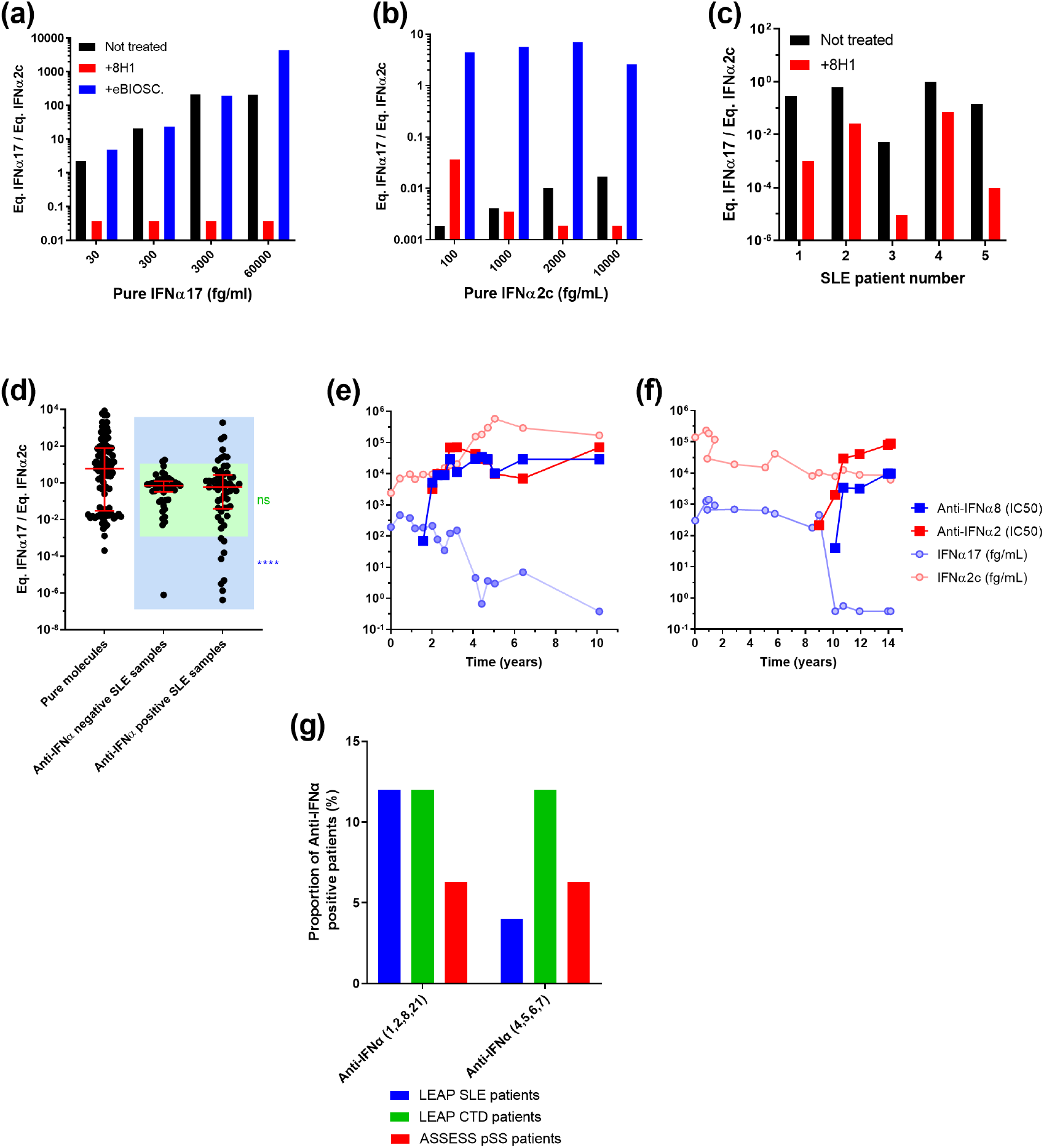
Association of anti-IFNα auto-antibodies and the abnormal IFNα subtype ratios. (a) Competition assays after addition of the 8H1 clone anti-IFNα antibody in pure IFNα17 solutions at different concentrations reduces the ratio. (b) Addition of the anti-IFNα2 antibody in pure IFNα2c solutions at different concentrations increases the ratio. (c) Addition of 8H1 in five SLE patient samples reduces the IFNα17/α2 ratio. (d) IFNα17/α2 protein ratios for SLE patient samples characterized positive for anti-IFNα antibodies in comparison with negative SLE patient samples and pure molecules. (e-f) Longitudinal analysis of anti-IFNα8 and anti-IFNα2 antibody concentrations, pan-IFNα and IFNα2 assays results over time (years) in two SLE patients. (g) Proportion of anti-IFNα auto-antibodies positive samples in LEAP SLE, LEAP CTD and ASSESS pSS patients.

Therefore, we tested SLE patients for auto-antibodies against IFNα (subtypes 1,2,4,5,6,7,8 and 21) in their serum. A comparison of the auto-antibody negative and positive patients using the Chi square test revealed a higher proportion of extremely low IFNα17/α2 ratios in the anti-IFNα positive group supporting this hypothesis (Fig 5d). We next examined SLE patients for whom longitudinal sampling was available over multiple years, which revealed interesting inter-patient variability. Strongly supporting our hypothesis for the role of anti-IFNα autoantibodies, certain patients showed continuous high levels of IFNα2 and initial high levels of pan-IFNα that suddenly dropped to undetectable concentrations. Strikingly these sudden undetectable pan-IFNα concentrations occur in parallel to detectable presence of auto-antibodies against IFNα2 and IFNα8. Two patients (Fig 5e and 5f) out of 11 patients examined showed this pattern of high IFNα2, and low pan-IFNα, in the presence of anti-IFN autoantibodies. One other patient with high longitudinal anti-IFN auto-antibodies did not have elevated levels of IFNα protein at any time point examined (Fig S2). The other eight patients showed lower anti-IFNα auto-antibodies concentrations. Finally, we tested the LEAP SLE, the LEAP CTD and the ASSESS pSS cohorts for anti-IFNα (1,2,8,21) and anti-IFNα (4,5,6,7) autoantibodies to estimate the frequency of this phenotype. 12% of the SLE and CTD patients, and 6.3% of the pSS patients produce anti-IFNα auto-antibodies (Fig 5g).

## Discussion

Type I interferons are essential for anti-viral immunity as recently highlighted in studies of severe COVID-19 disease ^22,23^. These studies and others have highlighted the need to directly quantify the cytokine protein in patient samples, which has been made possible by the recent development of digital ELISA ^11^. However new tools to differentiate the diverse IFNα subtypes are required to further our understanding of their potential differential functions. We illustrate here how two complementary anti-IFNα assays can be used in tandem to reveal differences in IFNα subtype biology. This included subtle but significant differences in subtype expression between acute and chronic viral infection, and autoimmune diseases, and also after TLR stimulation of whole blood and isolated pDCs. However, our results also reveal discrepancies that can occur depending on the specificity of the monoclonal antibodies used.

In our originally described Simoa pan-IFNα assay ^13^, 8H1 and 12H5 antibody clones were selected because of their ability to quantify all IFN alpha subtypes. Isolated from APECED patients ^12^, these mAbs also have extremely high affinity and neutralizing capacity making them good candidates for therapeutic use. These attributes help to explain the good correlations observed between results with this assay and ISG scores and IFN-activity across all patient groups. In contrast, the BMS216C and BMS216BK antibody clones used in the IFNα2 assay were developed in mice for ELISA applications. As such they recognize human-characteristic epitopes on IFNα molecules, that are likely distinct from the functional binding sites. As a result, we observed lower correlations between IFNα2c concentrations measured with this assay and ISG score or IFN-activity. In support of these hypotheses, we compared the amino acid sequences and three-dimensional structures published online in the PDB database (www.rcsb.org) with the reactivity of all IFN molecules we tested in our two assays. From this, we estimated the pan-alpha assay epitope to be the ^100^MQEVGV sequence in the IFNα17 molecule, and the IFNα2 assay epitope to be the ^110^LMKED sequence in the IFNα2 molecule. Akabayov *et al* ^24^ previously determined the IFNα2 amino-acids implicated in binding with the IFNAR1 receptor. From this, our estimated pan-IFNα epitope, but not the predicted IFNα2 epitope, includes a high concentration of IFNAR1 receptor binding amino-acids. These observations are consistent with the fact that the pan-IFNα assay uses neutralizing antibodies, but not the IFNα2 assay. These epitope considerations could also explain the results observed in longitudinal patients, where autoantibodies against IFNα8 may compete with the pan-IFNα antibodies for the functional epitopes to reduce the IFNα concentrations measured. In contrast, the autoantibodies may not block the epitopes recognized by the IFNα2 assay resulting in no modification of the IFNα2 concentrations measured. This implicates that autoantibodies have a neutralizing efficiency, a hypothesis that could be verified measuring ISG score or IFN-activity in these longitudinal samples.

Autoimmune disorders are well known to induce production of autoantibodies ^25^. Nevertheless, their presence can be detrimental in cases of acute viral infection where type I interferon responses are crucial for host protection, as recently demonstrated for COVID-19 ^23^. We observed anti-IFNα autoantibody positivity in 12% of SLE and CTD groups, and 6% in the pSS group. Gupta *et al.*^26^ previously quantified 11% in SLE patients and 9% in pSS patients which is in line with our results. We also observed extremely low IFNα ratios for autoimmune patients, which were absent in the cHCV, the dengue or the viral CNS infection patients. The proportion of these low ratio patients was 3%, lower than that observed for overall anti-IFNα autoantibody positivity. These differences may be due to autoantibodies targeting multiple distinct IFNα epitopes, or that extremely low ratios are only one possible consequence of autoantibody production, and explains why only two longitudinal patients out of 11 showed this phenotype.

Finally, this work highlights the additional information that can be obtained from using two different ELISA assays to quantify a single cytokine. The presence of autoantibodies, or enzymatic cleavage as previously described ^27^, could potentially decrease protein activity without decreasing the overall protein concentration. Autoantibodies could also reduce an ELISA measurement without decreasing the actual protein concentration, e.g. if the protein is captured to circulating immune complexes. Real or active concentrations in the presence of autoantibodies are not accessible using a single ELISA, depending on the epitope recognized by the mAbs. The use of a second ELISA assay may be required, or alternatively a functional activity assay for confirmation. Additional approaches such as proximity extension assays may also potential overcome these issues through the use of two complementary Abs for double target recognition^28^. The relevance of these issues is increased for cytokines that show multiple subtypes quantified using ELISA assays that show a particular response for each subtype. It becomes impossible to compare two results without the verification that the subtype balance is the same in the two related samples. However, across the multiple patient cohorts we examined the frequency of this phenotype was relatively low, at only 3% of total patient samples examined. Therefore, once mAbs are verified for specificity and cross-reactivity such challenges may only be encountered when testing large patient cohorts.

## Individual contributions to this work

**Vincent Bondet**: Conceptualization, Methodology, Validation, Formal analysis, Investigation, Data curation, Writing - original draft, Writing - review & editing, Visualization. **Mathieu P. Rodero**: Conceptualization, Validation, Resources, Writing - review & editing. **Céline Posseme**, **Jérémie Decalf**, **Pierre Bost**, **Liis Haljasmägi**, **Nassima Bekaddour, Gillian Rice, Vinit Upasani**: Investigation, Writing - review & editing. **Jean-Philippe Herbeuval**, **John A Reynolds**, **Tracy A. Briggs**, **Ian N. Bruce**, **Claudia Mauri**, **David Isenberg**, **Madhvi Menon**, **David Hunt**, **Benno Schwikowski**, **Xavier Mariette**, **Stanislas Pol, Tineke Cantaert, J. Eric Gottenberg, Flore Rozenberg**: Resources, Validation, Investigation, Writing - review & editing. **Kai Kisand**: Conceptualization, Validation, Investigation, Resources, Writing - review & editing. **Darragh Duffy**: Methodology, Writing - original draft, Writing - review & editing, Supervision, Project administration, Funding acquisition.

## Supplementary material

**Figure S1.**
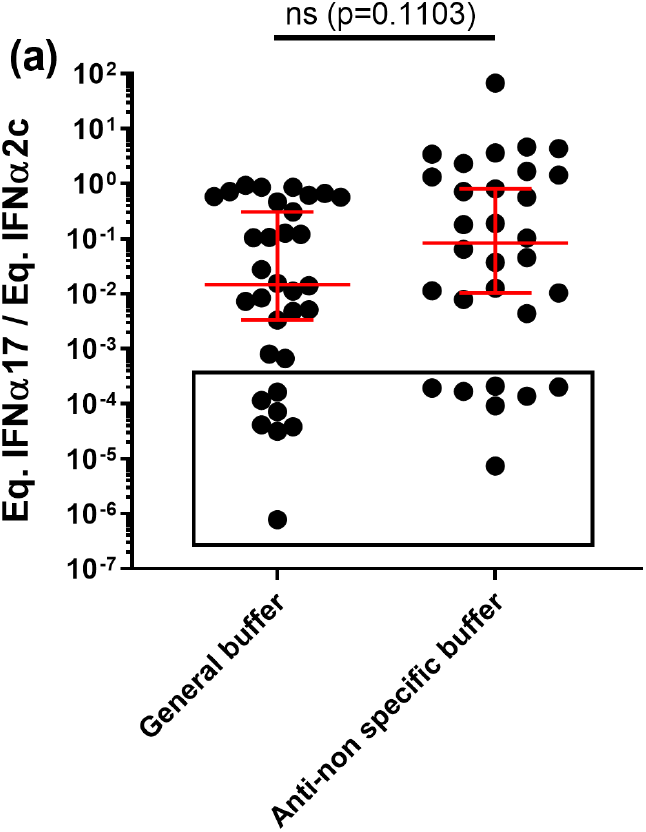
Abnormal IFN subtype ratios are not due to unspecific reactions. IFNα17/α2 protein ratios obtained using general Detector / Sample Diluent or buffer B for a selection of anti-IFNα positive and anti-IFNα negative patients.

**Figure S2.**
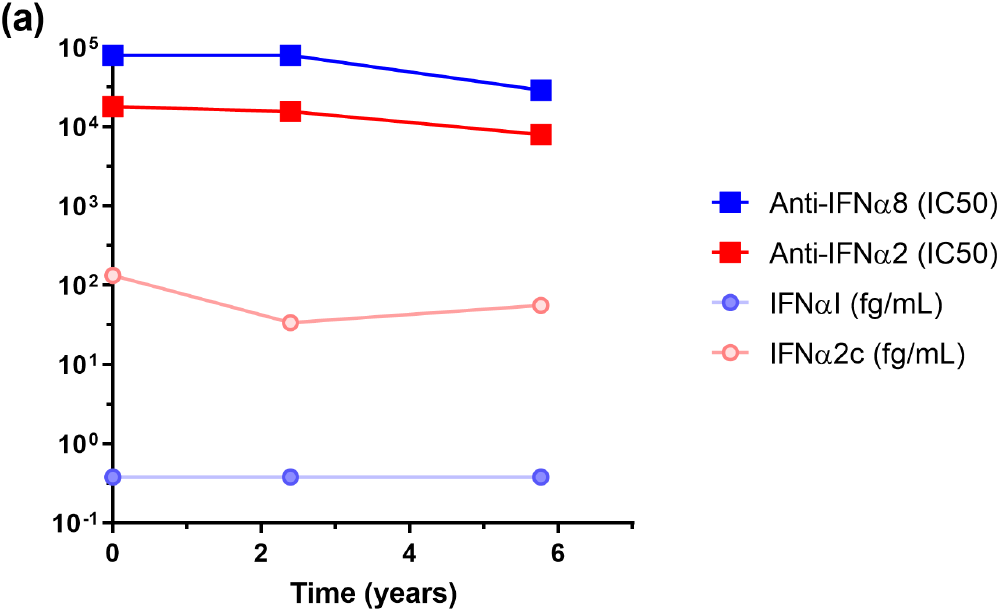
Longitudinal analysis of anti-IFNα8 and anti-IFNα2 antibody concentrations, pan-IFNα and IFNα2 assays results over time (years) in one other SLE patient.

**Table S1. Patient data sets.** For each patient included in this study, origin cohort, inclusion cohort, diagnosis, gender, age, autoantibodies presence, IFNα17 and IFNα2 concentrations, IFNα17/IFNα2 ratio, ISG score, IFN activity, and quantification results for antibodies against IFNα are done when available. LEAP (lupus extended autoimmune phenotype) cohort: see Reynolds *et al.*^16^. Rodero *et al.* cohort: see reference 13. ASSESS (assessment of systemic complications (signs) and evolution in Sjögren’s syndrome) cohort: see Bost *et al.* (In review). Menon *et al.* cohort: paper in preparation. C10-08 cohort: see Sultanik *et al.*^17^. Upasani *et al.* cohort: see reference 18. UCDT: undifferentiated CTD. MCTD: mixed CTD. CTD: connective tissue disease. pSS: primary Sjögren’s syndrome. SLE: systemic lupus erythematosus. JSLE: juvenile SLE. cHCV: chronic hepatitis C virus infection. CNS: central nervous system.

