## Supplementary Table 1 for "Differential levels of IFNα subtypes in autoimmunity and viral infection"

**Table S1 : Patient data sets**

| |
|---|
| Limit of detection, Negative |
| - : Not determined |
| na : Not available |

| Patient_id | Cohort | Origin | Short diagnosis | Gender | Age | IFNα17 (fg/mL) | IFNα2c (fg/mL) | IFNα17/IFNα2c | ISG Score | IFN activity (UI/ml) | anti-IFNa (AU) | Anti-IFNa (1,2,8,21) | Anti-IFNa (4,5,6,7) |
|---|---|---|---|---|---|---|---|---|---|---|---|---|---|
| 28 | CTD | LEAP Cohort | UCTD | F | 55 | 3,2394383535 | 125491,365904049 | 0,000025814 | 0,789 | 2 | - | 0,72 | 0,50 |
| 31 | CTD | LEAP Cohort | UCTD | F | 43 | 0,5333176606 | 115,6016027106 | 0,0046134106 | 0,153 | - | - | - | - |
| 33 | CTD | LEAP Cohort | UCTD | F | 30 | 2,245732631 | 0,1387143064 | 16,1896252012 | 1,064 | 2 | - | 0,51 | 0,50 |
| 34 | CTD | LEAP Cohort | UCTD | F | 35 | 7,5563166843 | 0,1387143064 | 54,4739535473 | 0,995 | 2 | - | 0,72 | 0,51 |
| 35 | CTD | LEAP Cohort | UCTD | F | 61 | 0,117449015 | 0,1387143064 | 0,8466972014 | 0,241 | - | - | - | - |
| 36 | CTD | LEAP Cohort | UCTD | F | 27 | 21,4993418984 | 683,5903411453 | 0,0314506227 | 12,578 | 2 | - | 0,47 | 0,39 |
| 39 | CTD | LEAP Cohort | UCTD | F | 41 | 26,3472785379 | 72,5980869611 | 0,3629197358 | 0,29 | 2 | - | 0,61 | 0,71 |
| 40 | CTD | LEAP Cohort | UCTD | F | 49 | 0,006210424 | 58013,7316137215 | 1,07050932143524E-007 | 0,176 | 2 | - | 0,61 | 0,57 |
| 41 | CTD | LEAP Cohort | UCTD | F | 29 | 0,006210424 | 6,3598799532 | 0,0009765002 | 0,302 | 2 | - | 0,63 | 0,46 |
| 44 | CTD | LEAP Cohort | MCTD | F | 51 | 90,3761395853 | 75,775178075 | 1,1926879208 | 5,676 | - | - | - | - |
| 45 | CTD | LEAP Cohort | MCTD | F | 32 | 128,4280321963 | 220,7554303601 | 0,5817661291 | 20,922 | 2 | - | 0,71 | 0,47 |
| 47 | CTD | LEAP Cohort | UCTD | F | 61 | 0,787664032 | 0,1387143064 | 5,6783186419 | 0,136 | 2 | - | 0,49 | 0,49 |
| 52 | CTD | LEAP Cohort | UCTD | F | 26 | 123,2372946237 | 185,0496789789 | 0,6659687026 | 10,772 | - | - | - | - |
| 55 | CTD | LEAP Cohort | MCTD | F | 49 | 3,4777474631 | 0,2793216771 | 12,4506894662 | 1,571 | 2 | - | 0,54 | 0,39 |
| 58 | CTD | LEAP Cohort | UCTD | M | 71 | 1,3016390133 | 101964,195537351 | 1,27656478473539E-005 | 0,837 | 2 | - | 0,80 | 0,91 |
| 63 | CTD | LEAP Cohort | MCTD | F | 47 | 41,6058901332 | 0,1387143064 | 299,9394309574 | 0,617 | 6 | - | 150,84 | 118,89 |
| 67 | CTD | LEAP Cohort | MCTD | M | 50 | 13,3091498343 | 5,4263195563 | 2,4527029225 | 3,732 | - | - | - | - |
| 68 | CTD | LEAP Cohort | UCTD | F | 28 | 0,0457736818 | 3,2388986243 | 0,0141324836 | 0,269 | 2 | - | 0,64 | 0,37 |
| 69 | CTD | LEAP Cohort | UCTD | F | 48 | 0,0316512951 | 86,9871183813 | 0,0003638619 | 0,158 | 3 | - | 3,51 | 2,78 |
| 72 | CTD | LEAP Cohort | UCTD | F | 60 | 0,0500324919 | 1,0643554386 | 0,0470073155 | 0,299 | 2 | - | 0,48 | 0,49 |
| 78 | CTD | LEAP Cohort | UCTD | F | 53 | 0,0091396821 | 205,427344745 | 4,44910684699703E-005 | 0,281 | - | - | 0,98 | 0,55 |
| 79 | CTD | LEAP Cohort | UCTD | F | 40 | 0,3612291893 | 505,6596132694 | 0,0007143722 | 0,314 | 2 | - | 0,70 | 0,55 |
| 81 | CTD | LEAP Cohort | UCTD | F | 49 | 0,4747955229 | 10847,7983702963 | 4,37688373876858E-005 | 0,439 | 3 | - | 0,37 | 0,36 |
| 83 | CTD | LEAP Cohort | UCTD | F | 34 | 0,117100946 | 9,3261713426 | 0,012556165 | 0,234 | 3 | - | 0,33 | 0,27 |
| 93 | CTD | LEAP Cohort | UCTD | F | 31 | 0,0939889503 | 1,7704622313 | 0,0530872383 | 0,12 | 3 | - | 0,41 | 0,50 |
| 95 | CTD | LEAP Cohort | UCTD | F | 51 | 373,6784292222 | 3869,2449502448 | 0,0965765761 | 15,818 | 2 | - | 5,25 | 3,62 |
| 96 | CTD | LEAP Cohort | UCTD | F | 52 | 1,2723819742 | 0,1387143064 | 9,1726802161 | 0,746 | 2 | - | 0,30 | 0,41 |
| 102 | CTD | LEAP Cohort | UCTD | F | 41 | 0,8409625081 | 0,7966570617 | 1,0556142016 | 0,187 | - | - | - | - |
| 108 | CTD | LEAP Cohort | UCTD | F | 63 | 2,0680900705 | 449,4899947445 | 0,0046009702 | 0,762 | - | - | - | - |
| 113 | CTD | LEAP Cohort | UCTD | F | 53 | 0,5299975963 | 363,577337397 | 0,00145773 | 0,323 | - | - | - | - |
| 114 | CTD | LEAP Cohort | UCTD | F | 57 | 0,006210424 | 23,9690773398 | 0,0002591015 | 0,209 | 2 | - | 0,45 | 0,63 |
| 124 | CTD | LEAP Cohort | UCTD | F | 41 | 0,3036280102 | 20039,617836656 | 1,51513872499364E-005 | 0,397 | 2 | - | 1,24 | 1,42 |
| 127 | CTD | LEAP Cohort | MCTD | M | 69 | 31,0003269072 | 46,0326517053 | 0,6734421277 | 10,836 | 2 | - | 0,84 | 0,58 |
| CTD002 | CTD | Rodero *et al.* (2017) | CTD | F | 44 | 501,4992167352 | 415,7746778012 | 1,2061802787 | - | - | - | - | - |
| CTD009 | CTD | Rodero *et al.* (2017) | CTD | F | 30 | 292,3958868316 | 222,4175413605 | 1,3146260184 | - | - | - | - | - |
| CTD014 | CTD | Rodero *et al.* (2017) | CTD | F | 40 | 111,1096191726 | 121,6037969903 | 0,9137018903 | - | - | - | - | - |
| CTD017 | CTD | Rodero *et al.* (2017) | CTD | F | 35 | 267,9669659894 | 190,1632800256 | 1,4091414807 | - | - | - | - | - |
| CTD021 | CTD | Rodero *et al.* (2017) | CTD | F | 29 | 1657,4684069455 | 2940,6614999938 | 0,5636379457 | - | - | - | - | - |
| CTD025 | CTD | Rodero *et al.* (2017) | CTD | F | 62 | 3,099495849 | 0,7164402454 | 4,326244748 | - | - | - | - | - |
| CTD026 | CTD | Rodero *et al.* (2017) | CTD | F | 49 | 328,9313436254 | 4,7172656686 | 69,7292386599 | - | - | - | - | - |
| CTD031 | CTD | Rodero *et al.* (2017) | CTD | F | 32 | 0,1354360775 | 3,2015540317 | 0,0423032303 | - | - | - | - | - |
| CTD033 | CTD | Rodero *et al.* (2017) | CTD | F | 58 | 236,8330233205 | 279,7493175116 | 0,8465901738 | - | - | - | - | - |
| CTD060 | CTD | Rodero *et al.* (2017) | CTD | F | 23 | 0,1076691437 | 0,7164402454 | 0,1502834945 | - | - | - | - | - |
| CTD061 | CTD | Rodero *et al.* (2017) | CTD | M | 17 | 67,6201078425 | 31,3866985063 | 2,1544192623 | - | - | - | - | - |
| CTD064 | CTD | Rodero *et al.* (2017) | CTD | F | 62 | 12,1071774868 | 55,059002073 | 0,2198946045 | - | - | - | - | - |
| CTD066 | CTD | Rodero *et al.* (2017) | CTD | F | 51 | 75,1476249235 | 21,7486966445 | 3,4552702698 | - | - | - | - | - |
| CTD067 | CTD | Rodero *et al.* (2017) | CTD | F | 52 | 0,2011365249 | 2,6062653831 | 0,0771742303 | - | - | - | - | - |
| CTD073 | CTD | Rodero *et al.* (2017) | CTD | F | 47 | 13,2274466646 | 548,9490472715 | 0,0240959461 | - | - | - | - | - |
| CTD074 | CTD | Rodero *et al.* (2017) | CTD | F | 45 | 1,8195291664 | 1,4619928063 | 1,2445541172 | - | - | - | - | - |
| CTD075 | CTD | Rodero *et al.* (2017) | CTD | F | 30 | 1035,8938603581 | 1035,0802824925 | 1,0007860046 | - | - | - | - | - |
| CTD096 | CTD | Rodero *et al.* (2017) | CTD | F | 51 | 2044,5242169973 | 3401,3242519923 | 0,6010965334 | - | - | - | - | - |
| CTD099 | CTD | Rodero *et al.* (2017) | CTD | F | 55 | 0,2997122793 | 2,5192283541 | 0,1189698738 | - | - | - | - | - |
| 29 | pSS | LEAP Cohort | pSS | F | 61 | 1,1167416279 | 37387,4046555558 | 2,98694610704913E-005 | 0,538 | 2 | - | 0,80 | 0,69 |
| 37 | pSS | LEAP Cohort | pSS | F | 57 | 0,8605962183 | 0,3656968797 | 2,3533047892 | 0,272 | - | - | - | - |
| 49 | pSS | LEAP Cohort | pSS | F | 58 | 13,1351825067 | 2236,4722146042 | 0,0058731704 | 3,426 | - | - | - | - |
| 57 | pSS | LEAP Cohort | pSS | F | 54 | 10,6404993791 | 756,0772436476 | 0,0140732967 | 5,485 | 2 | - | 1,20 | 1,14 |
| 60 | pSS | LEAP Cohort | pSS | F | 48 | 13,3542823385 | 5868,7108637669 | 0,0022755052 | 2,664 | - | - | - | - |
| 76 | pSS | LEAP Cohort | pSS | F | 33 | 263,8272827846 | 710,489458487 | 0,3713317342 | 16,473 | 2 | - | 0,45 | 0,30 |
| 80 | pSS | LEAP Cohort | pSS | F | 49 | 52,4055962974 | 67,7957149756 | 0,7729927521 | 10,774 | - | - | - | - |
| 82 | pSS | LEAP Cohort | pSS | F | 58 | 12,5184201265 | 200000 | 6,25921006324695E-005 | 3,558 | 2 | - | 0,58 | 0,90 |
| 99 | pSS | LEAP Cohort | pSS | F | 37 | 66,0637263222 | 4010,8847920841 | 0,0164711104 | 18,633 | 2 | - | 0,49 | 0,49 |
| 116 | pSS | LEAP Cohort | pSS | F | 46 | 0,0322010912 | 491,3320045929 | 6,55383547532449E-005 | 0,386 | 2 | - | 0,45 | 0,44 |
| 120 | pSS | LEAP Cohort | pSS | F | 55 | 0,0094973021 | 0,3656968797 | 0,0259704214 | 0,245 | 2 | - | 3,45 | 3,05 |
| 122 | pSS | LEAP Cohort | pSS | F | 62 | 268,1110827822 | 106389,072099404 | 0,0025200998 | 11,122 | - | - | - | - |
| 01-0001 | pSS | ASSESS Cohort | pSS | F | 59 | 6,4447290183 | 1,5822805628 | 4,0730633807 | 8,0019786739 | - | - | - | - |
| 01-0002 | pSS | ASSESS Cohort | pSS | F | 74 | 0,1014614551 | 1,0630154498 | 0,0954468301 | 6,1928334679 | - | - | - | - |

| | | | | | | | | | | | | | |
|---|---|---|---|---|---|---|---|---|---|---|---|---|---|
| 01-0003 | pSS | ASSESS Cohort | pSS | F | 56 | 62,0518223882 | 18,1765024124 | 3,413848329 | - | - | - | - | - |
| 01-0004 | pSS | ASSESS Cohort | pSS | F | 68 | 34,9734309347 | 16,7145073342 | 2,092399748 | 8,7382191735 | - | - | - | - |
| 01-0005 | pSS | ASSESS Cohort | pSS | F | 41 | 95,1180348452 | 19,8496846305 | 4,7919166786 | - | - | - | - | - |
| 01-0006 | pSS | ASSESS Cohort | pSS | F | 74 | 30,2267968543 | 0,3656968797 | 82,6553315985 | 6,5372324916 | - | - | - | - |
| 01-0007 | pSS | ASSESS Cohort | pSS | F | 42 | 52,5293964428 | 0,3656968797 | 143,6419049818 | 8,5895961128 | - | - | - | - |
| 01-0008 | pSS | ASSESS Cohort | pSS | F | 75 | 0,1183425522 | 0,3656968797 | 0,3236083181 | 5,3717228668 | - | - | - | - |
| 01-0009 | pSS | ASSESS Cohort | pSS | M | 57 | 0,0269867913 | 0,5655509296 | 0,047717703 | 6,5149271548 | - | - | - | - |
| 01-0010 | pSS | ASSESS Cohort | pSS | F | 49 | 22,7069147949 | 33,3263514072 | 0,6813501579 | - | - | - | - | - |
| 01-0011 | pSS | ASSESS Cohort | pSS | F | 74 | 362,8918498749 | 165039,271131048 | 0,0021988212 | 6,2345910311 | - | - | - | - |
| 01-0012 | pSS | ASSESS Cohort | pSS | F | 53 | 106,4998482396 | 170,188707358 | 0,6257750581 | 8,5989520135 | - | - | - | - |
| 01-0013 | pSS | ASSESS Cohort | pSS | F | 54 | 41,8132355488 | 159,9812058126 | 0,2613634229 | 8,875060525 | - | - | - | - |
| 01-0014 | pSS | ASSESS Cohort | pSS | F | 35 | 38,3364116452 | 22,7958195883 | 1,6817299109 | 8,1063398459 | - | - | - | - |
| 01-0015 | pSS | ASSESS Cohort | pSS | F | 61 | 18,619398626 | 0,8178873118 | 22,7652371644 | - | - | - | - | - |
| 01-0016 | pSS | ASSESS Cohort | pSS | F | 75 | 35,4630301328 | 13,6708850426 | 2,594055178 | 8,1093880585 | - | - | - | - |
| 01-0017 | pSS | ASSESS Cohort | pSS | F | 48 | 66,7467049639 | 486,2354494237 | 0,1372723956 | 9,3618496248 | - | - | - | - |
| 01-0018 | pSS | ASSESS Cohort | pSS | F | 37 | 27,1315039949 | 713,5710718007 | 0,0380221467 | - | - | - | - | - |
| 01-0019 | pSS | ASSESS Cohort | pSS | F | 71 | 93,3632649216 | 784,880652691 | 0,1189521803 | 6,7292405688 | - | - | - | - |
| 01-0020 | pSS | ASSESS Cohort | pSS | F | 59 | 8,8657625362 | 0,3656968797 | 24,2434732941 | 5,7451426161 | - | - | - | - |
| 01-0021 | pSS | ASSESS Cohort | pSS | F | 81 | 21,7412086085 | 20,2097019351 | 1,075780765 | 8,6444567484 | - | - | - | - |
| 01-0022 | pSS | ASSESS Cohort | pSS | F | 70 | 3,9722689466 | 1,0233080251 | 3,8817920404 | 9,182073602 | - | - | - | - |
| 01-0023 | pSS | ASSESS Cohort | pSS | F | 62 | 0,1744650745 | 2,6772070608 | 0,0651668214 | - | - | - | - | - |
| 01-0024 | pSS | ASSESS Cohort | pSS | F | 32 | 84,366050726 | 45,2622320832 | 1,8639392457 | 9,301829313 | - | - | - | - |
| 01-0025 | pSS | ASSESS Cohort | pSS | F | 72 | 11,7991714761 | 9,0519292618 | 1,3034979765 | 8,1621346471 | - | - | - | - |
| 01-0026 | pSS | ASSESS Cohort | pSS | M | 69 | 0,0842275924 | 0,7592079702 | 0,1109413965 | 6,5679420719 | - | - | - | - |
| 01-0027 | pSS | ASSESS Cohort | pSS | F | 66 | 5,6881449302 | 3,2548901828 | 1,7475689226 | 7,4190785508 | - | - | - | - |
| 01-0028 | pSS | ASSESS Cohort | pSS | F | 71 | 5,0132207265 | 33,5153867718 | 0,149579677 | - | - | - | - | - |
| 01-0029 | pSS | ASSESS Cohort | pSS | F | 70 | 13,5603178501 | 61,7260747313 | 0,219685407 | 7,3321061731 | - | - | - | - |
| 01-0030 | pSS | ASSESS Cohort | pSS | F | 45 | 41,130896899 | 483,7555450469 | 0,0850241353 | 9,1900834017 | - | - | - | - |
| 01-0031 | pSS | ASSESS Cohort | pSS | F | 50 | 31,5358889033 | 60,5972061253 | 0,5204181994 | 8,8112359186 | - | - | - | - |
| 01-0032 | pSS | ASSESS Cohort | pSS | M | 58 | 182,991194374 | 243,5762784077 | 0,7512685372 | 6,3831839474 | - | - | - | - |
| 01-0033 | pSS | ASSESS Cohort | pSS | F | 69 | 116,0209034231 | 323,7759805863 | 0,3583369687 | 6,5768863245 | - | - | - | - |
| 01-0034 | pSS | ASSESS Cohort | pSS | F | 68 | 31,6094019233 | 1,0011055571 | 31,5744945179 | 6,5433593811 | 2 | - | 0,60 | 0,46 |
| 01-0035 | pSS | ASSESS Cohort | pSS | F | 56 | 0,9597414484 | 29,5224020058 | 0,0325089215 | 6,5327852556 | - | - | - | - |
| 01-0036 | pSS | ASSESS Cohort | pSS | F | 56 | 8,1250206048 | 3,789525092 | 2,1440735732 | 7,7597322082 | - | - | - | - |
| 01-0037 | pSS | ASSESS Cohort | pSS | M | 71 | 13,4547166036 | 11,1608884513 | 1,2055237952 | 8,8385907951 | - | - | - | - |
| 01-0038 | pSS | ASSESS Cohort | pSS | M | 55 | 145,6748403401 | 241,5530717476 | 0,6030757174 | 10,2366047685 | 2 | - | 0,39 | 0,42 |
| 01-0039 | pSS | ASSESS Cohort | pSS | F | 54 | 157,006112699 | 225,1709202749 | 0,6972752632 | 8,7261697812 | - | - | - | - |
| 01-0040 | pSS | ASSESS Cohort | pSS | F | 61 | 41,0596161941 | 0,8451879805 | 48,5804544545 | 9,8997477605 | - | - | - | - |
| 01-0041 | pSS | ASSESS Cohort | pSS | F | 48 | 64,7266133356 | 50,3028651343 | 1,2867381045 | 9,7193038063 | - | - | - | - |
| 01-0042 | pSS | ASSESS Cohort | pSS | F | 78 | 0,4472835301 | 2,9484016675 | 0,1517037299 | - | - | - | - | - |
| 01-0043 | pSS | ASSESS Cohort | pSS | F | 74 | 12,7178596383 | 3,2739607376 | 3,8845486117 | 8,2117662554 | - | - | - | - |
| 01-0044 | pSS | ASSESS Cohort | pSS | F | 68 | 41,8533188425 | 472,786944926 | 0,0885246923 | 9,2636258469 | - | - | - | - |
| 01-0045 | pSS | ASSESS Cohort | pSS | F | 72 | 121,6918903369 | 247,5127871003 | 0,4916590038 | 10,0210768742 | - | - | - | - |
| 01-0046 | pSS | ASSESS Cohort | pSS | F | 81 | 77,1662762037 | 775,026554298 | 0,0995659772 | 9,4956544832 | - | - | - | - |
| 01-0047 | pSS | ASSESS Cohort | pSS | F | 64 | 0,0269867913 | 0,3656968797 | 0,0737955197 | - | - | - | - | - |
| 01-0048 | pSS | ASSESS Cohort | pSS | F | 84 | 33,418111981 | 21,3984923175 | 1,5617040437 | 8,6270184151 | - | - | - | - |
| 01-0049 | pSS | ASSESS Cohort | pSS | F | 39 | 27,7045558636 | 37,9285006826 | 0,73044163 | 8,9161884377 | 2 | - | 0,42 | 0,38 |
| 01-0050 | pSS | ASSESS Cohort | pSS | F | 66 | 0,0269867913 | 0,3656968797 | 0,0737955197 | 5,8209463783 | - | - | - | - |
| 01-0053 | pSS | ASSESS Cohort | pSS | F | 62 | 69,4153661777 | 879,3489095788 | 0,0789395033 | 7,4063980706 | - | - | - | - |
| 01-0054 | pSS | ASSESS Cohort | pSS | F | 65 | 29,171711906 | 991,9987809335 | 0,0294070038 | 7,6042669075 | - | - | - | - |
| 01-0055 | pSS | ASSESS Cohort | pSS | F | 30 | 330,1849259047 | 576,1240381044 | 0,5731143019 | 10,212230054 | - | - | - | - |
| 01-0056 | pSS | ASSESS Cohort | pSS | F | 80 | 36,6176718835 | 118754,777071964 | 0,0003083469 | 8,5062047449 | - | - | 0,82 | 0,89 |
| 01-0057 | pSS | ASSESS Cohort | pSS | F | 66 | 25,1273721239 | 3250,5634810351 | 0,0077301589 | 8,5533627969 | - | - | - | - |
| 01-0058 | pSS | ASSESS Cohort | pSS | F | 36 | 46,5203444559 | 41,7071740069 | 1,1154038979 | 9,0424796294 | - | - | - | - |
| 01-0059 | pSS | ASSESS Cohort | pSS | F | 62 | 27,5528899914 | 0,3682130897 | 74,8286542862 | 5,9202010484 | - | - | - | - |
| 01-0060 | pSS | ASSESS Cohort | pSS | F | 69 | 2,0921901989 | 271,1591316635 | 0,0077157283 | - | - | - | - | - |
| 01-0062 | pSS | ASSESS Cohort | pSS | F | 47 | 72,6427436107 | 33,2904564665 | 2,1820891427 | - | - | - | - | - |
| 01-0063 | pSS | ASSESS Cohort | pSS | F | 61 | 36,8305435877 | 53,1332712455 | 0,6931728976 | 6,854890614 | - | - | - | - |
| 01-0065 | pSS | ASSESS Cohort | pSS | F | 70 | 30,5576217859 | 61,7915503436 | 0,4945275141 | - | - | - | - | - |
| 01-0067 | pSS | ASSESS Cohort | pSS | M | 51 | 47,8223550189 | 8,4809246559 | 5,6388138038 | - | - | - | - | - |
| 01-0068 | pSS | ASSESS Cohort | pSS | M | 47 | 47,3018753393 | 0,3656968797 | 129,3472216143 | 5,9584272869 | - | - | - | - |
| 01-0071 | pSS | ASSESS Cohort | pSS | F | 70 | 4,085992676 | 2,909563566 | 1,4043318125 | - | - | - | - | - |
| 02-0004 | pSS | ASSESS Cohort | pSS | F | 73 | 31,3238512189 | 251,9674763424 | 0,1243170415 | 9,2003212936 | 2 | - | 0,48 | 0,28 |
| 02-0005 | pSS | ASSESS Cohort | pSS | F | 58 | 331,4002950631 | 582,1235337891 | 0,5692954774 | 10,519786918 | - | - | - | - |
| 02-0006 | pSS | ASSESS Cohort | pSS | F | 66 | 0,0269867913 | 0,3656968797 | 0,0737955197 | 6,2203920769 | - | - | - | - |
| 02-0008 | pSS | ASSESS Cohort | pSS | F | 36 | 211,6948178414 | 279,1899725544 | 0,758246494 | 10,1704386489 | 2 | - | 0,39 | 0,57 |
| 03-0001 | pSS | ASSESS Cohort | pSS | F | 31 | 41,6565940542 | 86,0157716866 | 0,4842901858 | 7,4133881378 | - | - | - | - |
| 03-0002 | pSS | ASSESS Cohort | pSS | F | 76 | 397,5192300269 | 120,8358090847 | 3,2897469139 | 7,0891668363 | - | - | - | - |
| 03-0003 | pSS | ASSESS Cohort | pSS | F | 38 | 14,237960536 | 15,0110881114 | 0,9484962336 | 9,8318954935 | - | - | - | - |
| 03-0004 | pSS | ASSESS Cohort | pSS | F | 54 | 0,0269867913 | 0,3656968797 | 0,0737955197 | 6,6112346312 | 2 | - | 1,30 | 1,28 |
| 03-0005 | pSS | ASSESS Cohort | pSS | F | 58 | 44,9332626263 | 0,3656968797 | 122,8702379575 | 7,2135303844 | - | - | - | - |
| 03-0006 | pSS | ASSESS Cohort | pSS | F | 69 | 7,7550651298 | 21,1973345568 | 0,3658509568 | 6,9573111556 | - | - | - | - |
| 03-0007 | pSS | ASSESS Cohort | pSS | M | 63 | 6,8426070364 | 44,9787266605 | 0,1521298521 | - | - | - | - | - |

| | | | | | | | | | | | | | |
|---|---|---|---|---|---|---|---|---|---|---|---|---|---|
| 03-0008 | pSS | ASSESS Cohort | pSS | F | 58 | 1,2195075654 | 2,8344727849 | 0,4302414092 | 7,0774882728 | - | - | - | - |
| 03-0009 | pSS | ASSESS Cohort | pSS | F | 56 | 0,30046685 | 0,5026956174 | 0,5977112982 | 7,2459075617 | - | - | - | - |
| 03-0010 | pSS | ASSESS Cohort | pSS | F | 54 | 1,4795722719 | 0,3656968797 | 4,045897994 | 6,8359163244 | - | - | - | - |
| 03-0011 | pSS | ASSESS Cohort | pSS | F | 48 | 95,6726377865 | 65,5607064849 | 1,4592984566 | 10,934228952 | - | - | - | - |
| 03-0012 | pSS | ASSESS Cohort | pSS | F | 33 | 113,585404945 | 139,6353875852 | 0,8134428307 | 9,6069248636 | - | - | - | - |
| 03-0013 | pSS | ASSESS Cohort | pSS | F | 60 | 167,6644141977 | 202,448847291 | 0,8281816194 | 10,1991618057 | - | - | - | - |
| 03-0014 | pSS | ASSESS Cohort | pSS | F | 56 | 48,2992019503 | 1426,2548945597 | 0,0338643549 | 9,536258809 | - | - | - | - |
| 03-0015 | pSS | ASSESS Cohort | pSS | F | 61 | 37,9346458132 | 15,5160653591 | 2,4448624658 | 6,8916034469 | - | - | - | - |
| 03-0016 | pSS | ASSESS Cohort | pSS | F | 44 | 115,7834754136 | 49,7366737587 | 2,3279296073 | 10,8192772324 | - | - | - | - |
| 03-0017 | pSS | ASSESS Cohort | pSS | F | 80 | 0,0357299572 | 1,1198953346 | 0,031904729 | 6,9642680595 | - | - | - | - |
| 03-0018 | pSS | ASSESS Cohort | pSS | F | 44 | 17,4977730909 | 2,0150679675 | 8,6834654578 | 6,6747021831 | - | - | - | - |
| 03-0019 | pSS | ASSESS Cohort | pSS | F | 57 | 0,1161047022 | 0,5024328254 | 0,2310850254 | 6,8202575604 | - | - | - | - |
| 03-0020 | pSS | ASSESS Cohort | pSS | F | 29 | 75,7191172599 | 56,0799038035 | 1,3502005554 | 9,4192522205 | - | - | - | - |
| 03-0021 | pSS | ASSESS Cohort | pSS | F | 57 | 12,9687719267 | 2019,1379109008 | 0,0064229253 | 7,6508163806 | - | - | - | - |
| 03-0022 | pSS | ASSESS Cohort | pSS | F | 45 | 3,795995635 | 1,4186357493 | 2,6758071175 | - | - | - | - | - |
| 03-0023 | pSS | ASSESS Cohort | pSS | F | 60 | 47,3276730009 | 31,1306257106 | 1,5202930208 | 8,4075208321 | - | - | - | - |
| 03-0024 | pSS | ASSESS Cohort | pSS | F | 66 | 2,5810821218 | 2,7537139307 | 0,9373094616 | - | - | - | - | - |
| 03-0025 | pSS | ASSESS Cohort | pSS | F | 71 | 34,9859093858 | 8203,6687755227 | 0,0042646663 | 6,4795905574 | - | - | - | - |
| 03-0026 | pSS | ASSESS Cohort | pSS | F | 60 | 1,468665597 | 0,3656968797 | 4,0160736355 | 7,056270238 | - | - | - | - |
| 03-0027 | pSS | ASSESS Cohort | pSS | F | 55 | 60,5636037476 | 15,2982605177 | 3,9588555625 | 9,3202332843 | - | - | - | - |
| 03-0028 | pSS | ASSESS Cohort | pSS | F | 51 | 391,4492702507 | 784,0479500658 | 0,499267003 | - | - | - | - | - |
| 03-0029 | pSS | ASSESS Cohort | pSS | F | 52 | 90,2630142338 | 806,779197205 | 0,1118806912 | 9,3368070116 | - | - | - | - |
| 03-0030 | pSS | ASSESS Cohort | pSS | F | 58 | 85,4592729163 | 237,8283498834 | 0,3593317321 | - | - | - | - | - |
| 03-0031 | pSS | ASSESS Cohort | pSS | F | 67 | 27,8625391876 | 123,0179561728 | 0,2264916444 | 6,980416434 | 2 | - | 0,47 | 0,64 |
| 03-0032 | pSS | ASSESS Cohort | pSS | F | 40 | 0,0269867913 | 0,9236895926 | 0,029216299 | 6,5614322118 | - | - | - | - |
| 03-0033 | pSS | ASSESS Cohort | pSS | F | 59 | 9,1284662092 | 0,3656968797 | 24,9618378402 | 7,5903106735 | - | - | - | - |
| 03-0034 | pSS | ASSESS Cohort | pSS | F | 54 | 0,7403552145 | 0,3656968797 | 2,0245051452 | 6,8705669831 | - | - | - | - |
| 03-0035 | pSS | ASSESS Cohort | pSS | F | 60 | 59,9282230782 | 105,5760747096 | 0,5676307179 | - | - | - | - | - |
| 03-0036 | pSS | ASSESS Cohort | pSS | F | 47 | 45,7461022912 | 125,30640738 | 0,3650739276 | 8,8312731338 | - | - | - | - |
| 03-0037 | pSS | ASSESS Cohort | pSS | F | 44 | 69,6656737996 | 6,8289331598 | 10,2015457128 | 9,99326338 | - | - | - | - |
| 03-0038 | pSS | ASSESS Cohort | pSS | F | 51 | 23,5439738431 | 0,3656968797 | 64,381117673 | - | - | - | - | - |
| 03-0039 | pSS | ASSESS Cohort | pSS | F | 50 | 167,7230175072 | 2032,8117378741 | 0,082507895 | 9,9041169643 | - | - | - | - |
| 03-0040 | pSS | ASSESS Cohort | pSS | F | 70 | 146,446672426 | 2678,7550732996 | 0,0546696762 | 9,6199830876 | - | - | - | - |
| 03-0041 | pSS | ASSESS Cohort | pSS | F | 36 | 176,5435225736 | 129,1102075403 | 1,3673862504 | - | - | - | - | - |
| 04-0002 | pSS | ASSESS Cohort | pSS | F | 30 | 0,370327506 | 0,3656968797 | 1,0126624715 | 7,8433492302 | - | - | - | - |
| 04-0003 | pSS | ASSESS Cohort | pSS | M | 51 | 0,41376523 | 0,3656968797 | 1,1314431514 | 6,0420922731 | - | - | - | - |
| 04-0004 | pSS | ASSESS Cohort | pSS | F | 54 | 0,058328619 | 0,3656968797 | 0,159499909 | 6,2084429483 | - | - | - | - |
| 04-0005 | pSS | ASSESS Cohort | pSS | F | 71 | 56,0803447437 | 0,3656968797 | 153,35199139 | 6,4774432838 | - | - | - | - |
| 04-0006 | pSS | ASSESS Cohort | pSS | F | 78 | 8,3846015948 | 14549676,1584305 | 5,7627410421154E-007 | 8,1734742029 | - | - | - | - |
| 04-0007 | pSS | ASSESS Cohort | pSS | F | 64 | 40,093614078 | 0,3656968797 | 109,6361940889 | 9,1542695786 | 2 | - | 0,49 | 0,47 |
| 04-0008 | pSS | ASSESS Cohort | pSS | F | 59 | 0,0869592972 | 0,3656968797 | 0,2377906459 | 6,2816769132 | - | - | - | - |
| 05-0001 | pSS | ASSESS Cohort | pSS | F | 39 | 7,3093629512 | 0,3656968797 | 19,987490617 | 7,80811932 | - | - | - | - |
| 05-0002 | pSS | ASSESS Cohort | pSS | F | 45 | 36,8797093074 | 0,3656968797 | 100,8477549492 | 8,73478485 | - | - | - | - |
| 05-0003 | pSS | ASSESS Cohort | pSS | F | 46 | 246,5988254986 | 316,056404765 | 0,7802367608 | 10,2414555239 | 2 | - | 0,34 | 0,38 |
| 05-0004 | pSS | ASSESS Cohort | pSS | F | 41 | 0,1321721478 | 0,3656968797 | 0,3614254186 | 6,0175720008 | 2 | - | 0,67 | 0,58 |
| 05-0005 | pSS | ASSESS Cohort | pSS | F | 59 | 0,0606440878 | 8629,9929522276 | 7,02713062928976E-006 | 6,375584379 | 2 | - | 0,49 | 0,46 |
| 05-0006 | pSS | ASSESS Cohort | pSS | F | 64 | 0,0203594309 | 0,3656968797 | 0,0556729685 | 6,3832736467 | - | - | - | - |
| 05-0008 | pSS | ASSESS Cohort | pSS | F | 53 | 14,5286861236 | 20,5108587518 | 0,7083411913 | 8,0651032487 | 2 | - | 0,43 | 0,48 |
| 05-0010 | pSS | ASSESS Cohort | pSS | F | 53 | 202,2532169819 | 196,631710506 | 1,0285890127 | 10,1457131053 | - | - | - | - |
| 05-0011 | pSS | ASSESS Cohort | pSS | F | 53 | 9,1375446104 | 0,3656968797 | 24,9866627749 | 7,0109775653 | - | - | - | - |
| 05-0013 | pSS | ASSESS Cohort | pSS | F | 61 | 24,0903427088 | 0,3656968797 | 65,8751661487 | 8,3319940978 | 3 | - | 0,45 | 0,33 |
| 05-0014 | pSS | ASSESS Cohort | pSS | F | 53 | 48,044476276 | 13,9977827415 | 3,4322918967 | 8,7368520611 | - | - | - | - |
| 05-0015 | pSS | ASSESS Cohort | pSS | F | 51 | 15,733118964 | 0,3656968797 | 43,0222947974 | 7,950596577 | 3 | - | 0,51 | 0,51 |
| 05-0016 | pSS | ASSESS Cohort | pSS | F | 39 | 298,6084085409 | 293,8408899988 | 1,0162248302 | 10,2307386852 | - | - | - | - |
| 05-0019 | pSS | ASSESS Cohort | pSS | F | 60 | 39,1974415818 | 137,0386654845 | 0,2860319855 | 9,0367404702 | 2 | - | 0,56 | 0,33 |
| 05-0020 | pSS | ASSESS Cohort | pSS | F | 52 | 33,7730534946 | 323,8763085969 | 0,1042776288 | 9,1590978553 | - | - | - | - |
| 05-0021 | pSS | ASSESS Cohort | pSS | F | 50 | 0,0072302156 | 0,3656968797 | 0,0197710619 | 6,2526610117 | - | - | - | - |
| 05-0022 | pSS | ASSESS Cohort | pSS | F | 72 | 29,1456778285 | 0,3656968797 | 79,6990060572 | 8,1989665947 | - | - | - | - |
| 05-0023 | pSS | ASSESS Cohort | pSS | F | 54 | 2,5892893636 | 1,1081723627 | 2,3365402809 | 6,6899446843 | - | - | - | - |
| 05-0024 | pSS | ASSESS Cohort | pSS | F | 25 | 328,9016794858 | 498,2401486577 | 0,6601268091 | 11,152099013 | - | - | - | - |
| 05-0025 | pSS | ASSESS Cohort | pSS | F | 56 | 52,1726908432 | 492,1500627232 | 0,106009721 | 6,5452937368 | - | - | - | - |
| 05-0026 | pSS | ASSESS Cohort | pSS | F | 62 | 79,9282250285 | 22,7894056094 | 3,5072536072 | - | - | - | - | - |
| 05-0027 | pSS | ASSESS Cohort | pSS | F | 80 | 6,5708989812 | 649,9908279086 | 0,010109218 | 7,3652543725 | - | - | - | - |
| 05-0028 | pSS | ASSESS Cohort | pSS | F | 70 | 0,0072302156 | 0,3656968797 | 0,0197710619 | 6,3009614352 | - | - | - | - |
| 05-0030 | pSS | ASSESS Cohort | pSS | F | 48 | 324,297456846 | 342,3239033656 | 0,9473409647 | 10,286607549 | - | - | - | - |
| 05-0031 | pSS | ASSESS Cohort | pSS | F | 55 | 0,3821248648 | 0,3656968797 | 1,0449224098 | 6,3137763522 | - | - | - | - |
| 05-0032 | pSS | ASSESS Cohort | pSS | F | 59 | 0,0184132067 | 0,3656968797 | 0,0503510084 | 6,4262518738 | - | - | - | - |
| 05-0033 | pSS | ASSESS Cohort | pSS | M | 67 | 0,494799106 | 0,3656968797 | 1,3530307025 | 6,2958736768 | - | - | - | - |
| 05-0034 | pSS | ASSESS Cohort | pSS | M | 53 | 0,0130643016 | 0,3656968797 | 0,0357244 | 6,99387676 | - | - | - | - |
| 06-0001 | pSS | ASSESS Cohort | pSS | F | 64 | 53,1972551746 | 45,6160221539 | 1,1661967147 | 8,3253295636 | - | - | - | - |
| 06-0002 | pSS | ASSESS Cohort | pSS | F | 59 | 1,0034673546 | 0,3656968797 | 2,7439866469 | 7,311236997 | - | - | - | - |
| 06-0003 | pSS | ASSESS Cohort | pSS | F | 56 | 2,2467316921 | 0,3656968797 | 6,1436993778 | 6,7339460773 | - | - | - | - |
| 06-0004 | pSS | ASSESS Cohort | pSS | F | 59 | 108,5805176486 | 2014,4877490672 | 0,0538998153 | - | - | - | - | - |

| | | | | | | | | | | | | | |
|---|---|---|---|---|---|---|---|---|---|---|---|---|---|
| 06-0005 | pSS | ASSESS Cohort | pSS | F | 49 | 34,3255826957 | 17,4884492147 | 1,9627573763 | 8,7167253435 | - | - | - | - |
| 06-0006 | pSS | ASSESS Cohort | pSS | F | 47 | 91,6813572212 | 23,292450488 | 3,9360975467 | 7,1541481271 | - | - | - | - |
| 06-0008 | pSS | ASSESS Cohort | pSS | F | 61 | 37,8996798433 | 0,3656968797 | 103,6368696303 | - | - | - | - | - |
| 06-0009 | pSS | ASSESS Cohort | pSS | F | 50 | 0,5573099294 | 0,3656968797 | 1,5239668708 | 6,69708901 | - | - | - | - |
| 06-0010 | pSS | ASSESS Cohort | pSS | F | 47 | 40,3000296806 | 0,3656968797 | 110,2006386166 | 7,6292706539 | - | - | - | - |
| 07-0001 | pSS | ASSESS Cohort | pSS | F | 75 | 138,9570059431 | 96,909475687 | 1,433884612 | 9,5869145552 | - | - | - | - |
| 07-0002 | pSS | ASSESS Cohort | pSS | F | 53 | 53,6781847807 | 0,3656968797 | 146,7832726056 | 8,6990585031 | 3 | - | 0,57 | 0,51 |
| 07-0003 | pSS | ASSESS Cohort | pSS | F | 54 | 0,1890772507 | 0,3656968797 | 0,5170327153 | 5,7466439017 | - | - | - | - |
| 07-0004 | pSS | ASSESS Cohort | pSS | F | 75 | 271,6965698839 | 6151,9461128475 | 0,0441643286 | 10,093961326 | - | - | - | - |
| 07-0005 | pSS | ASSESS Cohort | pSS | F | 71 | 2,6675066029 | 0,3656968797 | 7,2943105374 | 8,3287650231 | - | - | - | - |
| 07-0006 | pSS | ASSESS Cohort | pSS | F | 44 | 0,5472185247 | 0,3656968797 | 1,4963718727 | 6,4826080079 | - | - | - | - |
| 07-0007 | pSS | ASSESS Cohort | pSS | F | 65 | 0,8235254614 | 0,3656968797 | 2,2519346135 | 6,6731768793 | - | - | - | - |
| 07-0008 | pSS | ASSESS Cohort | pSS | F | 50 | 736,0928440519 | 1618,6649006489 | 0,4547530769 | 10,9747908072 | 9 | - | 0,32 | 0,25 |
| 07-0009 | pSS | ASSESS Cohort | pSS | F | 80 | 149,4384571478 | 0,3656968797 | 408,6402303449 | 7,6117403451 | 2 | - | 0,37 | 0,44 |
| 07-0010 | pSS | ASSESS Cohort | pSS | F | 68 | 60,065688351 | 12713,4799346433 | 0,004724567 | 8,5249333579 | - | - | - | - |
| 07-0011 | pSS | ASSESS Cohort | pSS | F | 58 | 16,416043011 | 0,3656968797 | 44,8897541195 | 8,3559094436 | 2 | - | 0,98 | 0,35 |
| 07-0012 | pSS | ASSESS Cohort | pSS | F | 53 | 65,221002029 | 165,5011795138 | 0,3940817958 | 9,6308854199 | - | - | - | - |
| 07-0013 | pSS | ASSESS Cohort | pSS | F | 41 | 131,9420446833 | 44,2708926982 | 2,980333954 | 9,6864531178 | - | - | - | - |
| 07-0014 | pSS | ASSESS Cohort | pSS | F | 38 | 163,3166843705 | 243,5021030279 | 0,6706992767 | 9,4497635453 | 3 | - | 0,30 | 0,25 |
| 07-0015 | pSS | ASSESS Cohort | pSS | F | 60 | 6,4354650502 | 0,3656968797 | 17,5978123081 | 8,092275599 | - | - | - | - |
| 07-0016 | pSS | ASSESS Cohort | pSS | F | 57 | 85,6286168012 | 271,4561447882 | 0,3154418069 | 7,0048060851 | 2 | - | 2,96 | 2,10 |
| 07-0017 | pSS | ASSESS Cohort | pSS | F | 44 | 746,9622882118 | 0,3656968797 | 2042,5722223024 | 8,4532382391 | 2 | - | 1,11 | 0,76 |
| 07-0018 | pSS | ASSESS Cohort | pSS | M | 73 | 2,9332197439 | 0,3656968797 | 8,0209044891 | 7,7492144504 | - | - | - | - |
| 07-0019 | pSS | ASSESS Cohort | pSS | F | 74 | 3,683027359 | 0,3656968797 | 10,0712572724 | 7,6955766777 | - | - | - | - |
| 07-0020 | pSS | ASSESS Cohort | pSS | F | 54 | 154,8250348258 | 1551,4036200419 | 0,0997967472 | 9,1493641191 | - | - | - | - |
| 07-0021 | pSS | ASSESS Cohort | pSS | F | 66 | 0,016617732 | 0,3656968797 | 0,0454412738 | 6,8462436162 | - | - | - | - |
| 07-0022 | pSS | ASSESS Cohort | pSS | F | 59 | 22,8936589287 | 23963,0567092533 | 0,0009553731 | 8,3779421876 | 2 | - | 0,49 | 0,52 |
| 07-0023 | pSS | ASSESS Cohort | pSS | F | 54 | 262,7414595813 | 280,9042738979 | 0,9353416234 | 10,1174095247 | 2 | - | 0,46 | 0,37 |
| 07-0024 | pSS | ASSESS Cohort | pSS | F | 65 | 1,8544431144 | 0,3656968797 | 5,0709842427 | 6,9627784315 | - | - | - | - |
| 07-0025 | pSS | ASSESS Cohort | pSS | F | 60 | 101,4312287468 | 1749,40948492 | 0,0579802668 | 9,566523634 | - | - | - | - |
| 07-0026 | pSS | ASSESS Cohort | pSS | F | 68 | 233,6847167839 | 599,534103268 | 0,3897771878 | 10,3542363939 | - | - | - | - |
| 07-0027 | pSS | ASSESS Cohort | pSS | F | 56 | 0,0364379303 | 0,3656968797 | 0,0996397081 | 5,8498114697 | - | - | - | - |
| 07-0028 | pSS | ASSESS Cohort | pSS | F | 67 | 40,4315614537 | 2604,7418756519 | 0,015522291 | 8,1931505926 | - | - | - | - |
| 07-0029 | pSS | ASSESS Cohort | pSS | F | 47 | 97,6631405366 | 0,3656968797 | 267,0603605443 | 6,3636220715 | 2 | - | 0,42 | 0,48 |
| 07-0030 | pSS | ASSESS Cohort | pSS | F | 60 | 3,4207778011 | 0,3656968797 | 9,3541345065 | 7,5484150108 | - | - | - | - |
| 07-0031 | pSS | ASSESS Cohort | pSS | F | 30 | 0,111349184 | 0,3656968797 | 0,3044849169 | 6,0887357215 | 2 | - | 0,37 | 0,40 |
| 07-0032 | pSS | ASSESS Cohort | pSS | F | 43 | 97,1983351244 | 19,5001872142 | 4,9844821517 | 9,6019611461 | - | - | - | - |
| 07-0033 | pSS | ASSESS Cohort | pSS | F | 57 | 53,0941418518 | 32,1806044663 | 1,6498801913 | 8,385474624 | - | - | - | - |
| 07-0034 | pSS | ASSESS Cohort | pSS | F | 54 | 1,1440849037 | 275,4725498405 | 0,0041531721 | 6,4529157843 | - | - | - | - |
| 07-0035 | pSS | ASSESS Cohort | pSS | F | 61 | 189,5637755734 | 324,3364139031 | 0,5844665213 | 8,7830931284 | - | - | - | - |
| 07-0036 | pSS | ASSESS Cohort | pSS | F | 69 | 528,0123534349 | 9037,9780079224 | 0,0584215134 | 9,9170032607 | - | - | - | - |
| 07-0037 | pSS | ASSESS Cohort | pSS | F | 34 | 74,722655861 | 409,1279830214 | 0,1826388293 | 9,0580015714 | 3 | - | 0,39 | 0,41 |
| 07-0038 | pSS | ASSESS Cohort | pSS | F | 60 | 0,3235770249 | 0,3656968797 | 0,8848230402 | 6,1070000855 | 2 | - | 0,46 | 0,47 |
| 07-0039 | pSS | ASSESS Cohort | pSS | F | 65 | 22,7201481976 | 49,7532072081 | 0,4566569569 | 8,1912118505 | - | - | - | - |
| 07-0040 | pSS | ASSESS Cohort | pSS | F | 64 | 12,3674395786 | 0,3656968797 | 33,8188271923 | 6,1182669821 | - | - | - | - |
| 07-0041 | pSS | ASSESS Cohort | pSS | F | 53 | 14,675290607 | 0,3656968797 | 40,1296577097 | - | - | - | - | - |
| 07-0042 | pSS | ASSESS Cohort | pSS | F | 27 | 426,3366077507 | 1746,5199971543 | 0,244106342 | 9,8220155152 | 5 | - | 0,58 | 0,48 |
| 07-0043 | pSS | ASSESS Cohort | pSS | F | 73 | 0,0791611091 | 0,3656968797 | 0,2164664603 | 6,7068777297 | - | - | - | - |
| 07-0044 | pSS | ASSESS Cohort | pSS | F | 33 | 45,5304476251 | 142,5851200077 | 0,319321172 | - | - | - | - | - |
| 07-0045 | pSS | ASSESS Cohort | pSS | M | 27 | 199,0188825274 | 144,5388397855 | 1,376923205 | 9,684163047 | - | - | - | - |
| 08-0001 | pSS | ASSESS Cohort | pSS | F | 64 | 0,1289508405 | 0,3656968797 | 0,3526167372 | 6,7325684304 | - | - | - | - |
| 08-0002 | pSS | ASSESS Cohort | pSS | F | 61 | 0,0504837085 | 0,3656968797 | 0,1380479608 | 6,3877138212 | - | - | - | - |
| 08-0003 | pSS | ASSESS Cohort | pSS | F | 61 | 4,0358308046 | 727,0515548153 | 0,0055509555 | 6,1556791674 | - | - | - | - |
| 08-0004 | pSS | ASSESS Cohort | pSS | M | 68 | 0,1894578518 | 0,3656968797 | 0,5180734711 | 6,472958812 | - | - | - | - |
| 08-0005 | pSS | ASSESS Cohort | pSS | F | 42 | 280,745137559 | 2127,3478249191 | 0,1319695511 | 8,8758131343 | - | - | - | - |
| 08-0006 | pSS | ASSESS Cohort | pSS | F | 68 | 67,0123105074 | 0,3656968797 | 183,2455080462 | 9,8005004544 | - | - | - | - |
| 08-0007 | pSS | ASSESS Cohort | pSS | F | 68 | 77,3916818589 | 0,3656968797 | 211,6279524375 | 7,4071876769 | - | - | - | - |
| 08-0008 | pSS | ASSESS Cohort | pSS | M | 56 | 36,4561558929 | 11,7589026111 | 3,1003025621 | 8,9404627298 | - | - | - | - |
| 08-0009 | pSS | ASSESS Cohort | pSS | F | 56 | 34,364490759 | 1140,3915953214 | 0,030133939 | 8,9602722009 | - | - | - | - |
| 08-0010 | pSS | ASSESS Cohort | pSS | F | 52 | 195,2615003226 | 93,2587496544 | 2,0937606503 | 9,6370501107 | - | - | - | - |
| 08-0013 | pSS | ASSESS Cohort | pSS | F | 59 | 60,3287321381 | 335,6507854454 | 0,1797366035 | 8,3089911106 | - | - | - | - |
| 08-0014 | pSS | ASSESS Cohort | pSS | F | 71 | 297,0551279336 | 2192,4025980413 | 0,1354929647 | 9,5865810964 | - | - | - | - |
| 08-0016 | pSS | ASSESS Cohort | pSS | F | 78 | 172,9296172254 | 79,7269712198 | 2,1690227859 | 9,7887949279 | - | - | - | - |
| 08-0018 | pSS | ASSESS Cohort | pSS | F | 76 | 38,1287677283 | 46,5697309282 | 0,8187457168 | 9,0001123294 | - | - | - | - |
| 08-0021 | pSS | ASSESS Cohort | pSS | F | 61 | 74,4300814055 | 37,4875642455 | 1,9854605895 | 8,7063529638 | - | - | - | - |
| 08-0022 | pSS | ASSESS Cohort | pSS | F | 52 | 42,1911767985 | 0,3656968797 | 115,3719901458 | 8,5167648655 | - | - | - | - |
| 09-0001 | pSS | ASSESS Cohort | pSS | F | 71 | 0,0504837085 | 0,3656968797 | 0,1380479608 | 6,0321157526 | - | - | - | - |
| 09-0002 | pSS | ASSESS Cohort | pSS | F | 59 | 177,8978575214 | 353,3127132535 | 0,5035138868 | 10,2029596562 | 2 | - | 0,59 | 1,32 |
| 09-0003 | pSS | ASSESS Cohort | pSS | F | 68 | 0,4065611985 | 133,6179404372 | 0,0030427142 | 6,3913851824 | - | - | - | - |
| 09-0004 | pSS | ASSESS Cohort | pSS | M | 66 | 50,2197523955 | 1842,2630311502 | 0,0272598166 | 6,9576027865 | - | - | - | - |
| 09-0005 | pSS | ASSESS Cohort | pSS | F | 58 | 0,0504837085 | 0,3656968797 | 0,1380479608 | 6,2840706193 | - | - | - | - |
| 09-0007 | pSS | ASSESS Cohort | pSS | F | 67 | 0,0504837085 | 0,3656968797 | 0,1380479608 | 6,5098345226 | - | - | - | - |
| 09-0008 | pSS | ASSESS Cohort | pSS | F | 37 | 410,7787601674 | 3681,1742810818 | 0,1115890552 | 10,321610573 | - | - | - | - |

| | | | | | | | | | | | | | |
|---|---|---|---|---|---|---|---|---|---|---|---|---|---|
| 09-0009 | pSS | ASSESS Cohort | pSS | F | 56 | 5,4207694231 | 0,3656968797 | 14,8231218924 | 6,2416795988 | - | - | - | - |
| 09-0010 | pSS | ASSESS Cohort | pSS | F | 48 | 71,7958422431 | 0,9903285025 | 72,4969967627 | 9,0823874257 | 2 | - | 0,42 | 0,41 |
| 09-0011 | pSS | ASSESS Cohort | pSS | F | 58 | 86,0879136281 | 114,6408316981 | 0,7509358782 | 8,8164097534 | 3 | - | 0,43 | 0,36 |
| 09-0012 | pSS | ASSESS Cohort | pSS | F | 61 | 45,5570269924 | 0,3656968797 | 124,5759248276 | 8,4970000115 | 2 | - | 0,32 | 0,50 |
| 09-0013 | pSS | ASSESS Cohort | pSS | F | 51 | 1659,4934826279 | 3492,5491346159 | 0,4751525086 | 10,8089162669 | 12 | - | 0,48 | 0,42 |
| 09-0014 | pSS | ASSESS Cohort | pSS | F | 50 | 13,5034315745 | 0,3656968797 | 36,9252031529 | 7,6010545661 | 2 | - | 1,27 | 1,20 |
| 09-0015 | pSS | ASSESS Cohort | pSS | F | 33 | 205,5018000144 | 151,1376225013 | 1,3596998326 | 10,177569875 | - | - | - | - |
| 09-0016 | pSS | ASSESS Cohort | pSS | F | 59 | 0,1258991301 | 0,3656968797 | 0,3442718194 | 6,4068190086 | - | - | - | - |
| 09-0018 | pSS | ASSESS Cohort | pSS | F | 56 | 0,3897787455 | 0,3656968797 | 1,065851986 | 6,4533211236 | - | - | - | - |
| 09-0019 | pSS | ASSESS Cohort | pSS | F | 61 | 108,0746351726 | 875,7048691857 | 0,1234144504 | 7,4106492282 | - | - | - | - |
| 09-0020 | pSS | ASSESS Cohort | pSS | F | 62 | 71,6756487841 | 2793,6016240549 | 0,0256570759 | 6,0968392429 | - | - | - | - |
| 09-0021 | pSS | ASSESS Cohort | pSS | F | 53 | 0,1814169058 | 0,3656968797 | 0,4960854627 | 6,4808351704 | - | - | - | - |
| 10-0001 | pSS | ASSESS Cohort | pSS | F | 45 | 98,415864614 | 0,3656968797 | 269,1186884089 | 7,153446394 | - | - | - | - |
| 10-0002 | pSS | ASSESS Cohort | pSS | F | 38 | 0,1774907807 | 0,3656968797 | 0,4853494536 | 6,0571300791 | - | - | - | - |
| 10-0003 | pSS | ASSESS Cohort | pSS | F | 49 | 37,4457180088 | 784,2537003153 | 0,0477469446 | - | - | - | - | - |
| 10-0004 | pSS | ASSESS Cohort | pSS | M | 79 | 19,9323457033 | 0,3656968797 | 54,5051019327 | 7,0919449532 | - | - | - | - |
| 10-0005 | pSS | ASSESS Cohort | pSS | F | 67 | 72,0026819809 | 0,3656968797 | 196,8917045296 | 8,6199940245 | - | - | - | - |
| 10-0006 | pSS | ASSESS Cohort | pSS | F | 74 | 0,6573013343 | 334,0012384272 | 0,0019679608 | 6,5519224357 | - | - | - | - |
| 10-0007 | pSS | ASSESS Cohort | pSS | F | 72 | 2,0477538913 | 0,3656968797 | 5,5995935572 | 6,877632442 | - | - | - | - |
| 10-0008 | pSS | ASSESS Cohort | pSS | F | 75 | 4,2000052867 | 0,3656968797 | 11,4849360771 | 7,2249097686 | - | - | - | - |
| 10-0009 | pSS | ASSESS Cohort | pSS | F | 74 | 147,6423794272 | 397,4293720578 | 0,3714933767 | 8,2231904861 | - | - | - | - |
| 10-0010 | pSS | ASSESS Cohort | pSS | F | 59 | 0,0892456472 | 0,3656968797 | 0,2440426816 | 6,4291418007 | - | - | - | - |
| 10-0011 | pSS | ASSESS Cohort | pSS | F | 57 | 115,3475886186 | 39,428632273 | 2,9254778056 | 10,0568629648 | - | - | - | - |
| 10-0012 | pSS | ASSESS Cohort | pSS | F | 72 | 30,7424856827 | 0,3656968797 | 84,0654853546 | 7,318739017 | - | - | - | - |
| 10-0013 | pSS | ASSESS Cohort | pSS | M | 60 | 54,604277908 | 100,0903309977 | 0,5455499783 | 8,492024632 | - | - | - | - |
| 10-0014 | pSS | ASSESS Cohort | pSS | F | 62 | 133,6679098366 | 0,3656968797 | 365,5155875393 | 6,9264617228 | - | - | - | - |
| 10-0015 | pSS | ASSESS Cohort | pSS | F | 61 | 579,8112911166 | 4245,4529236008 | 0,1365723049 | 10,6668529082 | - | - | - | - |
| 10-0016 | pSS | ASSESS Cohort | pSS | F | 71 | 43,6792372583 | 0,3656968797 | 119,4410991334 | 7,4217929733 | - | - | - | - |
| 10-0017 | pSS | ASSESS Cohort | pSS | F | 58 | 56,5055702125 | 0,3656968797 | 154,5147726236 | 8,0697975185 | - | - | - | - |
| 10-0018 | pSS | ASSESS Cohort | pSS | F | 78 | 2,9692430971 | 0,3656968797 | 8,1194105339 | 7,4774682139 | - | - | - | - |
| 10-0019 | pSS | ASSESS Cohort | pSS | F | 33 | 71,5022236559 | 11,6578995139 | 6,1333710735 | 8,9116991273 | - | - | - | - |
| 10-0020 | pSS | ASSESS Cohort | pSS | F | 61 | 64,115424232 | 264,7260071426 | 0,2421954115 | 6,2175521031 | - | - | - | - |
| 10-0021 | pSS | ASSESS Cohort | pSS | M | 70 | 75,9277521716 | 308,7655431335 | 0,2459074656 | 8,7513080738 | - | - | - | - |
| 10-0022 | pSS | ASSESS Cohort | pSS | F | 40 | 66,8434655323 | 112,3243719353 | 0,5950931608 | 9,0194575845 | - | - | - | - |
| 10-0023 | pSS | ASSESS Cohort | pSS | F | 79 | 2,279313836 | 2,6246805137 | 0,8684157268 | 7,8709338364 | - | - | - | - |
| 10-0024 | pSS | ASSESS Cohort | pSS | F | 63 | 0,2409369535 | 0,3656968797 | 0,6588433395 | - | - | - | - | - |
| 10-0025 | pSS | ASSESS Cohort | pSS | F | 64 | 20,8523975925 | 41,779473548 | 0,4991062793 | 7,6109649532 | - | - | - | - |
| 10-0026 | pSS | ASSESS Cohort | pSS | F | 55 | 6,7533573264 | 0,3656968797 | 18,4670903738 | 7,8465966891 | - | - | - | - |
| 10-0027 | pSS | ASSESS Cohort | pSS | F | 68 | 60,8484970718 | 985,4358949139 | 0,0617477985 | - | - | - | - | - |
| 10-0028 | pSS | ASSESS Cohort | pSS | F | 47 | 27,5832995797 | 0,3656968797 | 75,4266746929 | 8,7848300607 | - | - | - | - |
| 10-0029 | pSS | ASSESS Cohort | pSS | F | 37 | 81,9706602053 | 0,3656968797 | 224,1491922972 | - | - | - | - | - |
| 10-0030 | pSS | ASSESS Cohort | pSS | F | 61 | 56,8792398291 | 1197,3328358757 | 0,0475049528 | 9,210214233 | - | - | - | - |
| 10-0031 | pSS | ASSESS Cohort | pSS | F | 52 | 23,0682082064 | 0,3656968797 | 63,0801340902 | 8,3819863175 | - | - | - | - |
| 10-0032 | pSS | ASSESS Cohort | pSS | M | 52 | 0,0896710591 | 0,3656968797 | 0,2452059726 | 6,5215173224 | 2 | - | 0,35 | 0,35 |
| 10-0033 | pSS | ASSESS Cohort | pSS | M | 71 | 909,3151646076 | 1211,612441414 | 0,7505000226 | - | - | - | - | - |
| 11-0001 | pSS | ASSESS Cohort | pSS | F | 77 | 0,2889868134 | 0,3656968797 | 0,7902359289 | 6,8566829517 | - | - | - | - |
| 11-0002 | pSS | ASSESS Cohort | pSS | F | 71 | 11,4004854102 | 3121,7861799535 | 0,003651911 | 7,8043023858 | - | - | - | - |
| 11-0003 | pSS | ASSESS Cohort | pSS | F | 52 | 18,8445092382 | 0,3656968797 | 51,530407519 | 8,737642857 | - | - | - | - |
| 11-0004 | pSS | ASSESS Cohort | pSS | F | 55 | 272,8268502083 | 177,2796866178 | 1,5389628412 | 9,8871493387 | - | - | - | - |
| 11-0005 | pSS | ASSESS Cohort | pSS | F | 61 | 8,843125148 | 0,3656968797 | 24,1815712394 | 7,1034987653 | - | - | - | - |
| 11-0006 | pSS | ASSESS Cohort | pSS | F | 57 | 72,5665211126 | 639,5485048328 | 0,1134652346 | 9,3304367097 | - | - | - | - |
| 11-0007 | pSS | ASSESS Cohort | pSS | F | 59 | 0,0504837085 | 0,3656968797 | 0,1380479608 | 6,5964366931 | 2 | - | 0,83 | 0,75 |
| 11-0008 | pSS | ASSESS Cohort | pSS | F | 47 | 136,2116161741 | 751,9452518673 | 0,1811456563 | 8,7851843236 | - | - | - | - |
| 11-0009 | pSS | ASSESS Cohort | pSS | F | 82 | 66,5486811682 | 0,3656968797 | 181,9777112315 | 9,0898229484 | - | - | - | - |
| 11-0010 | pSS | ASSESS Cohort | pSS | F | 62 | 96,6102199837 | 0,3656968797 | 264,1811438723 | 6,9435040041 | 2 | - | 0,50 | 0,57 |
| 11-0011 | pSS | ASSESS Cohort | pSS | F | 74 | 38,9913927898 | 0,3656968797 | 106,6221643021 | 8,0362299845 | - | - | - | - |
| 11-0012 | pSS | ASSESS Cohort | pSS | F | 63 | 410,9856555704 | 481,6000587485 | 0,8533754266 | 10,4657951283 | 3 | - | 0,31 | 0,44 |
| 11-0013 | pSS | ASSESS Cohort | pSS | F | 48 | 0,0504837085 | 0,3656968797 | 0,1380479608 | 6,6095765533 | 2 | - | 0,44 | 0,39 |
| 11-0014 | pSS | ASSESS Cohort | pSS | F | 45 | 110,5613444778 | 975,4330877836 | 0,1133459033 | 9,7364997009 | - | - | - | - |
| 11-0015 | pSS | ASSESS Cohort | pSS | F | 55 | 399,4156627429 | 858,7655512982 | 0,4651044306 | 10,2219456489 | - | - | - | - |
| 11-0017 | pSS | ASSESS Cohort | pSS | F | 70 | 0,6401417737 | 515,3875829678 | 0,001242059 | 6,7752146097 | - | - | - | - |
| 11-0018 | pSS | ASSESS Cohort | pSS | F | 56 | 1,6732287791 | 0,3656968797 | 4,5754527099 | 6,7449290254 | - | - | - | - |
| 11-0019 | pSS | ASSESS Cohort | pSS | F | 57 | 0,2729513731 | 0,3656968797 | 0,7463869348 | 6,6679787049 | - | - | - | - |
| 11-0020 | pSS | ASSESS Cohort | pSS | F | 56 | 18,6184853656 | 0,3656968797 | 50,9123440758 | 7,7857968798 | - | - | - | - |
| 11-0021 | pSS | ASSESS Cohort | pSS | F | 59 | 0,5515742279 | 0,3656968797 | 1,5082825655 | 6,9668670065 | - | - | - | - |
| 11-0022 | pSS | ASSESS Cohort | pSS | F | 62 | 0,0504837085 | 0,3656968797 | 0,1380479608 | 6,1602427243 | - | - | - | - |
| 11-0023 | pSS | ASSESS Cohort | pSS | F | 74 | 0,0504837085 | 0,3656968797 | 0,1380479608 | 6,6061653974 | - | - | - | - |
| 11-0024 | pSS | ASSESS Cohort | pSS | F | 57 | 11,4291798306 | 0,3656968797 | 31,2531510816 | 7,6967428229 | 2 | - | 0,38 | 0,44 |
| 11-0025 | pSS | ASSESS Cohort | pSS | F | 56 | 16,3666109676 | 0,3656968797 | 44,7545819425 | 7,4060093509 | - | - | - | - |
| 11-0026 | pSS | ASSESS Cohort | pSS | F | 67 | 3,1136841748 | 0,3656968797 | 8,5143854046 | 6,3071057828 | - | - | - | - |
| 11-0027 | pSS | ASSESS Cohort | pSS | F | 77 | 186,9871637977 | 2737,6350904751 | 0,0683024427 | 10,667402245 | - | - | - | - |
| 11-0028 | pSS | ASSESS Cohort | pSS | F | 49 | 185,1781139797 | 92,8501354717 | 1,9943763468 | 9,2054572571 | - | - | - | - |
| 11-0029 | pSS | ASSESS Cohort | pSS | F | 57 | 0,6680724374 | 0,3656968797 | 1,8268475191 | 6,7557580204 | - | - | - | - |

| | | | | | | | | | | | | | |
|---|---|---|---|---|---|---|---|---|---|---|---|---|---|
| 11-0031 | pSS | ASSESS Cohort | pSS | F | 53 | 0,7222207652 | 0,3656968797 | 1,9749164004 | - | - | - | - | - |
| 11-0032 | pSS | ASSESS Cohort | pSS | F | 69 | 53,7559094752 | 0,3656968797 | 146,9958111826 | 8,4417206964 | - | - | - | - |
| 11-0033 | pSS | ASSESS Cohort | pSS | F | 44 | 0,2422412664 | 0,3656968797 | 0,6624099899 | 6,6545680169 | - | - | - | - |
| 11-0036 | pSS | ASSESS Cohort | pSS | F | 50 | 46,1086879854 | 0,3656968797 | 126,0844446527 | 9,5927095128 | 3 | - | 0,62 | 0,35 |
| 11-0037 | pSS | ASSESS Cohort | pSS | F | 53 | 253,8555703585 | 92,5555892821 | 2,7427362553 | 10,1366070572 | - | - | - | - |
| 11-0038 | pSS | ASSESS Cohort | pSS | F | 57 | 160,1586450845 | 28,6210492188 | 5,595834166 | 9,7064431565 | - | - | - | - |
| 11-0039 | pSS | ASSESS Cohort | pSS | F | 53 | 583,8011318722 | 460,4350625857 | 1,2679336986 | 9,913540158 | - | - | - | - |
| 11-0040 | pSS | ASSESS Cohort | pSS | F | 46 | 15,1296662141 | 375,4900785273 | 0,0402931185 | - | - | - | - | - |
| 12-0001 | pSS | ASSESS Cohort | pSS | F | 67 | 136,0349418515 | 22,9542576929 | 5,9263489881 | 9,4356483822 | - | - | - | - |
| 12-0002 | pSS | ASSESS Cohort | pSS | F | 39 | 257,4273922744 | 95,3342501586 | 2,700261363 | - | - | - | - | - |
| 12-0003 | pSS | ASSESS Cohort | pSS | F | 63 | 58,6160286789 | 0,3656968797 | 160,2858321642 | 8,9573449114 | 2 | - | 0,94 | 0,90 |
| 12-0004 | pSS | ASSESS Cohort | pSS | F | 69 | 41,9119563035 | 241,5153573367 | 0,1735374378 | 6,7581959852 | - | - | - | - |
| 12-0005 | pSS | ASSESS Cohort | pSS | F | 59 | 317,4124433611 | 750,2013814272 | 0,4231029844 | 9,6989367234 | - | - | - | - |
| 12-0006 | pSS | ASSESS Cohort | pSS | F | 76 | 71,5262146838 | 0,3656968797 | 195,5888022532 | 7,3004107261 | - | - | - | - |
| 12-0007 | pSS | ASSESS Cohort | pSS | F | 66 | 149,6792753891 | 40,6117934457 | 3,6856110674 | 9,7590570056 | - | - | - | - |
| 12-0008 | pSS | ASSESS Cohort | pSS | F | 75 | 77,3505657177 | 705,3430295649 | 0,1096637558 | 8,8834603929 | - | - | - | - |
| 12-0009 | pSS | ASSESS Cohort | pSS | F | 64 | 0,055995258 | 0,3656968797 | 0,1531193214 | 6,5749345316 | - | - | - | - |
| 12-0010 | pSS | ASSESS Cohort | pSS | F | 58 | 0,4392010054 | 0,3656968797 | 1,2009974103 | 6,4324955342 | - | - | - | - |
| 12-0011 | pSS | ASSESS Cohort | pSS | F | 49 | 152,1107198014 | 146,5512094471 | 1,0379356156 | 9,1182050072 | - | - | - | - |
| 12-0012 | pSS | ASSESS Cohort | pSS | F | 62 | 281,3945830019 | 3709,5844276786 | 0,0758560934 | 10,3007679514 | - | - | - | - |
| 12-0013 | pSS | ASSESS Cohort | pSS | F | 60 | 108,7961954077 | 0,3656968797 | 297,5037563997 | 6,5504092243 | - | - | - | - |
| 12-0014 | pSS | ASSESS Cohort | pSS | F | 57 | 0,055995258 | 0,3656968797 | 0,1531193214 | 6,3956638452 | 2 | - | 0,59 | 0,48 |
| 12-0015 | pSS | ASSESS Cohort | pSS | F | 41 | 66,9757271362 | 2,3996400199 | 27,9107393535 | 8,4107804013 | - | - | - | - |
| 12-0016 | pSS | ASSESS Cohort | pSS | F | 68 | 36,1840526536 | 0,3656968797 | 98,9454782483 | 8,3904219329 | - | - | - | - |
| 12-0017 | pSS | ASSESS Cohort | pSS | F | 37 | 285,6006494322 | 164,4641919298 | 1,7365521703 | 9,9630860573 | - | - | - | - |
| 12-0018 | pSS | ASSESS Cohort | pSS | F | 74 | 1,2391030842 | 0,3656968797 | 3,3883337623 | 7,017719689 | - | - | - | - |
| 12-0019 | pSS | ASSESS Cohort | pSS | F | 53 | 24,1209033014 | 4057,9505045175 | 0,0059441098 | 8,2088194711 | - | - | - | - |
| 12-0020 | pSS | ASSESS Cohort | pSS | F | 58 | 47,3949242833 | 0,3656968797 | 129,6016644308 | 6,5591446367 | - | - | 0,53 | - |
| 12-0021 | pSS | ASSESS Cohort | pSS | F | 59 | 50,8399298202 | 0,3656968797 | 139,0220498054 | 6,4038764042 | - | - | - | - |
| 12-0022 | pSS | ASSESS Cohort | pSS | F | 55 | 108,2976543351 | 386,7614386563 | 0,2800115097 | 9,1212543588 | - | - | - | - |
| 12-0023 | pSS | ASSESS Cohort | pSS | F | 65 | 73,8955334004 | 238,4650826617 | 0,3098798892 | 9,1179083746 | - | - | - | - |
| 12-0024 | pSS | ASSESS Cohort | pSS | F | 67 | 1,9237662464 | 0,3656968797 | 5,2605487039 | 7,5022310604 | - | - | - | - |
| 12-0025 | pSS | ASSESS Cohort | pSS | F | 49 | 0,0619562402 | 0,3656968797 | 0,1694196579 | 5,9205187015 | - | - | - | - |
| 12-0026 | pSS | ASSESS Cohort | pSS | F | 58 | 17,1283585454 | 0,3656968797 | 46,8375846152 | 6,2700824859 | - | - | - | - |
| 12-0027 | pSS | ASSESS Cohort | pSS | F | 69 | 159,8837224296 | 33,3212401639 | 4,7982524553 | - | - | - | - | - |
| 12-0028 | pSS | ASSESS Cohort | pSS | F | 66 | 0,055995258 | 0,3656968797 | 0,1531193214 | 5,7974156446 | - | - | - | - |
| 12-0029 | pSS | ASSESS Cohort | pSS | F | 65 | 87,6202498167 | 70,8317670765 | 1,2370191149 | 8,4869610797 | - | - | - | - |
| 12-0030 | pSS | ASSESS Cohort | pSS | F | 40 | 0,055995258 | 0,3656968797 | 0,1531193214 | 6,5394937124 | 2 | - | 0,89 | 1,02 |
| 13-0001 | pSS | ASSESS Cohort | pSS | F | 41 | 118,4938549511 | 93,3853750647 | 1,2688695084 | 10,5967171445 | - | - | - | - |
| 13-0002 | pSS | ASSESS Cohort | pSS | F | 60 | 5,1559578729 | 0,3656968797 | 14,098993345 | 8,175259488 | - | - | - | - |
| 13-0003 | pSS | ASSESS Cohort | pSS | F | 58 | 0,0633440403 | 0,3656968797 | 0,1732146043 | 7,100174316 | - | - | - | - |
| 13-0004 | pSS | ASSESS Cohort | pSS | F | 29 | 183,2927978907 | 477,6794670802 | 0,3837150443 | 9,2074953589 | 2 | - | 1,56 | 0,78 |
| 13-0005 | pSS | ASSESS Cohort | pSS | F | 66 | 0,802432599 | 0,3656968797 | 2,1942560727 | 7,9036915996 | - | - | - | - |
| 13-0006 | pSS | ASSESS Cohort | pSS | F | 72 | 252,4101243704 | 125,8462392882 | 2,0057025605 | 10,2895581505 | - | - | - | - |
| 13-0007 | pSS | ASSESS Cohort | pSS | F | 68 | 27,8976383785 | 0,3656968797 | 76,2862357566 | 7,5281090572 | - | - | - | - |
| 13-0008 | pSS | ASSESS Cohort | pSS | F | 56 | 0,055995258 | 0,3656968797 | 0,1531193214 | 6,4090231022 | - | - | - | - |
| 13-0009 | pSS | ASSESS Cohort | pSS | F | 41 | 188,9447060023 | 148,3951216825 | 1,27325416 | 10,0153343311 | - | - | - | - |
| 13-0010 | pSS | ASSESS Cohort | pSS | F | 51 | 0,055995258 | 0,3656968797 | 0,1531193214 | 6,287865428 | - | - | - | - |
| 13-0011 | pSS | ASSESS Cohort | pSS | F | 62 | 24,4210819057 | 0,3656968797 | 66,7795741853 | 6,8600026114 | - | - | - | - |
| 13-0012 | pSS | ASSESS Cohort | pSS | F | 51 | 0,5515992472 | 0,3656968797 | 1,5083509811 | 7,5559342886 | - | - | - | - |
| 14-0001 | pSS | ASSESS Cohort | pSS | na | na | 420,9905579926 | 1790,4650310881 | 0,2351291707 | 11,2226260735 | - | - | - | - |
| 14-0002 | pSS | ASSESS Cohort | pSS | F | 75 | 54,4235414307 | 0,3656968797 | 148,8214542019 | 7,107642877 | - | - | - | - |
| 14-0003 | pSS | ASSESS Cohort | pSS | na | na | 222,9936054493 | 0,3656968797 | 609,7771620201 | 6,7930282708 | - | - | - | - |
| 14-0004 | pSS | ASSESS Cohort | pSS | na | na | 0,055995258 | 735,8658608628 | 7,60943821774303E-005 | 6,5062096191 | 3 | - | 2,72 | 2,99 |
| 14-0005 | pSS | ASSESS Cohort | pSS | F | 70 | 0,055995258 | 0,3656968797 | 0,1531193214 | 6,3321809421 | - | - | - | - |
| 14-0006 | pSS | ASSESS Cohort | pSS | M | 50 | 0,2224668582 | 0,3656968797 | 0,6083367691 | 6,8257810114 | - | - | - | - |
| 14-0008 | pSS | ASSESS Cohort | pSS | F | 56 | 116,6839359816 | 3,9206261414 | 29,7615563873 | - | - | - | - | - |
| 15-0002 | pSS | ASSESS Cohort | pSS | F | 66 | 2,0305705291 | 0,3656968797 | 5,5526055646 | 6,7594515211 | - | - | - | - |
| 15-0003 | pSS | ASSESS Cohort | pSS | F | 68 | 2,0562066253 | 0,3656968797 | 5,6227076017 | 7,0551145733 | - | - | - | - |
| 15-0004 | pSS | ASSESS Cohort | pSS | F | 55 | 131,6993037576 | 24,4322888757 | 5,3903792816 | 10,0091451639 | - | - | - | - |
| 15-0005 | pSS | ASSESS Cohort | pSS | F | 73 | 24,1136471825 | 0,3656968797 | 65,938892352 | 8,8713071583 | 2 | - | 0,59 | 0,47 |
| 15-0006 | pSS | ASSESS Cohort | pSS | F | 56 | 0,09855295 | 0,3656968797 | 0,2694935492 | 6,4714869048 | - | - | - | - |
| 15-0010 | pSS | ASSESS Cohort | pSS | M | 63 | 1,4171782258 | 0,3656968797 | 3,8752811537 | 8,2653475079 | - | - | - | - |
| 15-0011 | pSS | ASSESS Cohort | pSS | F | 34 | 46,9000153761 | 0,3656968797 | 128,2483334763 | 8,6541772083 | 2 | - | 2,54 | 2,19 |
| 15-0012 | pSS | ASSESS Cohort | pSS | F | 44 | 269,3401076756 | 0,3656968797 | 736,5119109388 | 6,8057410234 | - | - | - | - |
| 15-0013 | pSS | ASSESS Cohort | pSS | F | 54 | 14,4744740909 | 2533,8679816338 | 0,0057124026 | 6,9892003063 | - | - | - | - |
| 15-0014 | pSS | ASSESS Cohort | pSS | F | 61 | 0,640672129 | 0,3656968797 | 1,7519212347 | 7,236576153 | - | - | - | - |
| 15-0015 | pSS | ASSESS Cohort | pSS | F | 49 | 0,1779386194 | 0,3656968797 | 0,4865740709 | - | - | - | - | - |
| 15-0016 | pSS | ASSESS Cohort | pSS | F | 58 | 76,4329960713 | 84767,8730243785 | 0,0009016741 | 9,9424784362 | 2 | - | 0,68 | 0,50 |
| 15-0017 | pSS | ASSESS Cohort | pSS | F | 55 | 1,2106192328 | 0,3656968797 | 3,3104445241 | 7,6669307016 | - | - | - | - |
| 15-0018 | pSS | ASSESS Cohort | pSS | F | 73 | 2,960100279 | 5090,2283805551 | 0,000581526 | 6,775250903 | 2 | - | 0,44 | 0,50 |
| 15-0019 | pSS | ASSESS Cohort | pSS | F | 35 | 8,4222901122 | 0,3656968797 | 23,0307956678 | 8,6354287757 | - | - | - | - |
| 15-0020 | pSS | ASSESS Cohort | pSS | F | 33 | 20,083514729 | 0,3656968797 | 54,9184744116 | 8,074484636 | 2 | - | 0,41 | 0,38 |

| | | | | | | | | | | | | | |
|---|---|---|---|---|---|---|---|---|---|---|---|---|---|
| 15-0021 | pSS | ASSESS Cohort | pSS | F | 71 | 43,4050576112 | 94,1101730533 | 0,4612153628 | 9,4671244122 | - | - | - | - |
| 15-0022 | pSS | ASSESS Cohort | pSS | F | 82 | 0,5392296183 | 86,463099029 | 0,0062365289 | 6,8961231769 | - | - | - | - |
| 15-0023 | pSS | ASSESS Cohort | pSS | F | 63 | 64,840413814 | 1,2676508947 | 51,1500556553 | 9,7287950938 | - | - | - | - |
| 15-0024 | pSS | ASSESS Cohort | pSS | F | 58 | 290,3514012411 | 144,2036486058 | 2,0134816563 | 10,9671795746 | - | - | - | - |
| 15-0025 | pSS | ASSESS Cohort | pSS | F | 61 | 589,4351672009 | 310,216935191 | 1,9000741105 | 10,5635233164 | - | - | - | - |
| 15-0026 | pSS | ASSESS Cohort | pSS | F | 79 | 64,8449692746 | 0,3656968797 | 177,3189022882 | 8,3244349061 | - | - | - | - |
| 15-0027 | pSS | ASSESS Cohort | pSS | F | 57 | 23,6010472053 | 0,3656968797 | 64,5371850754 | 8,9026500453 | 3 | - | 0,61 | 0,56 |
| 15-0028 | pSS | ASSESS Cohort | pSS | F | 55 | 0,055995258 | 0,3656968797 | 0,1531193214 | 6,7983472182 | - | - | - | - |
| 15-0029 | pSS | ASSESS Cohort | pSS | F | 55 | 0,055995258 | 0,3656968797 | 0,1531193214 | 7,0756863234 | - | - | - | - |
| 15-0030 | pSS | ASSESS Cohort | pSS | F | 66 | 0,9507427284 | 1690,2294765474 | 0,0005624933 | 7,5495796397 | 3 | - | 0,28 | 0,26 |
| 15-0031 | pSS | ASSESS Cohort | pSS | M | 51 | 65,9662136516 | 194,2390002248 | 0,3396136387 | - | - | - | - | - |
| 15-0032 | pSS | ASSESS Cohort | pSS | F | 48 | 4,831699395 | 0,3656968797 | 13,2123068682 | - | - | - | - | - |
| 15-0033 | pSS | ASSESS Cohort | pSS | F | 78 | 1,9667556998 | 0,3656968797 | 5,3781035856 | - | - | - | - | - |
| 23 | SLE | LEAP Cohort | SLE | F | 36 | 2,1641496157 | 24090,415526922 | 8,98344660466568E-005 | 0,815 | 2 | - | 0,52 | 0,46 |
| 24 | SLE | LEAP Cohort | SLE | F | 48 | 602,9752642883 | 1811,3232600988 | 0,3328921334 | 18,956 | 2 | - | 0,32 | 0,46 |
| 27 | SLE | LEAP Cohort | SLE | F | 38 | 15,3653785651 | 5,2586772618 | 2,9219094081 | 0,244 | - | - | - | - |
| 30 | SLE | LEAP Cohort | SLE | F | 53 | 38,5774531054 | 1238,9634722413 | 0,0311368769 | 9,684 | 2 | - | 0,85 | 0,52 |
| 32 | SLE | LEAP Cohort | SLE | F | 42 | 8,0957487974 | 2,792989621 | 2,8985960909 | 2,467 | - | - | - | - |
| 38 | SLE | LEAP Cohort | SLE | F | 65 | 0,1936365603 | 24,4382447634 | 0,0079235052 | 0,267 | - | - | - | - |
| 43 | SLE | LEAP Cohort | SLE | F | 42 | 513,8298794228 | 149501,958069394 | 0,0034369441 | 23,635 | - | - | - | - |
| 46 | SLE | LEAP Cohort | SLE | M | 27 | 21,3000915129 | 1360,3786931997 | 0,0156574722 | 2,128 | 3 | - | 2,43 | 0,92 |
| 51 | SLE | LEAP Cohort | SLE | F | 51 | 4,5262270368 | 0,1387143064 | 32,6298501835 | 0,733 | 2 | - | 0,38 | 0,45 |
| 54 | SLE | LEAP Cohort | SLE | F | 26 | 0,4406325119 | 0,1387143064 | 3,1765469856 | 0,21 | 2 | - | 4,34 | 0,62 |
| 56 | SLE | LEAP Cohort | SLE | F | 60 | 0,8082275461 | 0,1387143064 | 5,8265622845 | 0,562 | 2 | - | 0,37 | 0,41 |
| 61 | SLE | LEAP Cohort | SLE | F | 54 | 34,0964691681 | 22,5752237212 | 1,510349115 | 0,808 | - | - | - | - |
| 62 | SLE | LEAP Cohort | SLE | F | 60 | 1,5417903839 | 0,1387143064 | 11,1148620763 | 1,505 | 2 | - | 0,35 | 0,42 |
| 66 | SLE | LEAP Cohort | SLE | F | 37 | 372,781426882 | 4629,8090436138 | 0,0805176679 | 11,866 | 2 | - | 0,73 | 0,73 |
| 71 | SLE | LEAP Cohort | SLE | F | 53 | 151,805934509 | 184,1901650885 | 0,8241804574 | 18,921 | - | - | - | - |
| 75 | SLE | LEAP Cohort | SLE | F | 23 | 0,3749181124 | 1,4023871003 | 0,267342813 | 0,204 | 2 | - | 0,52 | 0,64 |
| 77 | SLE | LEAP Cohort | SLE | F | 43 | 1,8859541348 | 213,5317115761 | 0,008832197 | 0,304 | - | - | - | - |
| 85 | SLE | LEAP Cohort | SLE | M | 57 | 0,2612905014 | 200000 | 1,30645250698954E-006 | 0,468 | 3 | - | 0,54 | 0,52 |
| 86 | SLE | LEAP Cohort | SLE | F | 29 | 30,9813757936 | 55,7458351866 | 0,5557612634 | 7,231 | 2 | - | 0,30 | 0,46 |
| 87 | SLE | LEAP Cohort | SLE | F | 30 | 2210,5109678407 | 2378,0072074501 | 0,9295644525 | 20,486 | - | - | - | - |
| 88 | SLE | LEAP Cohort | SLE | F | 54 | 0,0071369601 | 5,9190118772 | 0,0012057688 | 0,408 | - | - | - | - |
| 90 | SLE | LEAP Cohort | SLE | F | 63 | 25,9215492301 | 798,0618025307 | 0,0324806289 | 0,523 | 2 | - | 0,69 | 0,86 |
| 91 | SLE | LEAP Cohort | SLE | F | 66 | 0,5878799917 | 0,1387143064 | 4,2380631593 | 0,381 | 2 | - | 0,45 | 0,59 |
| 92 | SLE | LEAP Cohort | SLE | F | 41 | 358,1849295425 | 488,2850479511 | 0,7335570299 | 16,955 | - | - | - | - |
| 94 | SLE | LEAP Cohort | SLE | F | 64 | 15,6635200857 | 1,5595926848 | 10,0433403144 | 2,505 | 3 | - | 0,47 | 0,44 |
| 97 | SLE | LEAP Cohort | SLE | F | 54 | 3,1781921213 | 117,7536584068 | 0,0269901773 | 1,227 | 2 | - | 0,52 | 0,41 |
| 98 | SLE | LEAP Cohort | SLE | F | 29 | 1,4791082114 | 546,2763126587 | 0,0027076192 | 0,638 | - | - | - | - |
| 100 | SLE | LEAP Cohort | SLE | F | 25 | 760,4685526793 | 2860,0186130087 | 0,2658963649 | 12,278 | 2 | - | 0,37 | 0,29 |
| 103 | SLE | LEAP Cohort | SLE | F | 57 | 0,0940478363 | 1767,4707599108 | 5,32104057486822E-005 | 0,45 | 2 | - | 0,48 | 0,49 |
| 105 | SLE | LEAP Cohort | SLE | F | 70 | 2,4275769265 | 0,1387143064 | 17,5005519558 | 0,569 | 2 | - | 0,47 | 0,41 |
| 106 | SLE | LEAP Cohort | SLE | F | 24 | 14,3253013284 | 19,5552564121 | 0,7325550239 | 2,329 | 2 | - | 0,50 | 1,23 |
| 107 | SLE | LEAP Cohort | SLE | F | 48 | 1,6799531433 | 622,0575471375 | 0,0027006394 | 0,932 | - | - | - | - |
| 110 | SLE | LEAP Cohort | SLE | F | 62 | 0,3341268141 | 0,1387143064 | 2,4087408334 | 0,383 | - | - | - | - |
| 112 | SLE | LEAP Cohort | SLE | F | 46 | 67,1191186196 | 365,0720518301 | 0,1838517035 | 11,97 | 2 | - | 0,48 | 0,42 |
| 115 | SLE | LEAP Cohort | SLE | F | 29 | 148,2769069259 | 4543,3417439526 | 0,0326360893 | 13,434 | 3 | - | 0,51 | 0,46 |
| 117 | SLE | LEAP Cohort | SLE | F | 34 | 6,0734793261 | 200000 | 3,03673966303078E-005 | 2,597 | 2 | - | 0,40 | 0,41 |
| 118 | SLE | LEAP Cohort | SLE | F | 47 | 16,3563349033 | 149,6700424741 | 0,1092826235 | 0,571 | 2 | - | 42,49 | 34,52 |
| 121 | SLE | LEAP Cohort | SLE | F | 30 | 0,0299325948 | 6,0606030961 | 0,0049388806 | 4,264 | - | - | - | - |
| 126 | SLE | LEAP Cohort | SLE | F | 55 | 0,2000941909 | 20215,373188104 | 9,89812006240655E-006 | 0,472 | 2 | - | 0,39 | 0,36 |
| F472 | SLE | Rodero *et al.* (2017) | JSLE | F | 15 | 4379,3661250017 | 8725,1053723354 | 0,5019270184 | 25,9 | 100 | - | - | - |
| F885 | SLE | Rodero *et al.* (2017) | JSLE | F | 15 | 21,3990333076 | 6 | 21,3990333076 | 5,364 | 2 | - | - | - |
| F898 | SLE | Rodero *et al.* (2017) | JSLE | F | 12 | 226,5339275769 | 317,9033734276 | 0,7125873662 | 11,3 | - | - | - | - |
| F1017 | SLE | Rodero *et al.* (2017) | JSLE | M | 15 | 9,3759036596 | 19,728236246 | 0,4752530101 | 1,9 | - | - | - | - |
| F1054 | SLE | Rodero *et al.* (2017) | JSLE | F | 11 | 986,2687091943 | 1863,5076467368 | 0,5292539105 | 24 | - | - | - | - |
| F1078 | SLE | Rodero *et al.* (2017) | JSLE | F | 15 | 112,2631579524 | 115,6090619098 | 0,9710584629 | 11,4 | - | - | - | - |
| 889-24 | SLE | Rodero *et al.* (2017) | SLE | F | 28 | 3336,4010924047 | 2223,8313949353 | 1,500294087 | - | 37 | - | - | - |
| LUP001 | SLE | Rodero *et al.* (2017) | SLE | F | 41 | 0,4414537026 | 5,4499265672 | 0,0810017708 | - | - | - | - | - |
| LUP002 | SLE | Rodero *et al.* (2017) | SLE | F | 61 | 186,7555594977 | 437,600458788 | 0,4267718549 | - | - | - | - | - |
| LUP003 | SLE | Rodero *et al.* (2017) | SLE | F | 53 | 60,0024307015 | 118,2902600369 | 0,5072474326 | - | - | - | - | - |
| LUP004 | SLE | Rodero *et al.* (2017) | SLE | F | 55 | 1026,7662672952 | 2794,2872357024 | 0,3674519406 | - | - | - | - | - |
| LUP005 | SLE | Rodero *et al.* (2017) | SLE | F | 59 | 23,1880620298 | 34,202264347 | 0,677968622 | - | - | - | - | - |
| LUP006 | SLE | Rodero *et al.* (2017) | SLE | F | 40 | 850,4457932884 | 2294,5036418875 | 0,3706447781 | - | - | - | - | - |
| LUP007 | SLE | Rodero *et al.* (2017) | SLE | F | 47 | 0,8400037159 | 7,6872904401 | 0,1092717548 | - | - | - | - | - |
| LUP008 | SLE | Rodero *et al.* (2017) | SLE | F | 75 | 9,4386603761 | 23,7407124433 | 0,3975727518 | - | - | - | - | - |
| LUP009 | SLE | Rodero *et al.* (2017) | SLE | F | 20 | 80,7879925549 | 237,2099885388 | 0,3405758461 | - | - | - | - | - |
| LUP010 | SLE | Rodero *et al.* (2017) | SLE | F | 67 | 2,3731159 | 27,7931795757 | 0,0853848295 | - | - | - | - | - |
| LUP011 | SLE | Rodero *et al.* (2017) | SLE | F | 57 | 0,3749730022 | 0,1237496375 | 3,0300937419 | - | - | - | - | - |
| LUP012 | SLE | Rodero *et al.* (2017) | SLE | F | 43 | 336,7224942922 | 1695,8174231759 | 0,1985605819 | - | - | - | - | - |
| LUP013 | SLE | Rodero *et al.* (2017) | SLE | F | 67 | 0,5596949819 | 4,3975960363 | 0,1272729412 | - | - | - | - | - |

| | | | | | | | | | | | | | |
|---|---|---|---|---|---|---|---|---|---|---|---|---|---|
| LUP014 | SLE | Rodero *et al.* (2017) | SLE | F | 51 | 737,6868027731 | 1766,4713842017 | 0,4176047285 | - | - | - | - | - |
| LUP015 | SLE | Rodero *et al.* (2017) | SLE | F | 23 | 5,428428911 | 14,5868935498 | 0,3721442741 | - | - | - | - | - |
| LUP016 | SLE | Rodero *et al.* (2017) | SLE | F | 67 | 2,3927558994 | 5,6826330947 | 0,4210646472 | - | - | - | - | - |
| LUP017 | SLE | Rodero *et al.* (2017) | SLE | F | 34 | 4,3093162231 | 7,1980884255 | 0,5986750882 | - | - | - | - | - |
| LUP018 | SLE | Rodero *et al.* (2017) | SLE | F | 30 | 153,370250892 | 428,7585803954 | 0,3577077122 | - | - | - | - | - |
| LUP019 | SLE | Rodero *et al.* (2017) | SLE | F | 76 | 0,3749730022 | 2,3772711179 | 0,157732536 | - | - | - | - | - |
| LUP020 | SLE | Rodero *et al.* (2017) | SLE | F | 54 | 9,1550403341 | 9,1335519122 | 1,0023526906 | - | - | - | - | - |
| LUP021 | SLE | Rodero *et al.* (2017) | SLE | F | 70 | 18,8636182994 | 18,9313543944 | 0,9964220154 | - | - | - | - | - |
| LUP022 | SLE | Rodero *et al.* (2017) | SLE | F | 28 | 165,3054807233 | 286,6292390073 | 0,5767223236 | 28,6 | - | - | - | - |
| LUP023 | SLE | Rodero *et al.* (2017) | SLE | F | 50 | 0,1250619226 | 1,1750478776 | 0,106431342 | 0,47 | - | - | - | - |
| LUP024 | SLE | Rodero *et al.* (2017) | SLE | F | 47 | 0,1793197514 | 1,8109665778 | 0,0990188077 | 0,74 | - | - | - | - |
| LUP025 | SLE | Rodero *et al.* (2017) | SLE | F | 63 | 1823,434513892 | 2731,358926985 | 0,6675924192 | 27,32 | - | - | - | - |
| LUP026 | SLE | Rodero *et al.* (2017) | SLE | F | 67 | 50,959399303 | 125,1545910692 | 0,4071716336 | 16,05 | - | - | - | - |
| LUP027 | SLE | Rodero *et al.* (2017) | SLE | F | 42 | 1204,6447714858 | 2193,1025281628 | 0,5492879407 | 22,84 | - | - | - | - |
| LUP028 | SLE | Rodero *et al.* (2017) | SLE | F | 43 | 35,5339592637 | 80,5384140878 | 0,4412051028 | 12,23 | - | - | - | - |
| LUP029 | SLE | Rodero *et al.* (2017) | SLE | F | 40 | 0,3749730022 | 0,1237496375 | 3,0300937419 | 25,11 | - | - | - | - |
| LUP029b | SLE | Rodero *et al.* (2017) | SLE | F | 40 | 93,5169046689 | 174,8074037181 | 0,5349710749 | - | - | - | - | - |
| LUP030 | SLE | Rodero *et al.* (2017) | SLE | F | 67 | 3,0726860991 | 6,7996935953 | 0,4518859646 | 1,92 | - | - | - | - |
| LUP032 | SLE | Rodero *et al.* (2017) | SLE | M | 60 | 209,5064554759 | 414,9833769109 | 0,5048550548 | 16,7 | - | - | - | - |
| LUP034 | SLE | Rodero *et al.* (2017) | SLE | F | 38 | 580,4515830865 | 873,4155818826 | 0,664576629 | 24,05 | - | - | - | - |
| LUP035 | SLE | Rodero *et al.* (2017) | SLE | F | 41 | 0,127835833 | 1,2712256453 | 0,1005610872 | 1,11 | - | - | - | - |
| LUP036 | SLE | Rodero *et al.* (2017) | SLE | M | 32 | 1,5892475968 | 2477,9522628455 | 0,0006413552 | 0,31 | - | - | - | - |
| LUP037 | SLE | Rodero *et al.* (2017) | SLE | F | 41 | 0,1076691437 | 0,1237496375 | 0,8700562348 | 0,75 | - | - | - | - |
| LUP039 | SLE | Rodero *et al.* (2017) | SLE | F | 48 | 0,1799252763 | 1,0977371714 | 0,163905606 | 0,18 | - | - | - | - |
| LUP040 | SLE | Rodero *et al.* (2017) | SLE | F | 50 | 56,7644338621 | 86,0047237398 | 0,6600153037 | 12,39 | - | - | - | - |
| LUP041 | SLE | Rodero *et al.* (2017) | SLE | F | nd | 0,1076691437 | 6,1956420688 | 0,0173782059 | - | - | - | - | - |
| LUP042 | SLE | Rodero *et al.* (2017) | SLE | F | 45 | 9,2755305466 | 3,9 | 9,2755305466 | 6,64 | - | - | - | - |
| LUP043 | SLE | Rodero *et al.* (2017) | SLE | F | 66 | 15,7338684902 | 34,9451138059 | 0,4502451638 | 6,81 | - | - | - | - |
| LUP044 | SLE | Rodero *et al.* (2017) | SLE | F | 24 | 0,7690019722 | 7,1213696568 | 0,1079851221 | 0,21 | - | - | - | - |
| LUP045 | SLE | Rodero *et al.* (2017) | SLE | F | 54 | 0,4287595569 | 3,5749206981 | 0,1199354036 | 0,17 | - | - | - | - |
| LUP046 | SLE | Rodero *et al.* (2017) | SLE | F | 52 | 83,7292104014 | 3,9 | 83,7292104014 | 0,27 | - | - | - | - |
| LUP047 | SLE | Rodero *et al.* (2017) | SLE | F | 30 | 7,6899230066 | 17,6943303941 | 0,434598136 | - | - | - | - | - |
| LUP048 | SLE | Rodero *et al.* (2017) | SLE | F | 57 | 118,7118124133 | 262,1417878016 | 0,4528534478 | 17,9 | - | - | - | - |
| LUP049 | SLE | Rodero *et al.* (2017) | SLE | F | 22 | 694,1942306653 | 2248,4691619653 | 0,3087408279 | 19,23 | - | - | - | - |
| LUP050 | SLE | Rodero *et al.* (2017) | SLE | F | 40 | 0,1280984679 | 5,9761714909 | 0,0214348715 | 0,26 | - | - | - | - |
| LUP051 | SLE | Rodero *et al.* (2017) | SLE | M | 44 | 11,2936210505 | 20,8279298924 | 0,5422344472 | 5,01 | - | - | - | - |
| SLE 022 | SLE | Menon *et al.* (date) | SLE | F | 66 | 8018,7581373323 | 2299,5605967533 | 3,4870827708 | - | - | 55,4 | - | - |
| SLE 039 | SLE | Menon *et al.* (date) | SLE | F | 63 | 47,1896918367 | 33,6839934813 | 1,4009530035 | - | - | 69,1 | - | - |
| SLE 046 | SLE | Menon *et al.* (date) | SLE | F | 56 | 0,0302100365 | 22666,4900616173 | 1,33280611356899E-006 | - | - | 154,5 | - | - |
| SLE 052 | SLE | Menon *et al.* (date) | SLE | F | 58 | 40765,9262517692 | 57151,8992659433 | 0,7132908403 | - | - | Negative | - | - |
| SLE 060 | SLE | Menon *et al.* (date) | SLE | F | 74 | 0,0541859678 | 17301,2445481428 | 3,13191155983809E-006 | - | - | 143,0 | - | - |
| SLE 073 | SLE | Menon *et al.* (date) | SLE | F | 72 | 0,1018946191 | 14,0421511222 | 0,0072563397 | - | - | Negative | - | - |
| SLE 074 | SLE | Menon *et al.* (date) | SLE | F | 68 | 22,6786070038 | 15,8024776717 | 1,4351298243 | - | - | Negative | - | - |
| SLE 087 | SLE | Menon *et al.* (date) | SLE | F | 55 | 58,1177174543 | 73,1477308881 | 0,7945252265 | - | - | Negative | - | - |
| SLE 090 | SLE | Menon *et al.* (date) | SLE | F | 55 | 0,8066598225 | 1,0645136885 | 0,7577730857 | - | - | Negative | - | - |
| SLE 101 | SLE | Menon *et al.* (date) | SLE | F | 66 | 191,2891599184 | 272,1322984715 | 0,7029270726 | - | - | Negative | - | - |
| SLE 103 | SLE | Menon *et al.* (date) | SLE | F | 79 | 761,8961546149 | 2480,1679260455 | 0,3071953905 | - | - | 85,8 | - | - |
| SLE 104 | SLE | Menon *et al.* (date) | SLE | F | 87 | 0,0557516799 | 60,3225283922 | 0,0009242265 | - | - | 148,6 | - | - |
| SLE 106 | SLE | Menon *et al.* (date) | SLE | F | 63 | 533,9738972051 | 5065,8684785307 | 0,1054061904 | - | - | Negative | - | - |
| SLE 107 | SLE | Menon *et al.* (date) | SLE | F | 70 | 22,9443666977 | 32,7818856221 | 0,6999099125 | - | - | Negative | - | - |
| SLE 117 | SLE | Menon *et al.* (date) | SLE | F | 74 | 169,4888382626 | 409,1548020567 | 0,4142413517 | - | - | 2,4 | - | - |
| SLE 119 | SLE | Menon *et al.* (date) | SLE | F | 57 | 33,6124801714 | 2508,841298714 | 0,0133976112 | - | - | 82,9 | - | - |
| SLE 123 | SLE | Menon *et al.* (date) | SLE | F | 60 | 0,1018946191 | 130954,726376978 | 7,78090428400286E-007 | - | - | Negative | - | - |
| SLE 129 | SLE | Menon *et al.* (date) | SLE | F | 79 | 105,1997272311 | 89,6098847549 | 1,1739745846 | - | - | 15,2 | - | - |
| SLE 132 | SLE | Menon *et al.* (date) | SLE | F | 82 | 1,585732227 | 1,0645136885 | 1,4896306587 | - | - | Negative | - | - |
| SLE 141 | SLE | Menon *et al.* (date) | SLE | F | 63 | 12,0392238403 | 0,7164402454 | 16,8042260569 | - | - | 32,1 | - | - |
| SLE 158 | SLE | Menon *et al.* (date) | SLE | F | 60 | 1358,1980025573 | 0,7164402454 | 1895,75894325 | - | - | 106,8 | - | - |
| SLE 159 | SLE | Menon *et al.* (date) | SLE | F | 74 | 0,920876054 | 17,7912961934 | 0,0517599192 | - | - | 3,6 | - | - |
| SLE 163 | SLE | Menon *et al.* (date) | SLE | F | 55 | 4333,3993807468 | 5065,9996542228 | 0,8553888031 | - | - | 2,4 | - | - |
| SLE 173 | SLE | Menon *et al.* (date) | SLE | F | 61 | 222,6361729602 | 8095,3382221458 | 0,0275017753 | - | - | Negative | - | - |
| SLE 181 | SLE | Menon *et al.* (date) | SLE | F | 57 | 0,9288492997 | 194,7773199263 | 0,0047687754 | - | - | 3,7 | - | - |
| SLE 187 | SLE | Menon *et al.* (date) | SLE | F | 48 | 221,3109725062 | 26,5143004428 | 8,3468531626 | - | - | Negative | - | - |
| SLE 194 | SLE | Menon *et al.* (date) | SLE | F | 46 | 51,844656135 | 18,1186338264 | 2,8613998512 | - | - | 4,8 | - | - |
| SLE 219 | SLE | Menon *et al.* (date) | SLE | F | 49 | 19,9890909162 | 0,7164402454 | 27,9005695785 | - | - | 3,9 | - | - |
| SLE 220 | SLE | Menon *et al.* (date) | SLE | F | 53 | 1,3899634955 | 1,0645136885 | 1,3057262772 | - | - | Negative | - | - |
| SLE 229 | SLE | Menon *et al.* (date) | SLE | F | 52 | 112,1902349159 | 33956,7888199501 | 0,0033039118 | - | - | 40,5 | - | - |
| SLE 237 | SLE | Menon *et al.* (date) | SLE | M | 40 | 820,5883463468 | 52910,8139804531 | 0,0155088967 | - | - | 86,9 | - | - |
| SLE 247 | SLE | Menon *et al.* (date) | SLE | F | 55 | 128,6196849144 | 45,586848778 | 2,8214208343 | - | - | Negative | - | - |
| SLE 250 | SLE | Menon *et al.* (date) | SLE | F | 53 | 83,8767387885 | 103,909809872 | 0,8072071241 | - | - | Negative | - | - |
| SLE 259 | SLE | Menon *et al.* (date) | SLE | F | 61 | 165,25802702 | 156,5649771669 | 1,0555235916 | - | - | 1,9 | - | - |

| | | | | | | | | | | | | | |
|---|---|---|---|---|---|---|---|---|---|---|---|---|---|
| SLE 265 | SLE | Menon *et al.* (date) | SLE | F | 83 | 90,4422165246 | 17,383997785 | 5,2026132103 | - | - | 8,8 | - | - |
| SLE 271 | SLE | Menon *et al.* (date) | SLE | F | 71 | 0,0302100365 | 0,7164402454 | 0,0421668614 | - | - | 117,6 | - | - |
| SLE 279 | SLE | Menon *et al.* (date) | SLE | F | 42 | 89,4339919537 | 62,4735508632 | 1,4315496833 | - | - | Negative | - | - |
| SLE 280 | SLE | Menon *et al.* (date) | SLE | F | 63 | 513,8418815914 | 904,6311872714 | 0,5680125656 | - | - | Negative | - | - |
| SLE 287 | SLE | Menon *et al.* (date) | SLE | F | 79 | 0,0076771615 | 18622,2328679674 | 4,12257837458629E-007 | - | - | 31,2 | - | - |
| SLE 288 | SLE | Menon *et al.* (date) | SLE | F | 62 | 1,6601652064 | 7,7736137711 | 0,2135641486 | - | - | 6,7 | - | - |
| SLE 290 | SLE | Menon *et al.* (date) | SLE | F | 42 | 468,1963099834 | 1002,0630363741 | 0,4672323926 | - | - | Negative | - | - |
| SLE 296 | SLE | Menon *et al.* (date) | SLE | F | 52 | 17,9356360752 | 0,7164402454 | 25,034378219 | - | - | 107,2 | - | - |
| SLE 297 | SLE | Menon *et al.* (date) | SLE | F | 52 | 998,4197031333 | 119,987766733 | 8,3210124692 | - | - | 44,5 | - | - |
| SLE 299 | SLE | Menon *et al.* (date) | SLE | M | 44 | 0,0302100365 | 6280,4744234737 | 4,81015198698405E-006 | - | - | 141,9 | - | - |
| SLE 301 | SLE | Menon *et al.* (date) | SLE | F | 61 | 10,5825325131 | 3,4967507204 | 3,0263903147 | - | - | Negative | - | - |
| SLE 309 | SLE | Menon *et al.* (date) | SLE | F | 39 | 90,1541265539 | 969,7210513426 | 0,0929691342 | - | - | Negative | - | - |
| SLE 311 | SLE | Menon *et al.* (date) | SLE | M | 66 | 2753,1183999504 | 4760,1495900005 | 0,5783680424 | - | - | 4,9 | - | - |
| SLE 312 | SLE | Menon *et al.* (date) | SLE | F | 54 | 1,641767001 | 22862,0742101894 | 7,18118131321243E-005 | - | - | 5,3 | - | - |
| SLE 319 | SLE | Menon *et al.* (date) | SLE | M | 56 | 0,6855057722 | 4401,0761586138 | 0,0001557587 | - | - | 3,1 | - | - |
| SLE 342 | SLE | Menon *et al.* (date) | SLE | F | 77 | 170,3340860323 | 135,4012529077 | 1,2579949031 | - | - | 9,3 | - | - |
| SLE 344 | SLE | Menon *et al.* (date) | SLE | F | 57 | 1668,9816273474 | 1860,9361733456 | 0,8968505483 | - | - | 2,4 | - | - |
| SLE 355 | SLE | Menon *et al.* (date) | SLE | F | 36 | 6,8546942612 | 11,5779229641 | 0,5920487019 | - | - | Negative | - | - |
| SLE 356 | SLE | Menon *et al.* (date) | SLE | F | 55 | 287,8035329621 | 34535,0745910915 | 0,0083336589 | - | - | 3,0 | - | - |
| SLE 363 | SLE | Menon *et al.* (date) | SLE | F | 41 | 1223,4553354119 | 2518,0730438343 | 0,4858696766 | - | - | 4,8 | - | - |
| SLE 364 | SLE | Menon *et al.* (date) | SLE | F | 35 | 10,0308340123 | 1972,9021235153 | 0,0050843039 | - | - | Negative | - | - |
| SLE 365 | SLE | Menon *et al.* (date) | SLE | M | 82 | 305,5581285579 | 238,8406846628 | 1,2793386897 | - | - | 2,1 | - | - |
| SLE 372 | SLE | Menon *et al.* (date) | SLE | F | 76 | 0,9056687862 | 1,6523004596 | 0,5481259664 | - | - | Negative | - | - |
| SLE 375 | SLE | Menon *et al.* (date) | SLE | F | 48 | 0,7181839643 | 1,9864276488 | 0,3615454934 | - | - | 13,1 | - | - |
| SLE 383 | SLE | Menon *et al.* (date) | SLE | F | 48 | 3456,3358531607 | 3739,9824448263 | 0,9241583093 | - | - | Negative | - | - |
| SLE 385 | SLE | Menon *et al.* (date) | SLE | F | 36 | 24,9940034193 | 683,7610501061 | 0,0365537104 | - | - | Negative | - | - |
| SLE 393 | SLE | Menon *et al.* (date) | SLE | F | 53 | 126,016115263 | 314,0289058478 | 0,4012882665 | - | - | Negative | - | - |
| SLE 398 | SLE | Menon *et al.* (date) | SLE | F | 57 | 61,4297856114 | 22,6530430428 | 2,7117674873 | - | - | 35,5 | - | - |
| SLE 399 | SLE | Menon *et al.* (date) | SLE | F | 37 | 19,0218332165 | 1,0645136885 | 17,8690358061 | - | - | Negative | - | - |
| SLE 400 | SLE | Menon *et al.* (date) | SLE | F | 64 | 3322,8648263053 | 146,602294019 | 22,6658446823 | - | - | 93,0 | - | - |
| SLE 402 | SLE | Menon *et al.* (date) | SLE | F | 59 | 280,4435773145 | 282,282116452 | 0,9934868735 | - | - | Negative | - | - |
| SLE 413 | SLE | Menon *et al.* (date) | SLE | M | 52 | 2,5401172979 | 3810,7450302408 | 0,0006665671 | - | - | 3,3 | - | - |
| SLE 422 | SLE | Menon *et al.* (date) | SLE | F | 32 | 1108,5326474742 | 8749,0701941856 | 0,1267029093 | - | - | 11,5 | - | - |
| SLE 423 | SLE | Menon *et al.* (date) | SLE | F | 40 | 14428,546275388 | 23717,2010672985 | 0,6083578848 | - | - | 19,8 | - | - |
| SLE 426 | SLE | Menon *et al.* (date) | SLE | F | 41 | 4,9819059568 | 8,7363643571 | 0,5702493341 | - | - | 12,8 | - | - |
| SLE 427 | SLE | Menon *et al.* (date) | SLE | F | 59 | 14,598955173 | 10,3118225843 | 1,4157492581 | - | - | Negative | - | - |
| SLE 428 | SLE | Menon *et al.* (date) | SLE | F | 52 | 15,6236488857 | 1,0645136885 | 14,6767947223 | - | - | Negative | - | - |
| SLE 431 | SLE | Menon *et al.* (date) | SLE | M | 74 | 1,4392628134 | 0,7164402454 | 2,0089083809 | - | - | 0,8 | - | - |
| SLE 433 | SLE | Menon *et al.* (date) | SLE | F | 50 | 329,5640265235 | 591,6112630115 | 0,5570617855 | - | - | Negative | - | - |
| SLE 438 | SLE | Menon *et al.* (date) | SLE | F | 33 | 338,8349706056 | 1,3781146456 | 245,8684926387 | - | - | 17,3 | - | - |
| SLE 442 | SLE | Menon *et al.* (date) | SLE | F | 35 | 2884,511153828 | 24214,6311443306 | 0,1191226551 | - | - | Negative | - | - |
| SLE 446 | SLE | Menon *et al.* (date) | SLE | M | 32 | 327,3428454992 | 341,094042833 | 0,9596850264 | - | - | Negative | - | - |
| SLE 460 | SLE | Menon *et al.* (date) | SLE | F | 38 | 0,0302100365 | 0,7164402454 | 0,0421668614 | - | - | 99,7 | - | - |
| SLE 462 | SLE | Menon *et al.* (date) | SLE | F | 54 | 1634,5753597097 | 455,4620395933 | 3,588828964 | - | - | Negative | - | - |
| SLE 473 | SLE | Menon *et al.* (date) | SLE | F | 65 | 32,4456442644 | 247,2867946018 | 0,1312065382 | - | - | 3,4 | - | - |
| SLE 485 | SLE | Menon *et al.* (date) | SLE | F | 63 | 184,6349358456 | 21,8091946282 | 8,4659217818 | - | - | 50,9 | - | - |
| SLE 486 | SLE | Menon *et al.* (date) | SLE | F | 39 | 404,0873885436 | 230,5405627559 | 1,752782173 | - | - | Negative | - | - |
| SLE 486-2 | SLE | Menon *et al.* (date) | SLE | F | 39 | 671,858325226 | 749,6822385286 | 0,8961908002 | - | - | Negative | - | - |
| SLE 489 | SLE | Menon *et al.* (date) | SLE | F | 36 | 4425,00113404 | 9494,524703484 | 0,4660582043 | - | - | Negative | - | - |
| SLE 491 | SLE | Menon *et al.* (date) | SLE | F | 34 | 65,9264191837 | 2410,9564835179 | 0,0273445081 | - | - | 22,6 | - | - |
| SLE 492 | SLE | Menon *et al.* (date) | SLE | F | 49 | 143,6817381667 | 31,9125837691 | 4,5023536548 | - | - | 0,5 | - | - |
| SLE 497 | SLE | Menon *et al.* (date) | SLE | F | 72 | 0,0302100365 | 0,7164402454 | 0,0421668614 | - | - | 196,4 | - | - |
| SLE 499 | SLE | Menon *et al.* (date) | SLE | F | 36 | 23,6710000617 | 427,7345303754 | 0,0553404001 | - | - | Negative | - | - |
| SLE 504 | SLE | Menon *et al.* (date) | SLE | F | 41 | 260,3181006948 | 404,2203354089 | 0,6440005064 | - | - | Negative | - | - |
| SLE 507 | SLE | Menon *et al.* (date) | SLE | F | 46 | 666,9558295339 | 1163,0576436446 | 0,5734503644 | - | - | 10,7 | - | - |
| SLE 508 | SLE | Menon *et al.* (date) | SLE | F | 50 | 277,9570755061 | 549,098632842 | 0,5062060965 | - | - | Negative | - | - |
| SLE 510 | SLE | Menon *et al.* (date) | SLE | F | 39 | 446,5518098769 | 1271,3677634486 | 0,3512373231 | - | - | Negative | - | - |
| SLE 512 | SLE | Menon *et al.* (date) | SLE | F | 48 | 1012,8974471876 | 515,9295943858 | 1,9632474241 | - | - | 32,7 | - | - |
| SLE 513 | SLE | Menon *et al.* (date) | SLE | M | 31 | 1432,8172579545 | 4685,168833919 | 0,3058197706 | - | - | Negative | - | - |
| SLE 514 | SLE | Menon *et al.* (date) | SLE | M | 36 | 838,4973477199 | 8132,8255986831 | 0,1031003724 | - | - | 12,6 | - | - |
| SLE 526 | SLE | Menon *et al.* (date) | SLE | F | 32 | 46,0959242067 | 0,7164402454 | 64,3402216744 | - | - | 118,3 | - | - |
| SLE 531 | SLE | Menon *et al.* (date) | SLE | F | 37 | 764,1122995129 | 2087,5847746226 | 0,3660269556 | - | - | 5,3 | - | - |
| SLE 536 | SLE | Menon *et al.* (date) | SLE | F | 34 | 1226,7427064559 | 3,9809448164 | 308,1536577423 | - | - | 70,2 | - | - |
| SLE 537 | SLE | Menon *et al.* (date) | SLE | F | 42 | 940,9313321034 | 1324,8754281135 | 0,7102036253 | - | - | Negative | - | - |
| SLE 537-2 | SLE | Menon *et al.* (date) | SLE | F | 42 | 578,1867708157 | 565,9283654405 | 1,0216607015 | - | - | Negative | - | - |
| SLE 543 | SLE | Menon *et al.* (date) | SLE | M | 35 | 207,7102108572 | 1876,0422543433 | 0,1107172348 | - | - | Negative | - | - |
| SLE 547 | SLE | Menon *et al.* (date) | SLE | F | 66 | 1754,987802599 | 3108,6724691317 | 0,564545741 | - | - | Negative | - | - |
| SLE 547-2 | SLE | Menon *et al.* (date) | SLE | F | 66 | 3043,1245531856 | 4571,1865004956 | 0,6657187478 | - | - | Negative | - | - |
| SLE 552 | SLE | Menon *et al.* (date) | SLE | F | 40 | 2,5003372676 | 1,0645136885 | 2,3488070604 | - | - | Negative | - | - |
| SLE 559 | SLE | Menon *et al.* (date) | SLE | F | 46 | 161,3527276019 | 145,9581978698 | 1,1054721828 | - | - | 92,9 | - | - |

| | | | | | | | | | | | | | |
|---|---|---|---|---|---|---|---|---|---|---|---|---|---|
| SLE 564 | SLE | Menon *et al.* (date) | SLE | F | 42 | 6599,4930291202 | 7715,4572763932 | 0,8553599343 | - | - | 5,5 | - | - |
| SLE 566 | SLE | Menon *et al.* (date) | SLE | F | 45 | 404,1409438649 | 36322,3128116462 | 0,0111265201 | - | - | Negative | - | - |
| SLE 583 | SLE | Menon *et al.* (date) | SLE | F | 27 | 614,9032161999 | 832,3096258454 | 0,738791427 | - | - | Negative | - | - |
| SLE 590 | SLE | Menon *et al.* (date) | SLE | F | 54 | 499,4188620808 | 35891,3297465545 | 0,0139147495 | - | - | Negative | - | - |
| SLE 590-2 | SLE | Menon *et al.* (date) | SLE | F | 54 | 1304,7024392359 | - | - | - | - | Negative | - | - |
| SLE 591 | SLE | Menon *et al.* (date) | SLE | F | 30 | 700,9387690873 | 1955,2478721184 | 0,3584910021 | - | - | Negative | - | - |
| SLE 621 | SLE | Menon *et al.* (date) | SLE | F | 35 | 152,1751976093 | 91,0459104591 | 1,671411674 | - | - | 17,6 | - | - |
| 1BB01 | cHCV | C10-08 | cHCV | M | 66 | 134,4622130174 | 44,65411275 | 3,0111943724 | - | - | - | - | - |
| 1BF12 | cHCV | C10-08 | cHCV | F | 67 | 24,5611984826 | 241,6622325 | 0,1016344103 | - | - | - | - | - |
| 1CJ11 | cHCV | C10-08 | cHCV | M | 48 | 19,1511719097 | 415,58153 | 0,0460828274 | - | - | - | - | - |
| 1DJ10 | cHCV | C10-08 | cHCV | M | 67 | 27,1221962889 | 1467,933532 | 0,0184764471 | - | - | - | - | - |
| 1DJ13 | cHCV | C10-08 | cHCV | M | 45 | 5,9605186985 | 1 | 5,9605186985 | - | - | - | - | - |
| 1FI03 | cHCV | C10-08 | cHCV | F | 53 | 1,6021001265 | 1 | 1,6021001265 | - | - | - | - | - |
| 1JG08 | cHCV | C10-08 | cHCV | M | 55 | 2,9016032213 | 1 | 2,9016032213 | - | - | - | - | - |
| 1LA20 | cHCV | C10-08 | cHCV | M | 59 | 2,4971922861 | 1 | 2,4971922861 | - | - | - | - | - |
| 1LE06 | cHCV | C10-08 | cHCV | M | 55 | 36,762603035 | 30,61968 | 1,2006200925 | - | - | - | - | - |
| 1MJ02 | cHCV | C10-08 | cHCV | F | 57 | 6,3718340257 | 1 | 6,3718340257 | - | - | - | - | - |
| 1ML07 | cHCV | C10-08 | cHCV | M | 48 | 33,849811512 | 1 | 33,8498115121 | - | - | - | - | - |
| 1MM16 | cHCV | C10-08 | cHCV | F | 55 | 11,0642009452 | 51,69164325 | 0,2140423529 | - | - | - | - | - |
| 1MP14 | cHCV | C10-08 | cHCV | M | 54 | 7,2798738677 | 672,83086 | 0,0108197681 | - | - | - | - | - |
| 1PA05 | cHCV | C10-08 | cHCV | M | 52 | 2,8181507099 | 13,2438605 | 0,2127892173 | - | - | - | - | - |
| 1PJ17 | cHCV | C10-08 | cHCV | M | 75 | 260,4655286802 | 475,27428 | 0,5480320304 | - | - | - | - | - |
| 1RL09 | cHCV | C10-08 | cHCV | M | 46 | 4,5590625919 | 639,6312419 | 0,0071276421 | - | - | - | - | - |
| 1SS18 | cHCV | C10-08 | cHCV | M | 68 | 4,1861593364 | 1 | 4,1861593364 | - | - | - | - | - |
| 1WC04 | cHCV | C10-08 | cHCV | F | 57 | 23,088640132 | 1 | 23,088640132 | - | - | - | - | - |
| 2BC23 | cHCV | C10-08 | cHCV | F | 52 | 4,180404001 | 1 | 4,180404001 | - | - | - | - | - |
| 2BF15 | cHCV | C10-08 | cHCV | M | 65 | 4,4628856029 | 1 | 4,4628856029 | - | - | - | - | - |
| 2BL13 | cHCV | C10-08 | cHCV | F | 44 | 13,0904463382 | 1 | 13,0904463382 | - | - | - | - | - |
| 2BO11 | cHCV | C10-08 | cHCV | M | 56 | 4,2288234563 | 1 | 4,2288234563 | - | - | - | - | - |
| 2BY08 | cHCV | C10-08 | cHCV | F | 64 | 17,5796795669 | 1 | 17,5796795669 | - | - | - | - | - |
| 2CF18 | cHCV | C10-08 | cHCV | M | 77 | 22,199400926 | 1 | 22,199400926 | - | - | - | - | - |
| 2DF19 | cHCV | C10-08 | cHCV | F | 70 | 2,2048454601 | 40,07025425 | 0,0550244939 | - | - | - | - | - |
| 2DM14 | cHCV | C10-08 | cHCV | F | 60 | 2,943497896 | 1 | 2,943497896 | - | - | - | - | - |
| 2FM24 | cHCV | C10-08 | cHCV | M | 36 | 4,5770426016 | 1 | 4,5770426016 | - | - | - | - | - |
| 2IR1 | cHCV | C10-08 | cHCV | F | 38 | 2,5537507943 | 1 | 2,5537507943 | - | - | - | - | - |
| 2JP07 | cHCV | C10-08 | cHCV | M | 59 | 1,8317725842 | 1 | 1,8317725842 | - | - | - | - | - |
| 2KA05 | cHCV | C10-08 | cHCV | F | 58 | 55,3901134564 | 1 | 55,3901134564 | - | - | - | - | - |
| 2LA21 | cHCV | C10-08 | cHCV | M | 60 | 4,3009882364 | 1 | 4,3009882364 | - | - | - | - | - |
| 2LG09 | cHCV | C10-08 | cHCV | F | 72 | 11,9718358851 | 1 | 11,9718358851 | - | - | - | - | - |
| 2LM16 | cHCV | C10-08 | cHCV | M | 66 | 11,7628390521 | 1 | 11,7628390521 | - | - | - | - | - |
| 2MC10 | cHCV | C10-08 | cHCV | F | 48 | 9,3161260217 | 72,53955475 | 0,1284282217 | - | - | - | - | - |
| 2MC22 | cHCV | C10-08 | cHCV | M | 54 | 7,4554653259 | 2297,5823 | 0,0032449176 | - | - | - | - | - |
| 2MM06 | cHCV | C10-08 | cHCV | F | 73 | 6,4638066156 | 1 | 6,4638066156 | - | - | - | - | - |
| 2NA3 | cHCV | C10-08 | cHCV | M | 42 | 4,9924365579 | 1361,855625 | 0,0036659074 | - | - | - | - | - |
| 2NE12 | cHCV | C10-08 | cHCV | F | 70 | 7,4294258872 | 1 | 7,4294258872 | - | - | - | - | - |
| 2SC17 | cHCV | C10-08 | cHCV | F | 57 | 8,5507784633 | 1 | 8,5507784633 | - | - | - | - | - |
| 2SJ02 | cHCV | C10-08 | cHCV | M | 46 | 2,6474235313 | 1 | 2,6474235313 | - | - | - | - | - |
| 2SS04 | cHCV | C10-08 | cHCV | F | 48 | 116,7232186424 | 40,96211815 | 2,8495405979 | - | - | - | - | - |
| 2TM25 | cHCV | C10-08 | cHCV | F | 63 | 1,5412211307 | 1 | 1,5412211307 | - | - | - | - | - |
| 2ZB20 | cHCV | C10-08 | cHCV | F | 59 | 2,8358443142 | 152,710155 | 0,0185701096 | - | - | - | - | - |
| 3AM01 | cHCV | C10-08 | cHCV | F | 53 | 1,8917343714 | 1 | 1,8917343714 | - | - | - | - | - |
| 3AS02 | cHCV | C10-08 | cHCV | M | 55 | 2,4792178193 | 11,421576075 | 0,2170644229 | - | - | - | - | - |
| 3BJ11-3BS11 | cHCV | C10-08 | cHCV | M | 60 | 16,4283881286 | 1338,17255 | 0,0122767338 | - | - | - | - | - |
| 3CH12 | cHCV | C10-08 | cHCV | M | 47 | 0,2057658292 | 482,0412266 | 0,0004268636 | - | - | - | - | - |
| 3CM05 | cHCV | C10-08 | cHCV | F | 75 | 5,7355141391 | 69,260945 | 0,0828102207 | - | - | - | - | - |
| 3HS06 | cHCV | C10-08 | cHCV | F | 66 | 11,8176073129 | 8,8547485 | 1,3346067721 | - | - | - | - | - |
| 3MG04 | cHCV | C10-08 | cHCV | F | 51 | 2,2250806384 | 1 | 2,2250806384 | - | - | - | - | - |
| 4CH18 | cHCV | C10-08 | cHCV | F | 60 | 99,156515061 | 22,89603875 | 4,3307279545 | - | - | - | - | - |
| 4CJ23 | cHCV | C10-08 | cHCV | F | 69 | 1,7347462395 | 1 | 1,7347462395 | - | - | - | - | - |
| 4CM44 | cHCV | C10-08 | cHCV | M | 58 | 4,1820445482 | 1 | 4,1820445482 | - | - | - | - | - |
| 4DE28 | cHCV | C10-08 | cHCV | F | 65 | 5,2388824604 | 1 | 5,2388824604 | - | - | - | - | - |
| 4DH16 | cHCV | C10-08 | cHCV | F | 62 | 12,4576565859 | 1 | 12,4576565859 | - | - | - | - | - |
| 4DJ09 | cHCV | C10-08 | cHCV | M | 63 | 1,1207438251 | 10,75379372 | 0,1042184604 | - | - | - | - | - |
| 4EA04 | cHCV | C10-08 | cHCV | M | 58 | 8,1593423324 | 1 | 8,1593423324 | - | - | - | - | - |
| 4EW22 | cHCV | C10-08 | cHCV | M | 44 | 15,034460719 | 1 | 15,034460719 | - | - | - | - | - |
| 4FJ25 | cHCV | C10-08 | cHCV | M | 46 | 14,4373622886 | 1 | 14,4373622886 | - | - | - | - | - |
| 4FJ39 | cHCV | C10-08 | cHCV | M | 58 | 1,6418189148 | 90,7761075 | 0,0180864653 | - | - | - | - | - |
| 4GC32 | cHCV | C10-08 | cHCV | F | 41 | 31,2365639262 | 4,89098805 | 6,386554947 | - | - | - | - | - |
| 4GJ42 | cHCV | C10-08 | cHCV | M | 78 | 2,5197559809 | 49,468848 | 0,0509362171 | - | - | - | - | - |
| 4GM10 | cHCV | C10-08 | cHCV | F | 70 | 9,6211914069 | 1 | 9,6211914069 | - | - | - | - | - |
| 4GR27 | cHCV | C10-08 | cHCV | M | 45 | 5,4132359171 | 1 | 5,4132359171 | - | - | - | - | - |
| 4GS05 | cHCV | C10-08 | cHCV | F | 56 | 1,7687329023 | 1 | 1,7687329023 | - | - | - | - | - |
| 4HC06 | cHCV | C10-08 | cHCV | F | 65 | 9,8294010413 | 1 | 9,8294010413 | - | - | - | - | - |

| | | | | | | | | | | | | | |
|---|---|---|---|---|---|---|---|---|---|---|---|---|---|
| 4HE41 | cHCV | C10-08 | cHCV | na | 65 | 0,5282914168 | 1 | 0,5282914168 | - | - | - | - | - |
| 4LA37 | cHCV | C10-08 | cHCV | M | 58 | 47,9309740429 | 131,648725 | 0,3640823262 | - | - | - | - | - |
| 4LE26 | cHCV | C10-08 | cHCV | M | 44 | 47,8056529908 | 299,6262425 | 0,1595509545 | - | - | - | - | - |
| 4LM21 | cHCV | C10-08 | cHCV | F | 70 | 5,4848474711 | 1 | 5,4848474711 | - | - | - | - | - |
| 4MA01 | cHCV | C10-08 | cHCV | M | 67 | 2,4726104178 | 1024,303475 | 0,0024139432 | - | - | - | - | - |
| 4MF31 | cHCV | C10-08 | cHCV | F | 50 | 2,4553206406 | 1 | 2,4553206406 | - | - | - | - | - |
| 4ML14 | cHCV | C10-08 | cHCV | M | 38 | 8,4613557455 | 1 | 8,4613557455 | - | - | - | - | - |
| 4MP13 | cHCV | C10-08 | cHCV | M | 51 | 3,5539600755 | 1238,046032 | 0,0028706203 | - | - | - | - | - |
| 4NE34 | cHCV | C10-08 | cHCV | F | 61 | 2,5232038272 | 1 | 2,5232038272 | - | - | - | - | - |
| 4NJ02 | cHCV | C10-08 | cHCV | M | 52 | 2,8947902735 | 1 | 2,8947902735 | - | - | - | - | - |
| 4NT33 | cHCV | C10-08 | cHCV | F | 75 | 53,1263837151 | 1 | 53,1263837151 | - | - | - | - | - |
| 4OL24 | cHCV | C10-08 | cHCV | F | 35 | 242,2965635843 | 20,6363625 | 11,7412438158 | - | - | - | - | - |
| 4PG29 | cHCV | C10-08 | cHCV | M | 57 | 13,9742386153 | 1 | 13,9742386153 | - | - | - | - | - |
| 4QT35 | cHCV | C10-08 | cHCV | M | 46 | 5,3217204181 | 3,824914925 | 1,3913304015 | - | - | - | - | - |
| 4RL11 | cHCV | C10-08 | cHCV | F | 49 | 26,1036510193 | 18,12669357 | 1,4400668781 | - | - | - | - | - |
| 4RS43 | cHCV | C10-08 | cHCV | M | 52 | 2,4027262036 | 1 | 2,4027262036 | - | - | - | - | - |
| 4SG07 | cHCV | C10-08 | cHCV | M | 59 | 4,4345162912 | 1 | 4,4345162912 | - | - | - | - | - |
| 4SP36 | cHCV | C10-08 | cHCV | M | 56 | 108,1596453872 | 1 | 108,1596453872 | - | - | - | - | - |
| 4TM45 | cHCV | C10-08 | cHCV | F | 74 | 4,3436003027 | 1 | 4,3436003027 | - | - | - | - | - |
| 4UJ38 | cHCV | C10-08 | cHCV | F | 65 | 530,6916503736 | 214,2097425 | 2,4774393741 | - | - | - | - | - |
| 4VM08 | cHCV | C10-08 | cHCV | M | 70 | 47,6330350763 | 26,69886012 | 1,7840849708 | - | - | - | - | - |
| 4VM20 | cHCV | C10-08 | cHCV | F | 63 | 38,7426438935 | 100,31571863 | 0,3862071111 | - | - | - | - | - |
| KTB079 | Dengue | Upasani *et al.* (2020) | Dengue shock syndrome | M | 9,33 | 626,7159842523 | 959,5281112771 | 0,6531502067 | - | - | - | - | - |
| KTB081 | Dengue | Upasani *et al.* (2020) | Dengue shock syndrome | M | 6,66 | 866,8652783818 | 2277,8521086363 | 0,3805625813 | - | - | - | - | - |
| KTB082 | Dengue | Upasani *et al.* (2020) | Dengue fever | M | 14,033 | 5936,038108849 | 12973,611010204 | 0,4575471011 | - | - | - | - | - |
| KTB085 | Dengue | Upasani *et al.* (2020) | Dengue fever | F | 16,25 | 3808,9191026671 | 10184,6544716247 | 0,3739860899 | - | - | - | - | - |
| KTB086 | Dengue | Upasani *et al.* (2020) | Dengue fever | F | 5,025 | 4638,4731914726 | 5569,2344133413 | 0,8328744756 | - | - | - | - | - |
| KTB087 | Dengue | Upasani *et al.* (2020) | Dengue haemorrhagic fever | M | 11 | 636,8135918796 | 1064,0376668253 | 0,5984878278 | - | - | - | - | - |
| KTB094 | Dengue | Upasani *et al.* (2020) | Dengue fever | M | 11,41 | 2374,1149053221 | 3798,1225972224 | 0,6250759012 | - | - | - | - | - |
| KTB096 | Dengue | Upasani *et al.* (2020) | Dengue fever | M | 12,5 | 2070,9979198556 | 3020,1320693911 | 0,6857309125 | - | - | - | - | - |
| KTB098 | Dengue | Upasani *et al.* (2020) | Dengue fever | F | 6,66 | 3822,0502575205 | 4514,5222554226 | 0,8466123415 | - | - | - | - | - |
| KTB099 | Dengue | Upasani *et al.* (2020) | Dengue haemorrhagic fever | F | 16,41 | 1820,7507829728 | 2912,8870745981 | 0,6250674112 | - | - | - | - | - |
| KTB102 | Dengue | Upasani *et al.* (2020) | Dengue haemorrhagic fever | M | 6,5 | 1063,202558627 | 1913,4843552088 | 0,5556369226 | - | - | - | - | - |
| KTB108 | Dengue | Upasani *et al.* (2020) | Dengue fever | F | 7,5 | 1152,6962345774 | 2060,8413547849 | 0,5593328336 | - | - | - | - | - |
| KTB109 | Dengue | Upasani *et al.* (2020) | Dengue fever | M | 5,75 | 1344,0628483137 | 4012,1513342441 | 0,3349980438 | - | - | - | - | - |
| KTB110 | Dengue | Upasani *et al.* (2020) | Dengue fever | F | 13,5 | 1353,5617055596 | 2538,483586729 | 0,5332166466 | - | - | - | - | - |
| KTB112 | Dengue | Upasani *et al.* (2020) | Dengue haemorrhagic fever | F | 12,25 | 2463,4718529618 | 3287,2417568755 | 0,7494039183 | - | - | - | - | - |
| KTB114 | Dengue | Upasani *et al.* (2020) | Dengue fever | F | 8,66 | 727,8882240122 | 1180,8885329588 | 0,6163902889 | - | - | - | - | - |
| KTB115 | Dengue | Upasani *et al.* (2020) | Dengue fever | M | 11,75 | 78,3753975902 | 103,0934362565 | 0,760236543 | - | - | - | - | - |
| KTB117 | Dengue | Upasani *et al.* (2020) | Dengue fever | F | 9,25 | 3280,0979610189 | 5235,1591671906 | 0,6265517162 | - | - | - | - | - |
| KTB118 | Dengue | Upasani *et al.* (2020) | Dengue fever | M | 9,66 | 17191,4181504615 | 25107,8835656993 | 0,6847020023 | - | - | - | - | - |
| KTB119 | Dengue | Upasani *et al.* (2020) | Dengue shock syndrome | M | 1 | 4060,6558419712 | 28586,1338240401 | 0,1420498437 | - | - | - | - | - |
| KTB120 | Dengue | Upasani *et al.* (2020) | Dengue fever | F | 9 | 1862,2913004316 | 2674,1633354752 | 0,6964014784 | - | - | - | - | - |
| KTB122 | Dengue | Upasani *et al.* (2020) | Dengue fever | F | 13 | 41964,6744950734 | 66745,0609337698 | 0,6287307841 | - | - | - | - | - |
| KTB123 | Dengue | Upasani *et al.* (2020) | Dengue fever | M | 12,33 | 2110,2654722554 | 2825,619674413 | 0,7468328068 | - | - | - | - | - |
| KTB126 | Dengue | Upasani *et al.* (2020) | Dengue fever | F | 14,16 | 35561,9096935754 | 56790,706169437 | 0,6261924194 | - | - | - | - | - |
| KTB129 | Dengue | Upasani *et al.* (2020) | Dengue haemorrhagic fever | F | 7 | 840,4792350508 | 1430,8797847215 | 0,5873863367 | - | - | - | - | - |
| KTB131 | Dengue | Upasani *et al.* (2020) | Dengue fever | M | 12 | 3123,8678731872 | 6977,8880223343 | 0,4476809979 | - | - | - | - | - |
| KTB133 | Dengue | Upasani *et al.* (2020) | Dengue fever | M | 14 | 667,4945465079 | 1682,0946828591 | 0,3968234091 | - | - | - | - | - |
| KTB136 | Dengue | Upasani *et al.* (2020) | Dengue fever | F | 10,41 | 1960,4452381066 | 3505,3987782908 | 0,5592645408 | - | - | - | - | - |
| KTB138 | Dengue | Upasani *et al.* (2020) | Dengue fever | M | 0,83 | 3571,3858174236 | 5149,2736278127 | 0,6935707977 | - | - | - | - | - |
| KTB139 | Dengue | Upasani *et al.* (2020) | Dengue fever | F | 6,41 | 2205,7433004624 | 3456,5642297735 | 0,6381317267 | - | - | - | - | - |
| KTB141 | Dengue | Upasani *et al.* (2020) | Dengue fever | M | 0,41 | 3452,5274236295 | 6022,3813486813 | 0,5732827637 | - | - | - | - | - |
| KTB142 | Dengue | Upasani *et al.* (2020) | Dengue haemorrhagic fever | F | 0,58 | 35263,8425716802 | 51888,7808035094 | 0,6796043774 | - | - | - | - | - |
| KTB146 | Dengue | Upasani *et al.* (2020) | Dengue fever | F | 6 | 1721,1455122814 | 3933,7093419982 | 0,4375375409 | - | - | - | - | - |
| KTB149 | Dengue | Upasani *et al.* (2020) | Dengue fever | F | 4 | 9846,636069503 | 10759,3508279018 | 0,9151700904 | - | - | - | - | - |
| KTB150 | Dengue | Upasani *et al.* (2020) | Dengue fever | M | 7 | 56878,3694048917 | 93976,8473509186 | 0,6052381093 | - | - | - | - | - |
| KTB153 | Dengue | Upasani *et al.* (2020) | Dengue fever | F | 13 | 11308,6413527998 | 16422,4295923259 | 0,6886095196 | - | - | - | - | - |
| KTB156 | Dengue | Upasani *et al.* (2020) | Dengue fever | M | 9,16 | 43229,828579944 | 89350,0805437132 | 0,4838252894 | - | - | - | - | - |
| KTB157 | Dengue | Upasani *et al.* (2020) | Dengue fever | F | 9 | 772,2148108752 | 992,7634249238 | 0,7778437355 | - | - | - | - | - |
| KTB159 | Dengue | Upasani *et al.* (2020) | Dengue haemorrhagic fever | M | 0,75 | 46039,9692890963 | 80848,6202164217 | 0,5694589365 | - | - | - | - | - |
| KTB163 | Dengue | Upasani *et al.* (2020) | Dengue fever | M | 0,66 | 9073,2632926493 | 11356,5222699444 | 0,7989473429 | - | - | - | - | - |
| KTB164 | Dengue | Upasani *et al.* (2020) | Dengue fever | F | 0,66 | 487,1840809885 | 838,2377403258 | 0,5812003654 | - | - | - | - | - |
| KTB172 | Dengue | Upasani *et al.* (2020) | Dengue fever | M | 10,58 | 11163,4112121392 | 12680,9851132518 | 0,8803268131 | - | - | - | - | - |
| KTB179 | Dengue | Upasani *et al.* (2020) | Dengue fever | F | 0,75 | 14461,6703443024 | 24239,9271105491 | 0,596605356 | - | - | - | - | - |
| KTB180 | Dengue | Upasani *et al.* (2020) | Dengue fever | M | 10 | 1164,5384162459 | 4316,6313795408 | 0,2697794446 | - | - | - | - | - |
| KTB187 | Dengue | Upasani *et al.* (2020) | Dengue fever | M | 4 | 1552,8291158956 | 3653,6511736073 | 0,4250074903 | - | - | - | - | - |
| KTB200 | Dengue | Upasani *et al.* (2020) | Dengue fever | F | 14 | 17081,120622456 | 27611,0148170352 | 0,6186342927 | - | - | - | - | - |
| KTB202 | Dengue | Upasani *et al.* (2020) | Dengue fever | F | 11,5 | 1194,4634927579 | 2521,3936102531 | 0,4737314666 | - | - | - | - | - |
| KTB205 | Dengue | Upasani *et al.* (2020) | Dengue fever | M | 10 | 6949,0921613842 | 15864,9397540126 | 0,4380156666 | - | - | - | - | - |
| KTB207 | Dengue | Upasani *et al.* (2020) | Dengue fever | F | 4,75 | 4366,8204216738 | 11033,9696088524 | 0,3957615053 | - | - | - | - | - |

| | | | | | | | | | | | | | |
|---|---|---|---|---|---|---|---|---|---|---|---|---|---|
| KTB210 | Dengue | Upasani *et al.* (2020) | Dengue fever | M | 10,66 | 57197,3522731472 | 86954,757244821 | 0,6577828987 | - | - | - | - | - |
| KTB212 | Dengue | Upasani *et al.* (2020) | Dengue fever | M | 12,33 | 48529,1770096211 | 131208,752729303 | 0,3698623453 | - | - | - | - | - |
| KTB215 | Dengue | Upasani *et al.* (2020) | Dengue fever | M | 10 | 65275,9344281327 | 57648,419703785 | 1,1323109075 | - | - | - | - | - |
| KTB218 | Dengue | Upasani *et al.* (2020) | Dengue fever | F | 10,83 | 2970,9743348238 | 5035,4074953792 | 0,5900166645 | - | - | - | - | - |
| KTB221 | Dengue | Upasani *et al.* (2020) | Dengue fever | F | 15,75 | 12409,1304951988 | 23400,2128582769 | 0,5302998981 | - | - | - | - | - |
| KTB222 | Dengue | Upasani *et al.* (2020) | Dengue fever | M | 15 | 6512,297318643 | 6174,4007803103 | 1,0547253977 | - | - | - | - | - |
| KTB225 | Dengue | Upasani *et al.* (2020) | Dengue fever | M | 10 | 6123,1156172778 | 18610,4041313558 | 0,3290157255 | - | - | - | - | - |
| 153-94 | CNS infection | Rodero *et al.* (2017) | CNS infection | na | na | 95577,43651537 | 88796,4039317922 | 1,0763660721 | - | na | - | - | - |
| 154-66 | CNS infection | Rodero *et al.* (2017) | CNS infection | na | na | 11348,9420420639 | 10162,7972966915 | 1,1167143957 | - | na | - | - | - |
| 159-51 | CNS infection | Rodero *et al.* (2017) | CNS infection | na | na | 12406,7884799275 | 10297,8456507761 | 1,2047945658 | - | na | - | - | - |
| 152-86 | CNS infection | Rodero *et al.* (2017) | Enteroviral meningitis | F | <1 | 9507,6979925191 | 12140,8579582001 | 0,7831158247 | - | 25 | - | - | - |
| 153-19 | CNS infection | Rodero *et al.* (2017) | Enteroviral meningitis | M | <1 | 3786,4067300994 | 2410,0511689773 | 1,5710897672 | - | 18 | - | - | - |
| 165-86 | CNS infection | Rodero *et al.* (2017) | Enteroviral meningitis | F | <1 | 11173,3719113757 | 9663,4733817985 | 1,1562480146 | - | 38 | - | - | - |
| 166-84 | CNS infection | Rodero *et al.* (2017) | Enteroviral meningitis | M | <1 | 6432,1513471865 | 9764,5055077123 | 0,6587278119 | - | 38 | - | - | - |
| 154-78 | CNS infection | Rodero *et al.* (2017) | Herpes simplex encephalitis | F | 36 | 19033,0057199175 | 20960,7770262418 | 0,9080295876 | - | 75 | - | - | - |
| 165-81 | CNS infection | Rodero *et al.* (2017) | Herpes simplex encephalitis | F | 46 | 2737,5638337236 | 4719,6332785809 | 0,5800374038 | - | 9 | - | - | - |
| 160-47 | CNS infection | Rodero *et al.* (2017) | Herpes zoster encephalitis | M | 76 | 2437,4305756901 | 2961,7353894921 | 0,8229737823 | - | 6 | - | - | - |
| 163-52 | CNS infection | Rodero *et al.* (2017) | Herpes zoster encephalitis | F | 90 | 1000,940463222 | 1218,0026860616 | 0,8217883874 | - | 3 | - | - | - |
| 160-92 | CNS infection | Rodero *et al.* (2017) | Herpes zoster meningitis | F | 26 | 2870,7227744097 | 4226,5523715192 | 0,6792114523 | - | 18 | - | - | - |
| 159-75 | CNS infection | Rodero *et al.* (2017) | Post-infectious encephalitis | M | <1 | 8581,599916561 | 3189,2110736555 | 2,6908221872 | - | 9 | - | - | - |
| 162-12 | CNS infection | Rodero *et al.* (2017) | Post-infectious encephalitis | M | 2 | 1711,0915820177 | 3049,1251820331 | 0,5611745927 | - | 6 | - | - | - |
| 165-42 | CNS infection | Rodero *et al.* (2017) | Post-infectious encephalitis | M | <1 | 4174,1615637777 | 2558,3163991052 | 1,6316048966 | - | 6 | - | - | - |
| 158-39 | CNS infection | Rodero *et al.* (2017) | Post-infectious encephalomyelitis | M | 14 | 11801,3686797387 | 3413,0233806693 | 3,457746216 | - | 12 | - | - | - |
| 152-100 | CNS infection | Rodero *et al.* (2017) | Viral meningitis | M | 5 | 5666,8310309994 | 15180,366727978 | 0,3733000087 | - | 18 | - | - | - |
| 159-3 | CNS infection | Rodero *et al.* (2017) | Viral meningoencephalitis | M | <1 | 22201,6053315561 | 2723,0728705329 | 8,1531440351 | - | >200 | - | - | - |
